# Effects of Nanopore Confinement on the Conformational, Dynamical, and Self-Assembly Properties of an FG-Repeat Peptide

**DOI:** 10.1101/2025.09.11.675621

**Authors:** Wancheng Zhao, Wai-Ming Yau, Robert Tycko

## Abstract

The central channels of nuclear pore complexes (NPCs) in eukaryotic cells are filled with protein chains whose sequences contain characteristic phenylalanine-glycine motifs. Properties of these nanopore-confined FG-repeat sequences are of central importance to NPC function. Here we demonstrate an approach to nuclear magnetic resonance (NMR) studies of FG-repeat sequences (or other polypeptides) that are tethered within pores with diameters similar to those of NPC channels. By attaching alkyl phosphonate groups to the N-terminus of a 30-residue peptide that contains four FG repeats, called FG30, we tether FG30 chains to walls of 20-nm-diameter pores in anodic aluminum oxide (AAO) wafers through phosphonate-surface bonds. Quantitative ^13^C and ^31^P NMR measurements indicate 90 mM peptide concentrations (300 mg/ml) within the pores. NMR spectra and spin relaxation measurements show that FG30 chains are dynamically disordered and random-coil-like in buffer-filled pores over a broad temperature range. In contrast, FG30 aggregates in free solution at concentrations above 2 mM, forming structurally ordered fibrils according to electron microscopy and NMR measurements. These results demonstrate the utility of AAO as a scaffold for studies of polypeptides in nanopore-confined environments and show that tethering to nanopore walls can dramatically alter the self-assembly properties of an FG-repeat sequence.

**TOC Graphic:** 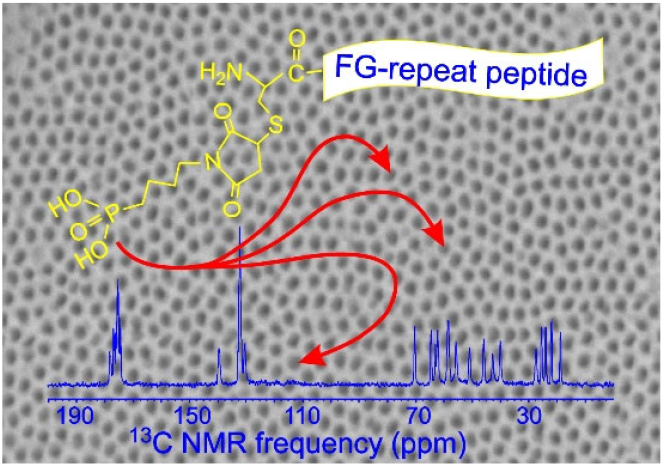

## INTRODUCTION

The behavior of polypeptide chains that are tethered at high densities to the inner surfaces of nanopores is biologically important in the context of the nuclear pore complexes (NPCs) of eukaryotic cells, which restrict and control the passage of proteins, RNA, and macromolecular complexes between the nucleus and the cytoplasm.^1–3^ The central channel of a NPC has a diameter in the 40-80 nm range, depending on the organism and the state of the cell.^4–7^ Protein segments known as FG-repeat domains, due to the presence of multiple phenylalanine-glycine pairs spaced 10-15 residues apart in their amino acid sequences, project into the central channel. The density of FG-repeat domains within the central channel is approximately 100-200 mg/ml,^8^ similar to the typical protein density in a liquid-liquid phase-separated protein droplet.^9–10^ The high density of FG-repeat domains prevents macromolecules and complexes with molecular weights greater than roughly 60 kDa from passing freely through the central channel,^11^ while diffusion of smaller entities remains relatively rapid due to the flexibility of and spaces between FG-repeat domains. Rapid passage of large macromolecules and complexes depends on their binding to nuclear transport receptors (NTRs) and on interactions of NTRs with FG motifs.^12–14^

Molecular details of the mechanisms by which FG-repeat domains control passage of macromolecules through NPC channels are not fully understood,^13,15–16^ motivating the development of NPC-mimetic approaches for biophysical studies.^17^ Hydrogel particles^18–20^ and droplets^21^ formed by isolated FG-repeat domains have been shown to exhibit size and sequence selectivity and NTR dependence similar to that of true NPC channels, but are larger than NPCs and lack a pore geometry. NPC-like pores containing internally tethered FG-repeat sequences have been created by DNA origami methods.^22–24^ Solid-state nanopores to which FG-repeat sequences can be attached have also been prepared by lithographic methods.^8,25–27^

In principle, nuclear magnetic resonance (NMR) measurements can provide a wealth of information about the conformational distributions, molecular motions, and interactions of FG-repeat domains. NMR studies of FG-repeat domains by several groups have explored properties of their monomeric states,^28–30^ interactions with NTRs in solution,^29–30^ molecular motions in hydrogel particles,^31^ and intermolecular interactions in FG-repeat hydrogels.^19,32^ None of these studies involved an NPC-mimetic nanopore environment. Unfortunately, the quantities of FG-repeat sequences in NPC-mimetic DNA origami constructs and solid-state nanopores are grossly insufficient for NMR studies.

This paper introduces a new method for creating samples in which FG-repeat sequences (or other polypeptides) are tethered to the walls of nanopores, with adequate quantities for detailed NMR studies. The method takes advantage of the reactivity of phosphonate groups with aluminum oxide surfaces^33–35^ and the commercial availability of nanoporous anodic aluminum oxide (AAO) wafers that contain a high density of long pores with uniform diameters.^36–37^ As depicted in Fig. 1, we report results from experiments on a 30-residue peptide from human Nup98, containing four FG motifs, which we call FG30. Nup98 was chosen due to its high density of FG motifs, its importance for NPC function, and its use in previous studies.^16,19–20,28,38–39^ We show that FG30 can be tagged with an N-terminal phosphonate linker through cysteine-maleimide chemistry and covalently attached to surfaces within 20-nm-diameter AAO pores, achieving peptide concentrations within the pores up to 90 mM (300 mg/ml).

**Figure 1.**
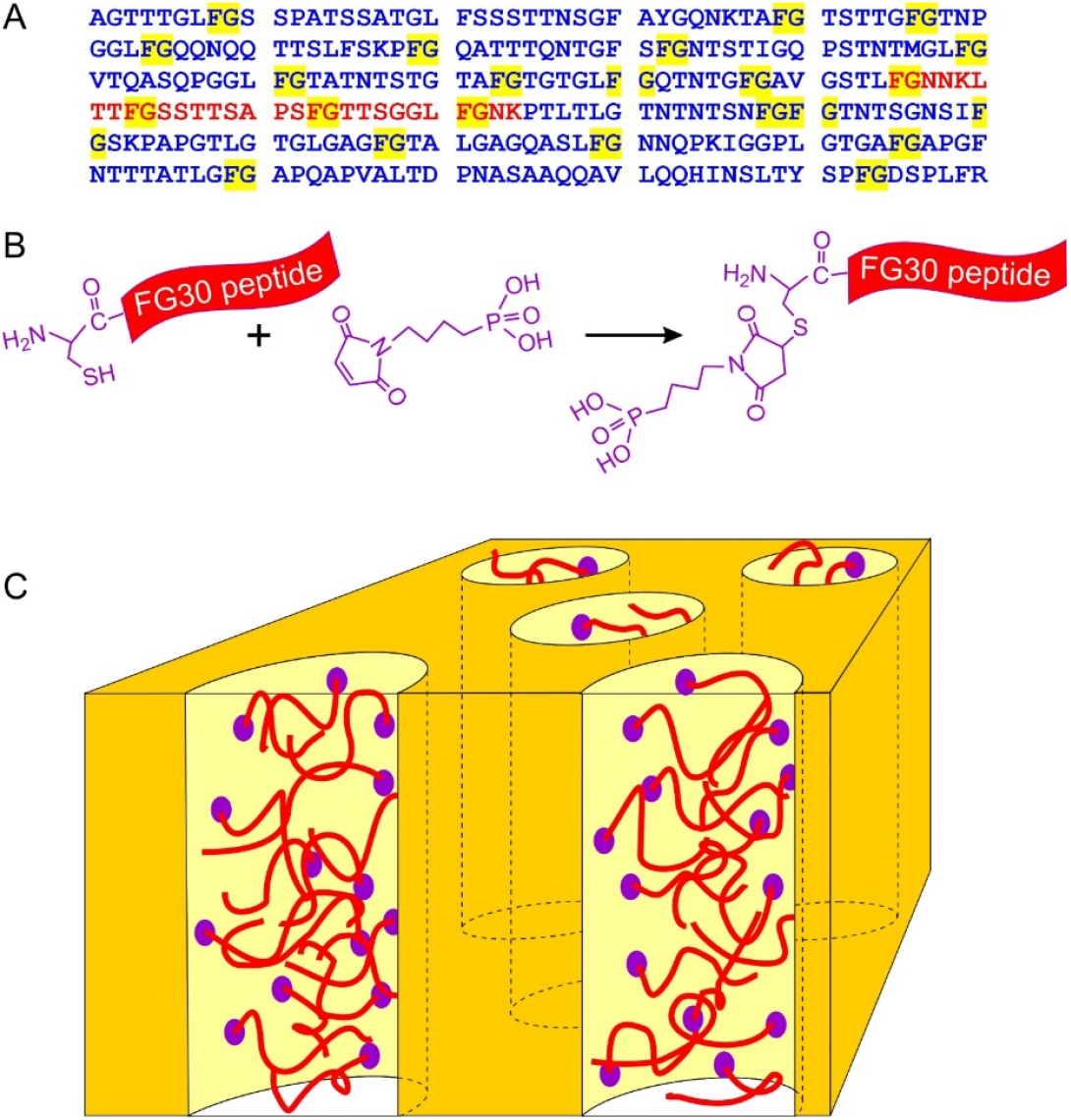
(A) Sequence of the second FG-repeat domain of human Nup98 (Uniprot code P52948, residues 221-520). FG dyads are highlighted in yellow. The FG30 segment studied in this work is shown in red. (B) Method for introducing a phosphonate group by conjugating 4-maleimidobutylphosphonic acid to an N-terminal cysteine to produce MBPA-C-FG30. (C) Cartoon representation of peptides tethered to the walls of nanopores in AAO by reaction of phosphonate groups with aluminum oxide surfaces.

NMR measurements described below reveal that FG30 peptides are dynamically disordered at high concentrations in fully hydrated, 20-nm-diameter AAO pores over a wide temperature range, with no signs of aggregation over many weeks. In contrast, the same peptide readily aggregates to form fibrils in free solution at concentrations above 2 mM (7 mg/ml). Thus, tethering and confinement within nanopores can strongly affect the properties of an FG-repeat sequence. Since full-length FG-repeat domains of nuclear pore proteins (Nups) are also known to be aggregation-prone in free solution,^28,32,40–42^ in analogy to the fibril-formation propensities of other low-complexity sequences,^43–45^ our results for nanopore-confined FG30 may have implications for the properties of FG-repeat domains in NPC channels.

## MATERIALS AND METHODS

### Preparation of FG30

C-FG30 peptides (sequence C-FGNNKLTTFGSSTTSAPSFGTTSGGLFGNK, with C-terminal amidation) were synthesized at a 0.1 mmol scale on a Biotage Initiator+ Alstra solid-phase peptide synthesizer, using standard Fmoc chemistry and activation with diisopropylcarbodiimide and OxymaPure in N-methyl-2-pyrrolidone. A rink amide ChemMatrix resin (Sigma-Aldrich) was used, with a substitution of 0.48 mEq/g as determined from the ultraviolet absorbance of the dibenzofulvene-piperidine adduct.^46^ Peptides were synthesized with uniform ^15^N,^13^C-labeled residues at L6, S11, A16, F19, T22 and G28 (called C-FG30-LSAFTG) or G2, G10, G20, and G29 (called C-FG30-4G). Unlabeled Fmoc-amino acids were coupled with a five-fold excess for 15 min at 75° C. Fmoc-Asn(Trt)-OH and Fmoc-Pro-OH were double-coupled, using 15 min at 75° C for each coupling. Isotopically labeled Fmoc-amino acids were coupled with a three-fold excess for 15 min at 75° C, followed by coupling of a five-fold excess of the corresponding unlabeled Fmoc-amino acids for 5 min at 75° C. Deprotection steps used 20% piperidine with 0.1M 1-Hydroxybenzotriazole hydrate in dimethyl formamide for 5 min at 75° C. Peptides were cleaved from the resin with a standard cocktail (8.75 ml trifluoroacetic acid, 0.75 g phenol, 0.5 ml thioanisole, 0.25 ml ethane-1,2-dithiol, 0.5 ml H_2_O) for 2 h. After filtration and precipitation with cold t-butyl methyl ether (TBME), the crude product was further washed twice with TBME. Peptides were purified by reverse-phase high-performance liquid chromatography (HPLC) on a Proto 300 C18 column (Higgins Analytical).

To prepare 4-maleimidobutylphosphonic acid (MBPA), 4-aminobutylphosphonic acid (1 mmol, ChemScene) and maleic anhydride (1 mmol, Sigma-Aldrich) were suspended in 10 ml glacial acetic acid (Macron) and stirred overnight at room temperature. After the reaction was refluxed for an additional 5 h, n-heptane (25-30 ml, Alfa Aesar) was added to the reaction and glacial acetic acid was distilled off as an azeotrope. Addition of n-heptane and distillation were repeated three times. The desired product, 4-maleimidobutylphosphonic acid, was collected as an oil, dried under vacuum overnight, and used without further purification.

To conjugate MBPA to the N-terminal cysteine residue of C-FG30, C-FG30 (3 mmol) and MBPA (10 mmol) were dissolved in 2.0 ml of dimethyl formamide/water (1:1 by volume) and stirred at room temperature. Completion of the conjugation reaction that was monitored by liquid chromatography-mass spectrometry (LC-MS; Thermo Scientific UltiMate 3000 and ISQEM-ESI single quadrupole mass spectrometer). The final product was purified by HPLC on a Proto 300 C18 reverse-phase column (Higgins Analytical).

Melittin with an additional N-terminal cysteine (sequence C-GIGAVLKVLTTGLPALISWIKRKRQQ, with C-terminal amidation) and with uniform ^15^N,^13^C-labeled residues at G3, V8, and A15 was synthesized and purified with the same methods. MBPA was conjugated to the N-terminal cysteine to produce MBPA-C-melittin-GVA. Fig. S1 shows LC-MS results for MBPA-C-FG30-LSAFTG, MBPA-C-FG30-4G, and MBPA-C-melittin-GVA.

For confocal fluorescence imaging, MBPA-C-FG30 was labeled with fluorescein by reaction of fluorescein isothiocyanate (FITC) with the peptide amine groups. Approximately 2 mg of MBPA-C-FG30-4G was dissolved in 1 ml of sodium carbonate buffer (0.1 M, pH 9.0), and 1 mg of FITC was dissolved in anhydrous dimethyl sulfoxide (DMSO). The FITC solution was slowly added to the peptide solution in 50 µl aliquots, with gentle stirring. The reaction mixture was incubated in the dark at 4 °C for approximately 16 h. After confirming product formation by LC-MS, the reaction was quenched by addition of excess ammonium chloride, with a final concentration of 50 mM, and an additional incubation at 4 °C for 1 h. The final mixture was purified by preparative HPLC on the C18 reverse-phase column and subsequently lyophilized. LC-MS measurements indicated a single fluorescein per peptide molecule.

Aggregated MBPA-C-FG30 for transmission electron microscopy (TEM) was prepared by dissolving 2.0 mg of purified and lyophilized MBPA-C-FG30-LSAFTG in 60 µl of 0.2 M Bis-Tris buffer (pH 6.5) to produce a 10 mM peptide solution. The solution was evenly divided into three 20 µl aliquots, each transferred into a 0.2 ml tube. The three tubes were incubated quiescently for 6 days at temperatures of 4° C, 24° C, and 37° C. After incubation, each sample was briefly vortexed, and 3 µl was withdrawn for subsequent TEM imaging, as described below.

For NMR measurements on aggregated MBPA-C-FG30, a second 60 µl volume of 10 mM MBPA-C-FG30-LSAFTG peptide solution was prepared in the same way, then incubated quiescently at 24° C for 13 days. Aggregated material was pelleted at 278,000 × g at 20° C for 120 min, using a Beckman Coulter Optima MAX ultracentrifuge with a TLA100.1 rotor. Approximately 3 mg of wet pellet, including residual buffer, was then transferred into a 3.2 mm magic-angle spinning (MAS) rotor by centrifugation at 71,000 × g and 4° C for 1 h using a Beckman Coulter Optima XL-100K Ultracentrifuge with an SW40Ti rotor. The aggregated sample remained fully hydrated throughout the subsequent NMR measurements.

### NMR measurements

NMR measurements on MBPA-C-FG30 samples were performed at 14.1 T (599.1 MHz, 242.5 MHz, and 150.7 MHz frequencies for ^1^H, ^31^P, and ^13^C, respectively) using a Tecmag Redstone spectrometer and a three-channel magic-angle spinning (MAS) NMR probe from Black Fox LLC. NMR measurements on MBPA-C-melittin-GVA were performed at 17.5 T (744.6 MHz, 301.4 MHz, 187.2 MHz for ^1^H, ^31^P, and ^13^C, respectively), also using a Tecmag Redstone spectrometer and a three-channel MAS NMR probe from Black Fox LLC. MAS frequencies were 12.00 kHz and 12.50 kHz in measurements on MBPA-C-FG30 and MBPA-C-melittin, respectively, unless otherwise stated. Sample temperatures were controlled with nitrogen gas from an FTS AirJet XR cooler and calibrated with measurements on KBr powder^47^ under the same gas flow and MAS conditions.

Radio-frequency (rf) pulse sequences for all NMR measurements on MBPA-C-FG30 are shown in Fig. S2, with information about pulse sequence parameters in the figure caption. One-dimensional (1D) ^13^C and ^31^P NMR spectra were recorded with either cross-polarization^48^ (CP) from ^1^H nuclei to ^13^C or ^31^P nuclei (Fig. S2A,B) or direct pulsing (DP) of ^13^C or ^31^P nuclei (Fig. S2C). Typically, the ^1^H rf amplitude was 66 kHz and the ^13^C or ^31^P rf amplitude was ramped linearly from 49 to 59 kHz during the CP contact time. The CP echo sequence in Fig. S2B was used in some cases to minimize baseline distortions in 1D spectra from rf pulse ring-down. 1D ^13^C NMR spectra were also recorded with the “insensitive nuclei enhanced by polarization transfer” (INEPT) technique^49^ (Fig. S2D), using a total polarization transfer period of 4.8 ms. ^1^H decoupling amplitudes were typically 88 kHz in CP and DP measurements, with two-pulse phase modulation^50^ (TPPM) during signal acquisition periods. Decoupling amplitudes were 10 kHz in INEPT measurements, with composite pulse decoupling^51^ based on a 125-pulse composite π sequence.^52^ For quantitative 1D ^13^C and ^31^P spectra, recycle delays were adjusted based on measurements of ^1^H, ^13^C, ^31^P spin-lattice relaxation times. Recycle delays were 4-8 s for ^13^C CP spectra, 2-25 s for ^13^C DP spectra, 2-12 s for ^31^P CP spectra, and 7-1000 s for ^31^P DP spectra.

Two-dimensional (2D) ^13^C-^13^C NMR spectra of MBPA-C-FG30 samples that show signals from sites with restricted mobility were recorded with ^1^H-^13^C CP and a 30 ms dipolar-assisted rotational resonance^53^ (DARR) mixing period for ^13^C-^13^C polarization transfers (Fig. S2E). For measurements at 14.1 T, the recycle delay was 1.0 s, 128 complex t_1_ points were acquired with a 25.0 μs t_1_ increment, and 256 or 512 scans were taken per complex t_1_ point. For measurements at 17.5 T, the recycle delay was 2.0 s, 160 complex t_1_ points were acquired with a 20.0 μs t_1_ increment, and 768 scans were taken per complex t_1_ point.

2D ^1^H-^13^C NMR spectra that show signals from highly mobile sites were acquired with ^1^H-^13^C INEPT (Fig. S2F), a 1.0 s recycle delay, 64 complex t_1_ points with a 100.0 μs t_1_ increment, and 512 scans per complex t_1_ point. 2D ^13^C-^13^C NMR spectra that show signals from highly mobile sites were acquired with ^1^H-^13^C INEPT and with ^13^C-^13^C polarization transfers driven by scalar couplings in a 12.5 ms “total correlation spectroscopy” (TOCSY) mixing period^54^ (Fig. S2G). During the mixing period, ^13^C chemical shift differences were removed by applying a 125-pulse composite pulse sequence^52^ twice, with a 10 kHz ^13^C rf amplitude. The recycle delay was 1.0 s, 64 complex t_1_ points were acquired with a 100.0 μs t_1_ increment, and 512-864 scans were taken per complex t_1_ point.

Site-specific ^1^H and ^13^C spin relaxation times were measured from the dependence of resolved ^13^C NMR signals on relaxation periods t_rel_. ^1^H spin-lattice relaxation times T_1_^H^, rotating-frame relaxation times T_1ρ_^H^, and transverse relaxation times T_2_^H^ were measured by inversion-recovery during t_rel_, spin-locking decay along a 10 kHz field during t_rel_, or spin-echo decay during t_rel_, respectively, followed by INEPT transfers to ^13^C nuclei for signal detection (Figs. S2H-S2J).

^13^C spin-lattice relaxation times T_1_^C^ were measured by inversion-recovery during t_rel_, followed by spin-echo detection (Fig. S2K). ^13^C transverse relaxation times T_2_^C^ were measured by spin-echo decay during t_rel_. To avoid spin-echo modulation due to ^13^C-^13^C scalar couplings, a Gaussian-shaped, frequency-selective π pulse (2.35 ms pulse length) was used in T_2_ ^C^ measurements, with a ^13^C carrier frequency offset Δf that was adjusted to select signals from individual sites or groups of sites (Fig. S2L). Recycle delays were 2.0 s in all relaxation measurements.

^13^C chemical shifts are relative to 4,4-dimethyl-4-silapentane-1-sulfonic acid, using the ^13^C_α_ line of L-valine powder at 63.46 ppm as an external standard. ^31^P chemical shifts are relative to phosphoric acid, using the ^31^P line of aminomethyl phosphonic acid (AMPA) at 18.8 ppm as an external standard.^55^ 1D spectra were processed, analyzed, and plotted with Tecmag TNMR software. 2D spectra were processed and plotted with nmrPipe software.^56^

Nuclear spin relaxation data were analyzed with GraphPad Prism 10. In most cases, signal amplitudes *A*(*t*_*rel*_) were measured from peak heights in 1D ^13^C NMR spectra and uncertainties were set equal to the root-mean-squared (RMS) noise in the spectra. For relaxation measurements on unresolved backbone CO signals and unresolved signals from F19 aromatic sites C2-C6, signal amplitudes were measured as peak areas and uncertainties were set equal to the RMS values of noise areas, using the same frequency ranges to calculate all areas. In all cases, the build-up or decay of signal amplitudes with increasing t_rel_ was fit initially with a single-exponential function, *e*.*g*., *A*(*t*_*rel*_) = *a* + *b* exp(-*t* / *T*_1_ ^*H*^), with *a* ≠ 0 for T_1_^H^ and T_1_^C^ data only. We considered the resulting single-exponential fit to be adequate if the reduced χ^2^ value reported by the GraphPad software was less than 2.0. Single-exponential fits were not adequate (χ^2^ > 2.0) for some of the T_1 ρ_^H^, T_2_^H^, and T_2_^C^ data. In those cases, we used a stretched-exponential function to determine the relaxation time, *e*.*g*., *A*(*t*_*rel*_) = *b* exp[-(*t* / *T*_2_ ^*H*^)^*β*^], with β being the stretching parameter. If a stretched-exponential function was required to fit a given relaxation measurement for a given site at any temperature, then stretched-exponential functions were used at all temperatures.

### Analyses of nuclear spin relaxation data

A subset of the relaxation data was chosen for analysis by comparison with expressions for relaxation rates for dipole-coupled two-spin systems (see Eqs. (1-3) of the Results), with the assumption that molecular orientational motions result in correlation functions of the form *C*(*t*) = *C*(0)[*a*_1_ exp(−*t τ*_*c*1_) + (1− *a*_1_) exp(−*t τ*_*c* 2_)]. Analyses were performed by evaluating chi-squared deviations 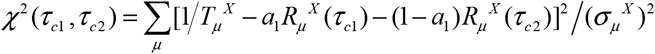 between experimental and calculated relaxation rates over a grid of 100 τ_c1_ and 100 τ_c2_ values, with 10 ns ≤ τ_c1_ ≤ 500 ns and 0.1 ns ≤ τ_c2_ ≤ 9 ns and with constant increments of log(τ_c1_) and log(τ_c2_). In this expression for *χ*^2^ (*τ*_c1_,*τ*_c2_), μ = 1 and 2 for X = C and μ = 1, 1ρ, and 2 for X = H, and σ_μ_^X^ is the uncertainty in the experimental value of 1/T_μ_^X^. For each choice of τ_c1_ and τ_c2_, a_1_ was determined analytically to minimize *χ*^2^ (*τ*_*c*1_,*τ*_*c*2_).

Since acceptably small values of *χ*^2^ (*τ*_*c*1_,*τ*_*c* 2_) were obtained over broad ranges of the fitting parameters, we calculated probability-weighted averages and standard deviations of the parameters, rather than simply reporting the best-fit values of the parameters. The probability-weighted average of τ_c1_ was defined as 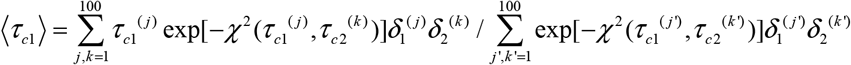, where the sums are over grid points, with δ_1_^(j)^ and δ_2_^(k)^ being the grid spacings in τ_c1_ and τ_c2_ dimensions, respectively. The standard deviation in τ_c1_ was defined as 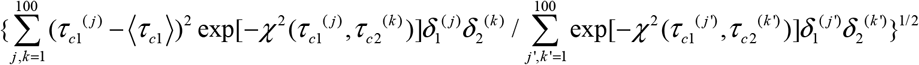.

Analogous definitions were used for probability-weighted averages and standard deviations of τ_c2_ and a_1_.

Simulations of ^1^H spin-echo decay in Fig. S22 were performed with a home-built program that was adapted specifically for this purpose (available upon request). The program calculates the quantum mechanical evolution of a dipole-coupled spin system under MAS for multiples of the MAS rotation period, with an ideal π pulse in the middle of the evolution period, starting with a density matrix proportional to the x component of total spin angular momentum. The complex NMR signal is evaluated at the end of the evolution period. Isotropic chemical shifts and powder averaging are included.

### Electron microscopy

Transmission electron microscope (TEM) images were acquired with an FEI Morgagni microscope, operating at 80 kV, and a side-mounted Advantage HR camera (Advanced Microscopy Techniques). TEM grids were lacey carbon on 300 mesh copper (Electron Microscopy Sciences), upon which carbon films were deposited by flotation. Solutions containing aggregated MBPA-C-FG30 were diluted 100-fold in H_2_O, then adsorbed in 10 μl aliquots to glow-discharged grids for 90 s. Grids were blotted, rinsed with H_2_O, blotted, rinsed again, blotted, and stained with 2% w/v uranyl acetate for 15 s before final blotting and drying in air.

Scanning electron microscope (SEM) images of AAO20 wafers were acquired with a Hitachi S-4800 field emission SEM instrument. Fragments of wafers were mounted on carbon-based conductive adhesive tape and further immobilized using ultraviolet-curable adhesive (Loctite AA 3494) to minimize drift during imaging, then coated with 1 nm of gold by argon plasma sputtering. The accelerating voltage was set to 1.0 keV. The detection mode in the Hitachi software was set to “L.A. 1” for a combination of secondary electron and low-angle back-scattered electron detection.

### Atomic force microscopy

Atomic force microscope (AFM) images were acquired in tapping mode with a Veeco Multimode instrument with Nanoscope IV controller, using a Nanosensors PPP-NCHAuD probe with 320 kHz resonant frequency. Images were recorded at 7.96 μm/s tip velocities with 512 × 512 points and 1.0 μm × 1.0 μm nominal dimensions. Image dimensions were corrected by measurements on a standard calibration grid.

### Optical and fluorescence microscopy

For confocal fluorescence microscopy, FITC-labeled MBPA-C-FG30-4G (prepared as described above) was dissolved in Bis-Tris buffer (0.2 M, pH 6.5) to obtain a 1.0 mM solution. This solution was then diluted 40-fold with a 1.0 mM solution of unlabeled MBPA-C-FG30-4G in the same buffer to reduce the FITC density. H_2_SO_4_-treated AAO20 wafer samples (~3 mg each) were incubated in 200 µl of the diluted solution in 0.5 ml tubes at 40° C with rotation and shaking. Four wafers were processed in parallel, each in a separate tube, with incubation times of 10 min, 1 h, 3 h, and 24 h. After incubation, wafer samples were washed in DMSO for 10 minutes, followed by three 10-minute washes with deionized water.

For control experiments, wafer pieces were incubated quiescently at 24° C for 10 min and 30 min in solutions containing 50 μg/ml fluorescein and 6 M guanidine hydrochloride (GuHCl), followed by rinsing with water. GuHCl was included to reduce adsorption of fluorescein to AAO20 surfaces, which otherwise resulted in high fluorescein densities that interfered with confocal fluorescence imaging. The fluorescein solution was prepared by diluting a 1 mg/ml fluorescein stock solution in DMSO with GuHCl/Bis-Tris buffer to the desired concentration. All prepared wafers were stored in water and kept in the dark until shortly before confocal imaging.

Confocal fluorescence imaging was performed with a Zeiss LSM 780 microscope equipped with a Plan-Apochromat 100x/1.40 Oil DIC objective lens. AAO20 wafer pieces were imaged in water-filled 35 mm glass-bottom microwell dishes (MatTek Corporation). Wafer pieces were covered by 12 mm circular cover glass (Erie Scientific Company), with an additional metal O-ring on top of the cover glass to keep the wafer in position during imaging. Fluorescence was excited with a 405 nm laser and collected with a 514 – 585 nm emission filter. The pinhole was set to 1.31 Airy units and z-stacks were acquired at 1.0 μm intervals. No refractive index corrections were applied. Imaging parameters were kept constant for all samples. To generate plots of fluorescence intensity as a function of z (the direction parallel to AAO20 pores), Fiji software^57^ was used to calculate the average intensity in each slice of the confocal z-stack, averaging over the entire field of view in each slice.

Bright-field optical images of AAO20 wafers were recorded with an Olympus BX50 microscope, equipped with a SPOT Idea camera.

## RESULTS

### Characterization of AAO wafers

AAO wafers with 13 mm diameter, 50 μm thickness, and 20 ± 3 nm diameter pores were obtained from InRedox LLC (part number AAO-013-020-050, subsequently called AAO20). SEM and AFM images of AAO20 wafers as received (Figs. S3A and S3B) show a pseudo-regular pattern of pores with an average density of 467 ± 69 pores/μm^2^ (482 ± 31 pores/μm^2^ from four AFM images with approximately 0.7 μm^2^ area in each image; 435 ± 91 pores/μm^2^ from five SEM images with approximately 1.0 μm^2^ area in each image). SEM images of fractured edges of AAO20 wafers show the expected straight pores, with diameters that are consistent with the expected 20 nm value to within the image resolution (Fig. S3A).

With a 20 nm pore diameter, the average pore density seen in SEM and AFM images implies a porosity value (*i*.*e*., fraction of wafer volume occupied by pores) equal to 14.7 ± 2.2%. Assuming smooth pore walls, a 13 mm wafer diameter, and a 50 μm wafer thickness, the calculated surface area per within pores is (1.95 ± 0.29) × 10^5^ mm^2^ per wafer. In contrast, the area of the exterior surfaces of the wafer is only 230 mm^2^. If molecules could be attached to the pore walls of AAO20 with an average spacing of 1 nm between molecules, these calculations suggest that one wafer could hold approximately 0.3 μmole of molecules, a quantity that is easily sufficient for NMR measurements.

We also estimated the porosity of AAO20 wafers by weighing dry wafers, soaking them in H_2_O, quickly blotting them with filter paper, and weighing them again within 5 s of removal from H_2_O. These measurements were performed three times on each of three AAO20 wafers. Assuming that mass differences between dry and hydrated states were due to H_2_O that completely filled the pores in wafers with 13 mm diameter and 50 μm thickness (6.64 μl volume), we obtained a porosity value of 16.3 ± 1.0%, in good agreement with the value estimated from SEM and AFM images.

^31^P NMR spectra of untreated AAO20 revealed the unanticipated presence of a relatively large signal (Fig. S4A). The directly-pulsed ^31^P signal area from untreated AAO20, compared with the ^31^P signal area from a known mass of ammonium phosphate, implied approximately 1.2 μmol of phosphorus per AAO20 wafer, which would correspond to a surface density of approximately 4 nm^−2^ if the phosphorus were localized on surfaces. Following a suggestion from Dr. Dmitri Routkevitch of InRedox LLC, we found that brief treatments with 0.1 M H_2_SO_4_ can be used to remove residual surface phosphates (Fig. S4B) without increasing the AAO pore diameters significantly (Figs. S3C and S3D).

Finally, we examined the effects on AAO20 of incubation at 40° C in aqueous buffers with pH values between 0.4 and 14.7. After 94 h incubation periods at pH values below 6.2 or above 8.1, AAO20 wafer samples were obviously damaged, dissolving or losing their normal transparency (Fig. S5). Consequently, incubations of AAO20 with MBPA-C-FG30 were performed with 0.2 M Bis-Tris buffer, pH 6.5 (except as noted). SEM images (Fig. S3C) confirmed that the nanoporous structure of AAO20 was retained after brief treatments with 0.1 M H_2_SO_4_ and incubation in 0.2 M Bis-Tris buffer, pH 6.5, at 40° C for 24 h. Additionally, NMR results described below show that MBPA-C-FG30/AAO20 samples remain stable in 0.2 M Bis-Tris buffer, pH 6.5, during many weeks of measurements over a wide temperature range.

### Attachment and quantification of FG30 in AAO nanopores

As described in the Materials and Methods, FG30 peptides were synthesized with additional N-terminal cysteine residues, then ligated with MBPA through a maleimide-thiol reaction to produce MBPA-C-FG30 (Figs. 1A and 1B). In one synthesis, ^15^N,^13^C-labeled amino acids were introduced at L6, S11, A16, F19, T22, and G28 (MBPA-C-FG30-LSAFTG), chosen to maximize resolution in NMR spectra and allow probing of sites along the FG30 chain. In another synthesis, labels were introduced at G2, G10, G20, and G29 (MBPA-C-FG30-4G) to produce simple ^13^C NMR spectra with maximal signal-to-noise. 1D and 2D NMR spectra of lyophilized MBPA-C-FG30-LSAFTG (Fig. S6) show inhomogeneously broadened ^13^C and ^31^P NMR lines, as expected for a conformationally disordered and rigid state.

Attachment of MBPA-C-FG30 to pore surfaces in AAO20 was performed by incubating wafers or wafer pieces at 40° C in 1.0 mM solutions of MBPA-C-FG30 in 0.2 M Bis-Tris buffer, pH 6.5. Typically, 1.0 ml of solution was contained in a 1.5 ml tube, with end-over-end rotation at 30 revolutions per minute and intermittent shaking in each rotation cycle (Roto-Therm Mini Plus, Benchmark Scientific). After incubation, the AAO20 material was rinsed with three successive 1.5 ml volumes of deionized water, then dried overnight under a stream of nitrogen gas. The dried material was crushed into small pieces, approximately 0.5-2 mm in diameter, and packed into MAS rotors for NMR measurements.

Figs. 2A and 2B show 1D ^13^C and ^31^P NMR spectra of dry samples of MBPA-C-FG30-LSAFTG/AAO20 and MBPA-C-FG30-4G/AAO20. From saturation-recovery measurements, spin-lattice relaxation times for ^1^H, ^13^C, and ^31^P nuclei in these dry samples (T_1_^H^, T_1_^C^, T_1_^P^) were determined to be approximately 0.6 s, 1.5 s (LSAFTG) or 30 s (4G), and 200 s, respectively. Similar spectra were obtained for samples prepared with a variety of incubation conditions (Figs. S7 and S8), using recycle delays greater than three times the relevant T_1_^H^ or T_1_^C^ values. Aliphatic ^13^C and ^31^P signal areas in these spectra were measured, normalized by the number of scans in each spectrum, and compared with signal areas in ^13^C NMR spectra of 1.25 mg of uniformly ^15^N,^13^C-labeled L-valine powder (10.1 μmol of L-valine, 40.6 μmol of aliphatic ^13^C) and 5.0 mg of aminomethyl phosphonic acid (AMPA) powder (45.0 μmol of ^31^P) (Fig. S9). The resulting estimates of the loading of MBPA-C-FG30 in AAO20 are listed in Table S1. In most cases, estimates of loading from ^13^C CP, ^13^C DP, and ^31^P CP signal areas are in good agreement for each sample, with discrepancies attributable to limited signal-to-noise ratios and uncertainties regarding baseline corrections in the spectra.

**Figure 2.**
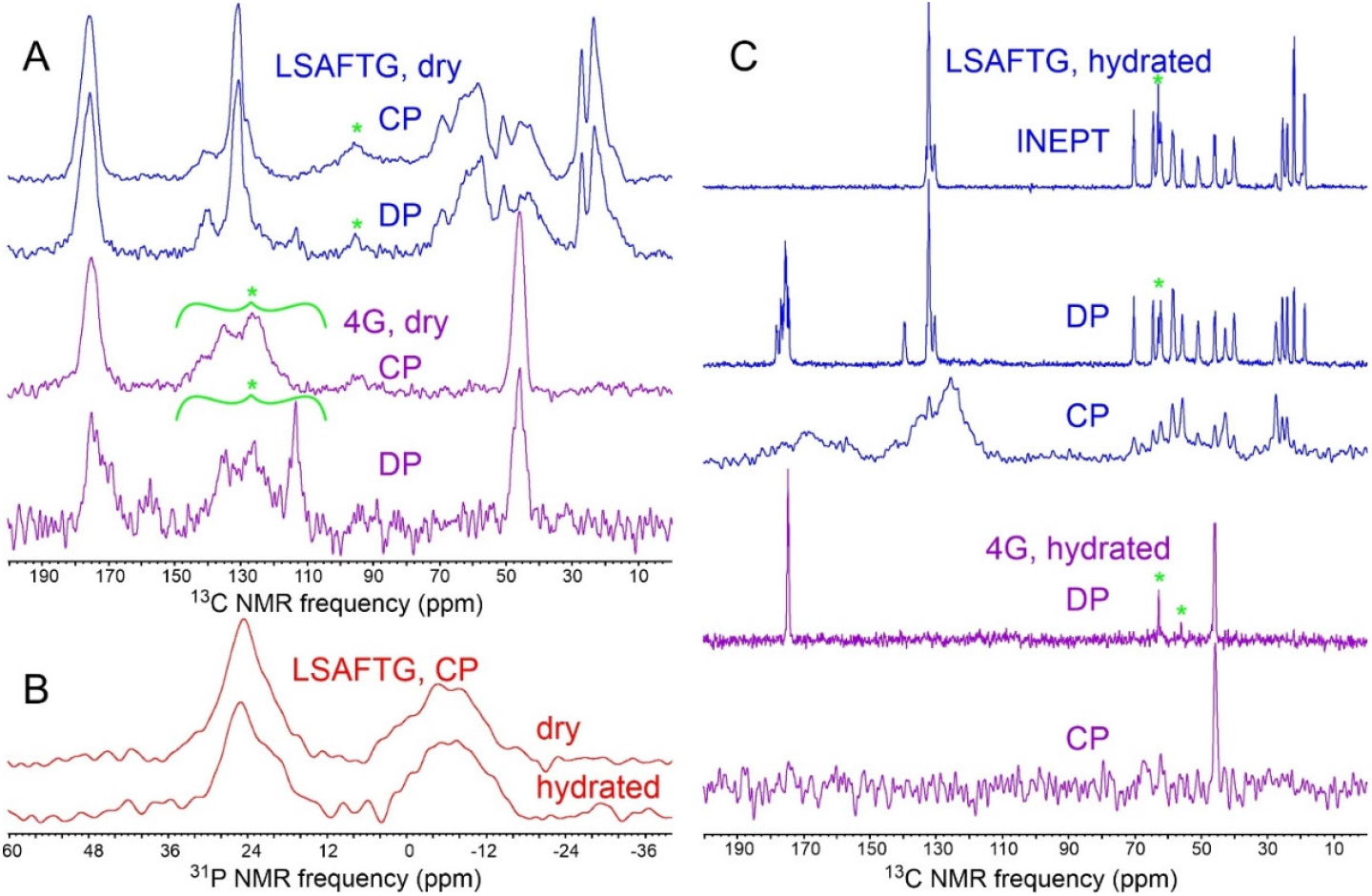
(A) 1D ^13^C NMR spectra of MBPA-C-FG30-LSAFTG/AAO20 and MBPA-C-FG30-4G/AAO20 in dry states, obtained with ^1^H-^13^C cross-polarization or with direct pulsing. Samples contained 45.7 mg of AAO20 and 0.27 μmol of MBPA-C-FG30-LSAFTG or 28.3 mg of AAO20 and 0.14 μmol of MBPA-C-FG30-4G (samples 12 and 2 in Table S1, respectively). Green asterisks indicate MAS sideband signals and probe background signals. (B) Cross-polarized 1D ^31^P NMR of the MBPA-C-FG30-LSAFTG/AAO20 sample in dry and hydrated states. Signals centered at 24 ppm and −6 ppm arise from MBPA-C-FG30 phosphonate groups and residual surface phosphate groups, respectively. (C) 1D ^13^C NMR spectra of the samples in panel A after hydration with 0.2 M Bis-Tris buffer, pH 6.5, obtained with cross-polarization, direct pulsing, or ^1^H-^13^C INEPT. Green asterisks indicate natural-abundance ^13^C signals from the Bis-Tris buffer. Spectra in panels A, B, and C were acquired with 2048, 5120, and 2048 scans, respectively. All spectra were acquired at 20° C and 14.1 T, with MAS at 12.00 kHz.

Variations in incubation conditions included variations in incubation times, substitution of 25 mM acetate buffer for 0.2 M Bis-Tris buffer (sample 9 in Table S1), use of AAO with 40 nm pores (sample 3 in Table S1), incubation of three wafers in a single tube containing 1.0 ml of 1.0 mM MBPA-C-FG30 solution rather one wafer per tube (samples 2, 6, and 7 in Table S1), and crushing of AAO20 into small pieces prior to incubation (sample 7 in Table S1). Incubation times greater than 24 h did not produce higher loading values. Loading values with 1.0 ml of peptide solution per wafer were higher than with 0.33 ml of solution per wafer. Crushing of AAO20 prior to incubation did not increase the loading. Similar results were obtained with acetate and Bis-Tris buffers.

A final MBPA-C-FG30-LSAFTG/AAO20 sample was prepared by incubating three intact wafers in separate tubes for 24 h, with 1.0 ml of peptide solution in each tube (sample 12 in Table S1). NMR signal areas for this sample indicate 5.8 ± 0.1 nmol of MBPA-C-FG30-LSAFTG per milligram of AAO20, with a total of 0.27 μmol of MBPA-C-FG30-LSAFTG in the MAS rotor. Given the 15% porosity value discussed above and the 3.3 kDa molecular weight of MBPA-C-FG30, the average peptide concentration within AAO20 nanopores is approximately 90 mM or 300 mg/ml. Assuming a density of 1.4 g/ml for the peptide,^58^ about 21% of the AAO20 pore volume is filled by MBPA-C-FG30 molecules on average. This sample was used for NMR measurements in Figs. 2, 3, S10, and S11, as well as for spin relaxation measurements described below.

**Figure 3.**
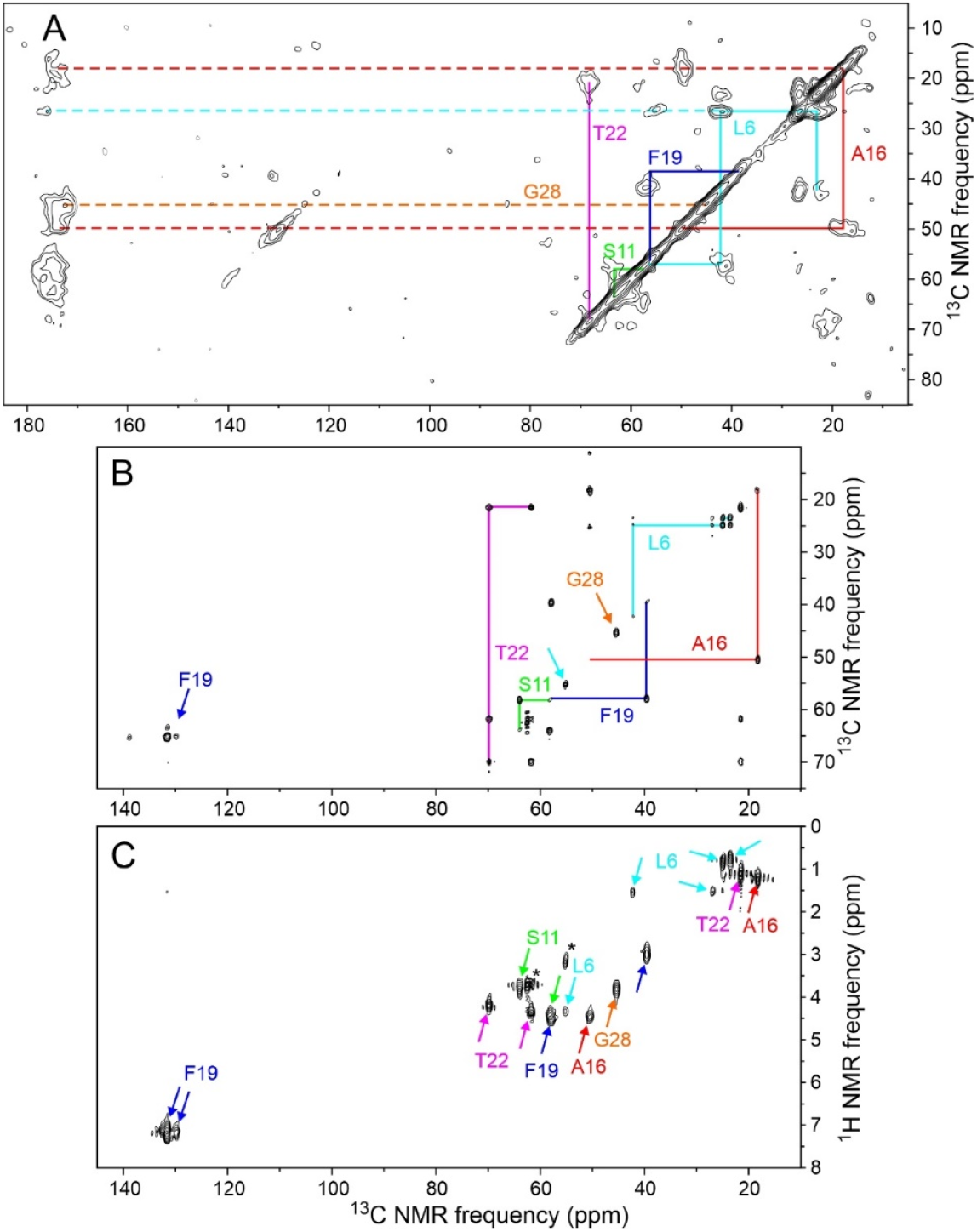
(A) 2D ^13^C-^13^C NMR spectrum of dry MBPA-C-FG30-LSAFTG/AAO20 (same sample as in Fig. 2A), obtained with a 30 ms DARR mixing period. Contour levels increase by successive factors of 1.5. (B) 2D ^13^C-^13^C NMR spectrum of hydrated MBPA-C-FG30-LSAFTG/AAO20 (same sample as in Fig. 2C), obtained with ^1^H-^13^C INEPT and a 12.5 ms isotropic ^13^C-^13^C mixing period. (C) 2D ^1^H-^13^C NMR spectrum of hydrated MBPA-C-FG30-LSAFTG/AAO20, obtained with INEPT polarization transfers. Asterisks indicate natural-abundance crosspeaks from the Bis-Tris buffer. Contour levels in panels B and C increase by successive factors of 1.8. All spectra were acquired at 20° C.

### Effects of hydration on FG30 conformation and dynamics in AAO nanopores

Fully hydrated states of MBPA-C-FG30/AAO20 samples were created by adding 0.2 M Bis-Tris buffer, pH 6.5, to the MAS rotors. The rotors were tightly sealed with cyanoacrylate glue, as verified by periodic measurements of the intensities of ^1^H NMR signals from liquid water throughout the experiments described below. In this context, it is worth emphasizing that AAO wafers are inherently hydrophilic, rapidly absorbing water and becoming almost fully transparent to visible light as the nanopores fill with water.

As shown in Fig. 2C, hydration has a dramatic effect on ^13^C NMR spectra of MBPA-C-FG30-LSAFTG/AAO20 and MBPA-C-FG30-4G. Directly-pulsed ^13^C NMR signals become sharper (0.15 ppm FWHM linewidths for doublet components of the T22 ^13^C_γ_ line at 20° C) while retaining the same signal areas as in the dry state (ratio of areas equal to 1.00 ± 0.05 for signals between 80 ppm and 0 ppm). Cross-polarized ^13^C signals become weak, with the total aliphatic signal area being reduced by a factor of approximately 18 relative to the dry state. Moreover, in the hydrated state it becomes possible to observe strong, sharp ^13^C NMR signals from aliphatic sites after polarization transfer from ^1^H spins with the INEPT technique^49^ and with low-power ^1^H decoupling. These observations show that FG30 peptide chains become highly dynamic within water-filled AAO20 nanopores, executing large-amplitude motions on submicrosecond time scales that strongly attenuate magnetic dipole-dipole couplings among nuclei. Molecular motions in the hydrated state also average out the local conformational and environmental differences that lead to inhomogeneously broadened NMR lines in the dry state.

(As is well-known, CP-based spectra depend on strong magnetic dipole-dipole couplings to ^1^H nuclei and therefore show strong signals only from sites with restricted mobility. INEPT-based spectra depend on weaker scalar couplings to ^1^H nuclei. Therefore, under our measurement conditions of moderate-speed MAS and low-power ^1^H decoupling, INEPT-based spectra show signals only from sites that undergo rapid, large-amplitude molecular motions. In principle, all sites contribute to DP-based spectra, regardless of mobility.)

In contrast to the ^13^C NMR signals, ^31^P NMR signals from MBPA-C-FG30-LSAFTG/AAO20 are not strongly affected by hydration, with linewidths and signal areas in cross-polarized spectra being nearly equal in dry and hydrated states (Fig. 2B). This observation is consistent with covalent bonding of the MBPA phosphonate group to the AAO20 nanopore wall surfaces, which renders the phosphonate group immobile regardless of hydration. In contrast, if MBPA-C-FG30 was associated with AAO20 without covalent bonding between MBPA phosphonates and nanopore walls, one would expect the ^31^P NMR linewidth to be reduced by molecular motions in the hydrated state (as observed for the ^13^C NMR linewidths in Fig. 2C).

Fig. 3 shows 2D NMR spectra of MBPA-C-FG30-LSAFTG/AAO20, including a 2D ^13^C-^13^C spectrum of the dry sample obtained with ^1^H-^13^C cross-polarization, high-power ^1^H decoupling in t_1_ and t_2_ periods, and a 30 ms DARR mixing period (Fig. 3A), a 2D ^13^C-^13^C spectrum of the hydrated sample obtained with ^1^H-^13^C INEPT, low-power ^1^H decoupling in t_1_ and t_2_ periods, and a 12.5 ms isotropic mixing (TOCSY) period (Fig. 3B), and a 2D ^1^H-^13^C spectrum of the hydrated sample obtained with ^1^H-^13^C INEPT and low-power ^1^H decoupling in the t_2_ period (Fig. 3C). All 2D spectra in Fig. 3 were acquired at 20° C. The 2D ^13^C-^13^C spectrum of the dry sample in Fig. 3A resembles the 2D ^13^C-^13^C spectrum of lyophilized MBPA-C-FG30-LSAFTG (Fig. S6B), but with significant differences in positions and intensities of crosspeaks that suggest differences in peptide conformational distributions between the dry, AAO20-bound state and the lyophilized state. Crosspeak widths are greatly reduced in the 2D ^13^C-^13^C and 2D ^1^H-^13^C spectra of the hydrated sample in Figs. 3B and 3C (<0.6 ppm for ^13^C, limited by pulse sequence conditions, 0.10 ppm for ^1^H).

Chemical shifts from 2D spectra in Figs. 3 and S6 are listed in Table 1. In the hydrated state, ^1^H and ^13^C chemical shifts are nearly identical to random-coil values,^59^ consistent with rapid exchange within a broad distribution of conformations. In the dry, AAO20-bound state and the lyophilized state, ^13^C chemical shifts measured at the centers of inhomogeneously broadened crosspeaks deviate more strongly from random-coil values.

**Table 1:**
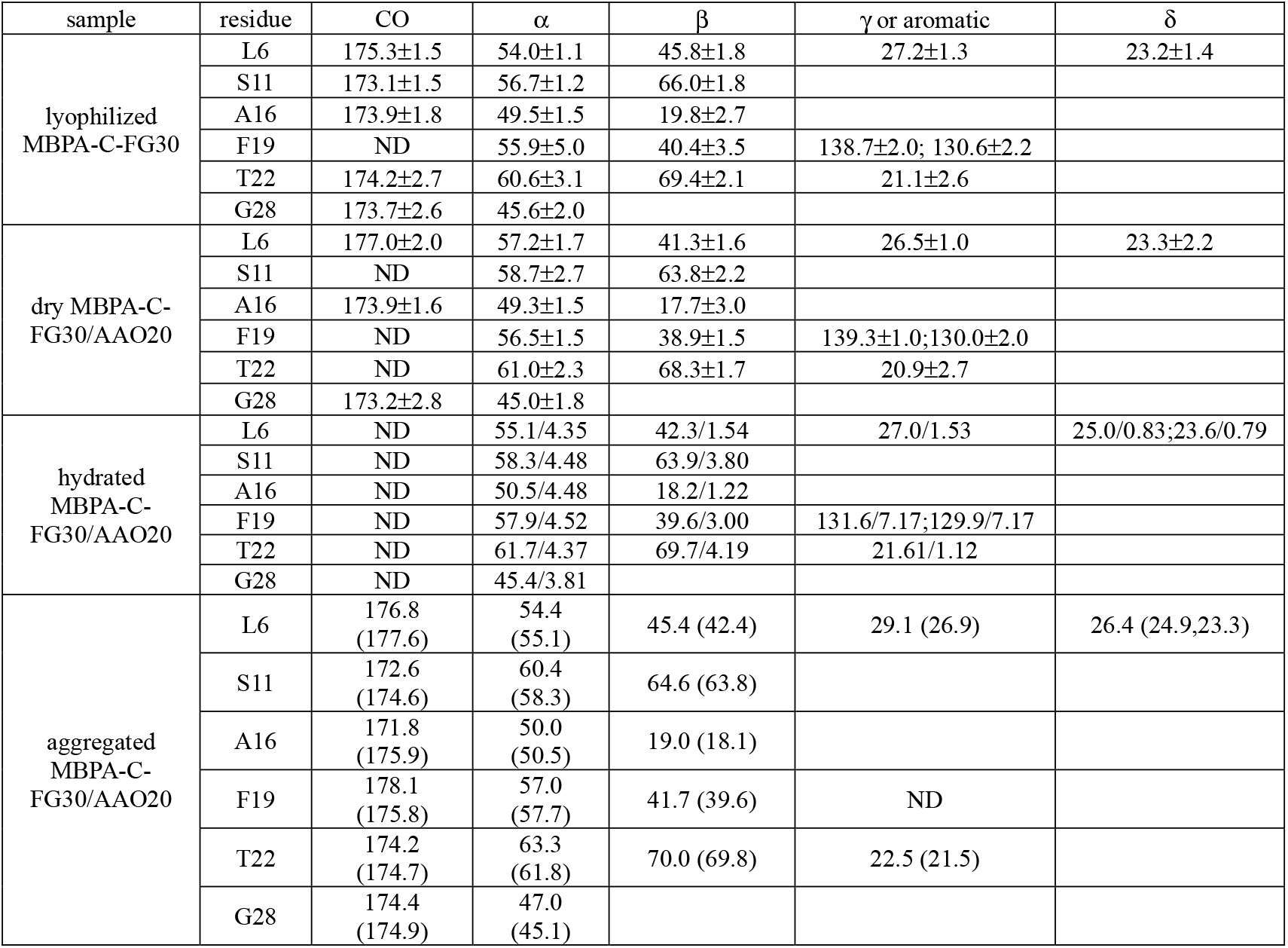
NMR chemical shifts (ppm relative to DSS) determined from 2D spectra at 20° C of lyophilized MBPA-C-FG30-LSAFTG, dry and hydrated MBPA-C-FG30-LSAFTG/AAO20, and aggregated MBPA-C-FG30-LSAFTG. For hydrated MBPA-C-FG30-LSAFTG/AAO20, both ^13^C and ^1^H chemical shifts are given. For other states, only ^13^C chemical shifts were determined. In lyophilized and dry states, chemical shift values were measured from center-of-mass crosspeak positions. Uncertainties represent FWHM linewidths. “ND” indicates values that could not be determined due to insufficient resolution or crosspeak intensity. In the hydrated and aggregated states, uncertainties are ±0.1 ppm for ^13^C and ±0.02 ppm for ^1^H. Random-coil values^59^ are given in parentheses along with ^13^C chemical shifts for the aggregated state.

### Absence of major temperature-dependent conformational changes for highly concentrated FG30 in AAO nanopores

To test for a possible temperature dependence of conformational preferences in hydrated MBPA-C-FG30-LSAFTG/AAO20, we recorded 1D ^13^C NMR spectra at temperatures between −5° C and 50° C. Spectra recorded with direct pulsing of ^13^C spins show sharper lines at 50° C than at lower temperatures, but total signal areas are independent of temperature to within experimental error (Fig. S10A). Spectra recorded with ^1^H-^13^C INEPT show sharper lines and larger signal areas at higher temperatures, indicating greater ^1^H-^13^C polarization transfer efficiencies at higher temperatures due to reductions in transverse spin relaxation rates (Fig. S10B). Temperature-dependent changes in ^13^C chemical shifts are less than 0.1 ppm for most ^13^C-labeled sites, except that the chemical shifts of the C_δ1_ and C_δ2_ sites of L6, the C_β_ sites of S11 and A16, the C_γ_ site of T22, and the C_α_ site of G28 increase by 0.25-0.66 ppm with increasing temperature. The A16 C_β_ site shows the largest change with temperature (0.66 ppm from −5° C to 50° C). For comparison, Kjaergaard *et al*. report a 0.0047 ppm per degree temperature dependence of the random-coil C_β_ site of alanine.^60^

Signals in spectra recorded with ^1^H-^13^C cross-polarization are generally weaker than in spectra recorded with direct pulsing, indicating inefficient cross-polarization due to attenuation of ^1^H-^13^C dipole-dipole couplings by molecular motions (Fig. S10C). From −5° C to 50° C, the ratio of the total aliphatic signal area per scan in CP spectra to the total aliphatic signal area per scan in DP spectra decreases from 0.48 ± 0.07 to 0.21 ± 0.03. The fact that signals in ^13^C CP spectra do not drop to zero at high temperatures indicates that rapid motions in MBPA-C-FG30-LSAFTG/AAO20 are not fully isotropic for all peptide molecules. At 50° C, the strongest signals in the cross-polarized ^13^C spectrum arise from aliphatic carbons of L6, the labeled residue that is closest to the point of attachment to the nanopore wall, where molecular motions are likely to be most restricted (Fig. S10C). Anisotropy of motion is discussed further below.

2D ^13^C-^13^C and ^1^H-^13^C spectra, with polarization transfers driven by scalar couplings, also show larger crosspeak intensities and sharper crosspeaks at higher temperatures but are otherwise nearly independent of temperature (Fig. S11). The absence of more strongly temperature-dependent chemical shifts shows that conformational preferences of MBPA-C-FG30 are not strongly temperature-dependent in the AAO20-bound, hydrated state.

### Absence of a temperature-driven phase transition for highly concentrated FG30 in AAO nanopores

Experimental studies of liquid-liquid phase-separation and hydrogel formation by FG-repeat domains in free solution^18–21^ suggest that a related phase transition might occur in the highly concentrated FG30 system within AAO20 nanopores. Since phase transitions generally involve changes in time scales or amplitudes of molecular motions that produce discontinuities in the temperature dependences of nuclear spin relaxation times,^61–62^ we therefore measured the temperature dependences of five relaxation times from −5° C to 50° C, namely the ^1^H and ^13^C spin-lattice relaxation times T_1_^H^ and T_1_^C^ (pulse sequences in Figs. S2H and S2K), the ^1^H and ^13^C transverse relaxation times T_2_^H^ and T_2_^C^ (pulse sequences in Figs. S2J and S2L), and the ^1^H rotating-frame relaxation time T_1ρ_^H^ (pulse sequence in Fig. S2I). All relaxation times were measured as build-up or decay times of ^13^C signals, allowing separate measurements for sites with resolved lines in 1D ^13^C spectra. ^1^H relaxation times were measured through ^13^C signals after INEPT polarization transfers. T_2_^C^ values were measured from spin echo decays, using frequency-selective ^13^C π pulses to minimize effects of ^13^C-^13^C scalar couplings.

Fig. 4 shows examples of spin relaxation data and the temperature dependences of relaxation times extracted from these data. The full sets of data appear in Figs. S12-S16. Values of relaxation times were extracted from these data by fitting with single-exponential or, in cases where single-exponential fits were deemed inadequate (see Materials and Methods), stretched-exponential functions. Best-fit values of relaxation times and stretching parameters β are given in Tables S2-S6. In all cases, fits converged to well-defined values of these parameters.

**Figure 4.**
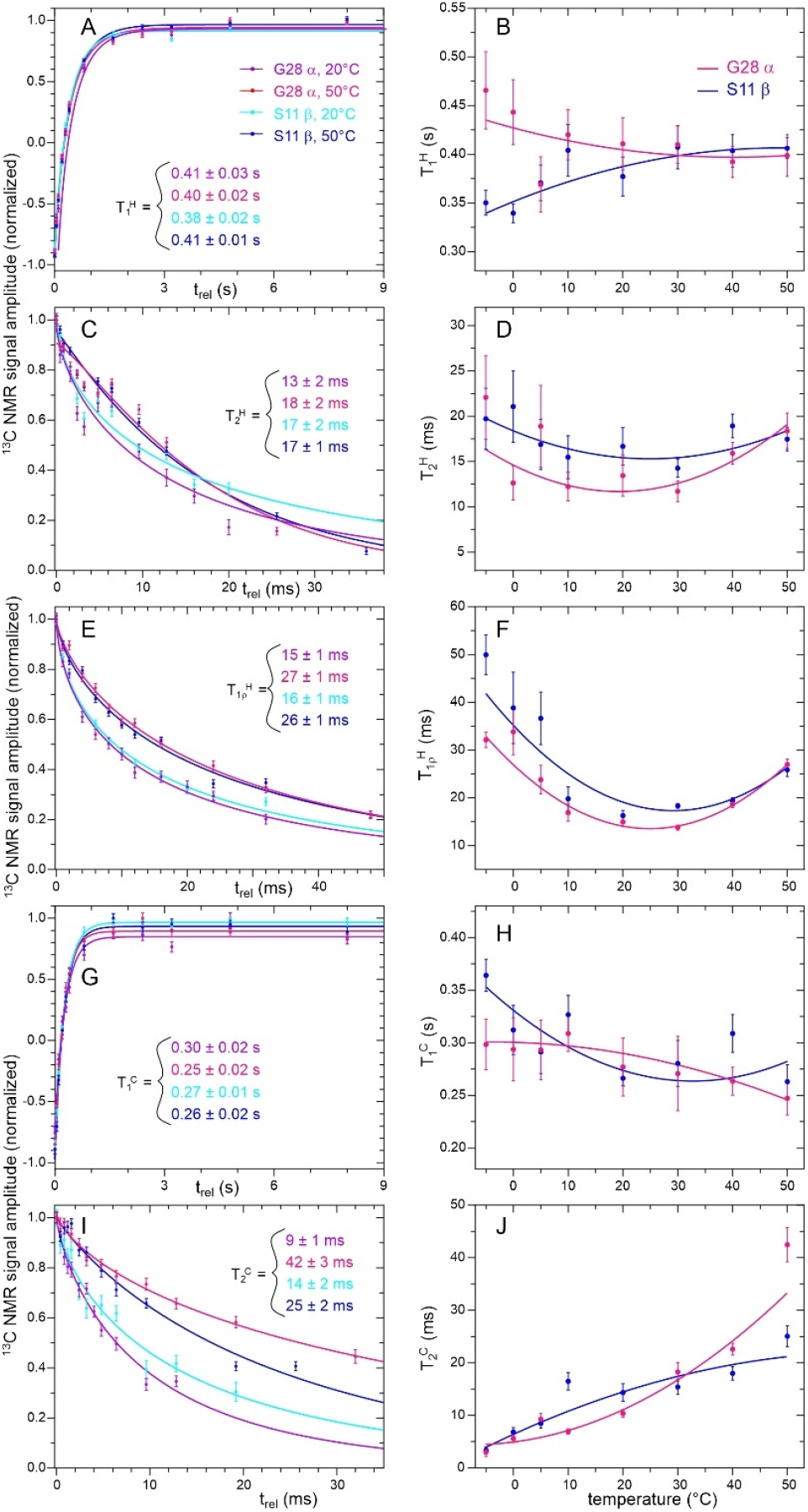
Examples of measurements of nuclear spin relaxation times for hydrated MBPA-C-FG30-LSAFTG/AAO20. (A,C,E,G,I) Measurements of ^13^C-detected T_1_ ^H, 13^C-detected T_2_^H, 13^C-detected T_1 ρ_^H^, T_1_^C^, and T_2_^C^ values for the S11 C_β_ and G28 C_α_ sites at 20° C and 50° C. Solid lines are stretched-exponential fits in panels A, C, E, and I, and single-exponential fits in panel G, yielding the indicated relaxation times. Error bars represent uncertainties derived from the RMS noise in the ^13^C NMR spectra. (B,D,F,H,J) Temperature dependences of the spin relaxation times. Solid lines are second-order polynomial fits, intended to serve as guides to the eye. Error bars represent standard errors in the fitted relaxation time values.

Plots of the temperature dependences of all measured relaxation times appear in Figs. S17-S21. With only two exceptions discussed below, all relaxation times show smooth dependences on temperature that are adequately fit with second-order polynomial functions. This observation argues strongly against a temperature-driven phase transition in MBPA-C-FG30/AAO20. In particular, the relaxation data and the temperature-dependent NMR spectra (Figs. S10 and S11) rule out a transition between a state in which peptide chains move independently of one another and a state in which neighboring peptide chains form long-lived clusters or cohesive assemblies through intermolecular contacts with microsecond or longer lifetimes. Formation of long-lived clusters above or below a specific transition temperature would be expected to produce an abrupt broadening of NMR lines and reduction in T_1ρ_^H^, T_2_^H^, and T_2_^C^ values, due to reductions in the amplitudes or increases in the time scales of molecular motions. If the phase transition involved major changes in site-specific conformational distributions, ^13^C NMR chemical shifts would also be expected to change abruptly. Rather than a phase transition, our data indicate continuous and gradual increases in molecular mobility with increasing temperature (analyzed further below).

T_2_^C^ values for the A16 C_β_ and T22 C_γ_ sites have apparent sigmoidal dependences on temperature (Fig. S21), increasing by factors of 2.8 and 3.2 from 20° C to 30° C. Although the increase in T_2_^C^ values for the A16 C_β_ and T22 C_γ_ sites between 20° C and 30° C suggests an approximate three-fold reduction in orientational correlation times for methyl axis directions at these sites around 25° C, similar increases in T_2_ ^C^ values for A16 C_α_ and T22 C_β_ sites would then be expected but are not observed.

### Interpretation of nuclear spin relaxation data for FG30 in AAO nanopores

All T_1_^H^ values are in the 0.3-0.6 s range and nearly independent of temperature (Fig. S17). T_1_^C^ values are in the 0.2-0.6 s range, either increasing or decreasing by less than 30% between −5° C and 50° C for most sites (Fig. S20). T_1_^C^ of the T22 C_γ_ site has the strongest temperature dependence, increasing by 50%. At a qualitative level, these observations indicate that motional amplitudes with correlation times of roughly 0.1-1 ns, which primarily contribute to T_1_^H^ and T_1_^C^ relaxation, are not strongly temperature dependent.

T_1_^H^ and T_2_^H^ values are in the 5-60 ms range, varying with temperature and among sites. T_1 ρ_^H^ values for most sites exhibit non-monotonic temperature dependences, with minima at temperatures in the 10-30° C range (Fig. S18). T_2_^H^ values for some sites (S11 C_β_, A16 C_α_, F19 C_β_, G28 C_α_, and the combined signal from S11 C_α_ and F19 C_α_) have similar non-monotonic temperature dependences (Fig. S19). For these sites, T_1 ρ_^H^ ≈ T_2_^H^ near 25° C, but values of T_1 ρ_^H^ are 1.5-2.0 times larger than values of T_2_^H^ at the highest and lowest temperatures. T_2_^C^ values increase with increasing temperature, from the 2-20 ms range at −5° C to the 4-60 ms range at 50° C (Fig. S21).

T_2_^H^ values for methyl sites (L6 C_δ1_ and C_δ2_, A16 C_β_, T22 Cγ) and for the F19 sidechain increase by factors of roughly 2-3 with increasing temperature, while other T_2_^H^ values are more weakly temperature-dependent (Fig. S19). T_2_^H^ values for the aromatic sidechain sites of F19 are smaller than T_1ρ_^H^ values over the entire temperature range, suggesting a role for slow sidechain ring motions. Slow ring motions may also account for the relatively small T_2_^C^ values for F19 sidechain sites.

The complexity of molecular motions in hydrated MBPA-C-FG30/AAO20, complications due to ^1^H-^1^H cross-relaxation, and other multi-spin effects preclude detailed analysis of the complete set of relaxation data. However, certain data are amenable to approximate treatments by simple models. Specifically, ^13^C relaxation data for α-carbon sites of S11/F19 and A16 and the β-carbon site of T22 can be treated approximately as data for two-spin ^1^H-^13^C systems with a one-bond internuclear distance. ^1^H relaxation data detected through β-carbon sites of L6, S11, and F19 and the α-carbon site of G28 can also be analyzed, since these methylene sites can be treated approximately as two-spin ^1^H-^1^H systems with additional one-bond couplings to one ^13^C spin. We assume that relaxation is driven by the random time-dependences of homonuclear and heteronuclear dipole-dipole couplings produced by orientational motions. We then use expressions for T_1_ and T_2_ relaxation rates in dipole-coupled two-spin systems derived by Solomon^63^ and for T_1ρ_ relaxation derived by Jones^64^ and by Bleich and Glasel^65^:

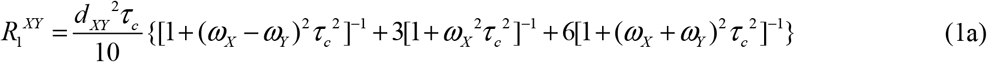

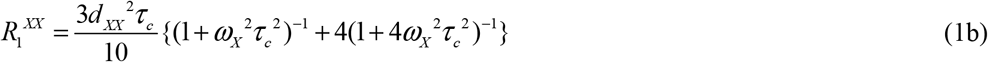

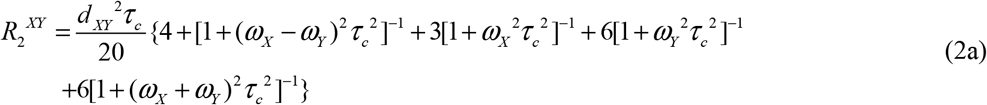

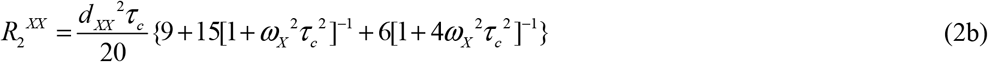

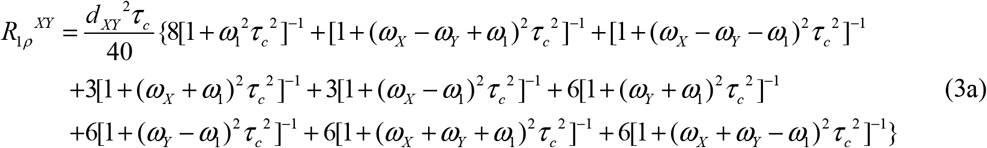

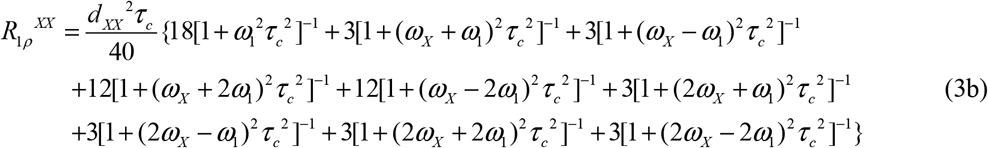

R_1_^XY^ is the contribution to T_1_ relaxation of X due to X-Y coupling, with X = ^1^H and Y = ^13^C or vice versa. R_1_^XX^ is the contribution to T_1_ relaxation of X due to X-X coupling. Similar definitions apply to R_2_ ^XY^, R_2_^XX^, R_1 ρ_^XY^, and R_1 ρ_ ^XX^. For simplicity, we ignore cross-correlations between different couplings, so that *T*_1_ ^*H*^ = (*R*_1_ ^*HH*^ + *R*_1_ ^*HC*^)^−1^, *T*_2_ ^*H*^ = (*R*_2_ ^*HH*^ + *R*_2_ ^*HC*^)^−1^, and *T*_1*ρ*_ ^*H*^ = (*R*_1*ρ*_ ^*HH*^ + *R*_1*ρ*_ ^*HC*^)^−1^. Dipole-dipole coupling constants are *d*_*XY*_ = *ħ*^2^*γ*_*X*_ *γ*_*Y/*_ *r*_*XY*_^3^ and *d*_*XX*_ = *ħ*^2^*γ*_*X*_^2^ *r*_*XX*_^3^, with gyromagnetic ratios γ_X_ and γ_Y_ and internuclear distances r_XY_ and r_XX_. For a 0.110 nm one-bond ^1^H-^13^C distance and a 0.176 nm methylene ^1^H-^1^H distance,^66^ d_XY_ = 1.426 × 10^5^ rad/s and d_XX_ = 1.384 × 10^5^ rad/s. In the case of ^1^H relaxation data for the methylene sites, our use of Eqs. (1-3) ignores possible cross-correlations between randomly fluctuating ^1^H-^1^H and ^1^H-^13^C couplings.^67^

Eqs. (1-3) apply to isotropic motion with a single correlation time τ_c_, *i*.*e*., orientational correlation functions of the form *C*(*t*) = *C*(0) exp(−*t τ*_*c*_). A single correlation time does not fit the experimental data, since (for example) T_1_^C^ ≈ 0.5 s would require τ_c_ ≈ 0.1 ns but T_2_ ≈ 20 ms would require τ_c_ ≈ 10 ns. We therefore assume that the correlation functions have the form *C*(*t*) = *C*(0)[*a*_1_ exp(−*t τ*_*c*1_) + (1− *a*_1_) exp(−*t τ*_*c*2_)]. The net relaxation rates are then sums of rates given by Eqs. (1-3) with τ_c_ = τ_c1_ and τ_c_ = τ_c2_, weighted by a_1_ and 1 − a_1_, respectively.

For each site and each temperature, we fit the three experimental ^1^H or two experimental ^13^C relaxation times by varying τ_c1_ and τ_c2_ over 10-500 ns and 0.1-9 ns ranges, respectively. Thus, τ_c2_ represents motions that partially randomize local molecular orientations on a short time scale, while τ_c1_ represents slower motions that complete the randomization of local molecular orientations. As described in the Materials and Methods, our fitting procedure produces probability-weighted average values of τ_c1_, τ_c2_, and a_1_ and standard deviations in these values, with probabilities proportional to exp[−*χ*^2^ (*τ*_*c*1_,*τ*_*c* 2_)]. The results from analyses of ^13^C and ^1^H relaxation data are listed in Tables S7 and S8, respectively.

^13^C relaxation data for all sites yield fitted values of τ_c1_ and τ_c2_ in the 80-200 ns and 1.0-1.4 ns ranges, respectively, and fitted values of a_1_ in the 0.12-0.47 range (Table S7). Fitted values of τ_c2_ and a_1_ tend to decrease with increasing temperature, although the trends are not pronounced. ^1^H relaxation data for all sites yield fitted values of τ_c1_ and τ_c2_ in the 40-120 ns and 0.3-0.9 ns ranges, respectively, and fitted values of a_1_ in the 0.03-0.29 range (Table S8). Temperature-dependent trends are not discernible.

Standard deviations for the fitted values in Tables S7 and S8 are similar to the probability-weighted average values themselves, which (not surprisingly) means that adequate fits to the data can be obtained with substantial variations in these correlation times. Nonetheless, a non-zero value of a_1_ is required. The value of a_1_ can be interpreted as a scaling factor for dipole-dipole couplings caused by motional averaging within time periods less than τ_c1_. If this motional averaging occurs by randomization of bond directions within a cone with half-angle β, then 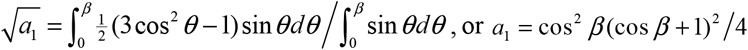. If a_1_ ≈ 0.25, as indicated by the relaxation data, then β ≈ 52°. If a_1_ ≈ 0.10, then β ≈ 64°. Thus, the non-zero value of a_1_ implies substantial restriction in the amplitude of the more rapid motions.

For completeness, we note that Eqs. (1-3) do not include cross-relaxation effects, *i*.*e*., effects on the X relaxation data that would depend on the polarization state of Y.^63^ Experimentally, we find that inversion-recovery T_1_^H^ data for MBPA-C-FG30-LSAFTG/AAO20 are unaffected by application of a ^13^C π pulse at the beginning of the recovery period. Although experimental inversion-recovery T_1_^C^ data are altered when a ^1^H π pulse is applied at the beginning of the recovery period, simulations for a ^1^H-^13^C system with τ_c1_ = 50 ns, τ_c2_ = 1 ns, and a_1_ = 0.25 show that the ^13^C recovery curve without a ^1^H π pulse but with cross-relaxation is nearly identical to the ^13^C recovery curve with cross-relaxation set to zero. Cross-relaxation effects are therefore negligible in our relaxation measurements (which do not include the extra π pulses).

### Isotropy of FG30 motion in AAO nanopores

Fully isotropic motion on sub-microsecond time scales would average ^1^H-^1^H and ^1^H-^13^C dipole-dipole couplings to zero. The observation of weak signals in cross-polarized ^13^C NMR spectra of hydrated MBPA-C-FG30/AAO20 (Fig. 2C) suggests that motional averaging of these couplings may be incomplete, meaning that molecular motions are not fully isotropic within the hydrated AAO20 pores on time scales up to several microseconds. Small residual dipole-dipole couplings would be averaged out efficiently by MAS at 12.00 kHz but could affect measurements at low MAS frequencies. Therefore, as a test for possible anisotropy of motion, we performed T_2_^H^ measurements on hydrated MBPA-C-FG30/AAO20 at MAS frequencies of 8.0 kHz, 4.0 kHz, 2.0 kHz, and 1.2 kHz, using the ^13^C-detected ^1^H spin-echo pulse sequence (Fig. S2J) and using the decay of total signal from ^13^C_α_ and ^13^C_β_ sites with increasing t_rel_ to determine T_2_^H^. These measurements show that T_2_^H^ at 24° C decreases from 12.2 ± 0.8 ms at 8.0 kHz MAS to 8.0 ± 0.1 ms at 1.2 kHz MAS, a relatively small effect (Fig. S22A,B). Comparison with numerical simulations of the ^1^H spin-echo signal decay for a dipole-coupled four-spin system (Fig. S22C) indicates that rapid molecular motions in MBPA-C-FG30/AAO20 reduce dipole-dipole couplings by a factor of at least 50 relative to their values in the absence of molecular motion. Equating the net motional scaling factor for all sub-microsecond motions with cos *β* (cos *β* + 1) 2 as discussed above yields β > 163°, where β = 180° would represent fully isotropic motion.

### Aggregation of FG30 in free solution

Results presented above show that MBPA-C-FG30 molecules are highly dynamic in aqueous solution at concentrations near 90 mM and over a wide temperature range when tethered to the nanopore walls of AAO20. Sharp lines and random-coil chemical shifts in NMR spectra (Figs. 2C, 3B, and 3C; Table 1) indicate the absence of oligomerization or aggregation, and no spectral changes occur over many weeks of measurements. In contrast, untethered MBPA-C-FG30 aggregates at 10 mM concentration in 0.2 mM Bis-Tris buffer, pH 6.5 when incubated at 4° C, 24° C, or 37° C. Negative-stain TEM images in Fig. 5, after six days of incubation, show that the aggregated material consists of straight, unbranched assemblies with morphologies and dimensions typical of amyloid fibrils^28,40,44^.

**Figure 5.**
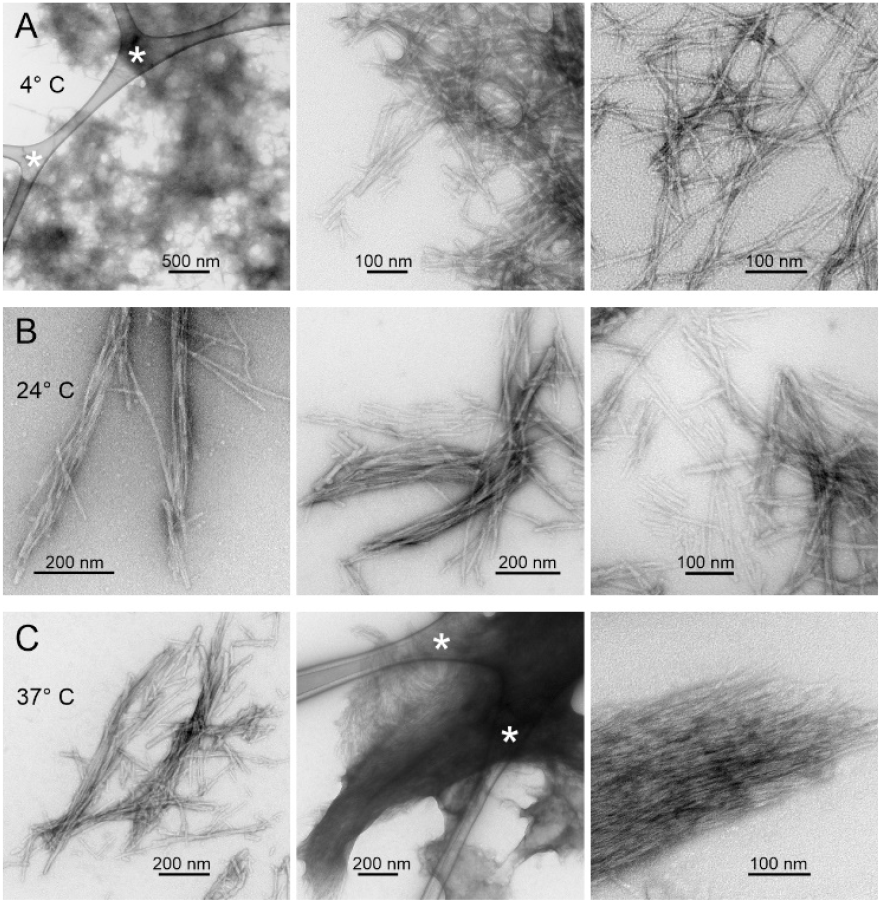
TEM images of aggregated MBPA-C-FG30, negatively stained with uranyl acetate. Aggregated material developed spontaneously at 10 mM peptide concentration in 0.2 M Bis-Tris buffer, pH 6.5, during incubation at 4° C (A), 24° C (B), or 37° C (C). Asterisks indicate features from the carbon lace that supports the thin carbon films to which aggregated MBPA-C-FG30 is adsorbed on the TEM grids.

To estimate the equilibrium solubility of MBPA-C-FG30, 7.1 mM solutions in the same buffer (23 mg/ml, 50-140 μl volumes) were incubated at 4° C, 24° C, and 40° C for four days, with brief bath sonication (10-15 s) once per day to accelerate the approach to equilibrium. After incubation, solutions were centrifuged at 16,000 × g for 15 min, yielding visible pellets at each temperature. Supernatants were then passed through 0.1 μm centrifugal filters (Millipore Ultrafree, 0.5 ml volume) and diluted with H_2_O by factors of four. Peptide concentrations were determined from absorbance measurements at 260 nm, performed in triplicate, using an extinction coefficient of 1070 M^−1^ cm^−1^ that was determined from the absorbance of the initial solutions. Based on these measurements, equilibrium solubilities of MBPA-C-FG30 in 0.2 mM Bis-Tris buffer, pH 6.5, are 4.5 ± 0.2 mM, 1.60 ± 0.04 mM, and 1.92 ± 0.08 mM at 4° C, 24° C, and 40° C. Thus, the concentration at which MBPA-C-FG30 remains unaggregated and dynamic when tethered to nanopore walls of AAO20 greatly exceeds the equilibrium solubilities in free solution. Figs. 6A and 6B show cross-polarized 1D ^13^C and directly-pulsed 1D ^31^P NMR spectra of aggregated MBPA-C-FG30-LSAFTG, prepared by incubation of a 10 mM solution at 24° C. Fig. 6C shows a 2D ^13^C-^13^C NMR spectrum, obtained with ^1^H-^13^C CP, a 30 ms DARR mixing period, and high-power ^1^H decoupling. For these measurements, aggregated material was pelleted by ultracentrifugation (278000 × g, 120 min, 20° C), then transferred by centrifugation into the MAS NMR rotor without lyophilization or drying. The aggregated material was therefore fully hydrated with 0.2 M Bis-Tris buffer, pH 6.5. Although the 1D ^13^C and 2D ^13^C-^13^C spectra in Fig. 6 were obtained with NMR measurement conditions appropriate for immobilized molecules, signals are sharp (0.6-1.0 ppm FWHM linewidths), consistent with an ordered peptide conformation and only small-amplitude motion at the ^13^C-labeled sites in the aggregated state (except for the aromatic sidechain of F19, from which signals are weak). Aliphatic signal areas in 1D ^13^C NMR spectra obtained with direct pulsing or with ^1^H-^13^C INEPT were 1.7 or 3.8 times smaller, respectively, than the aliphatic signal area in Fig. 6A.

**Figure 6.**
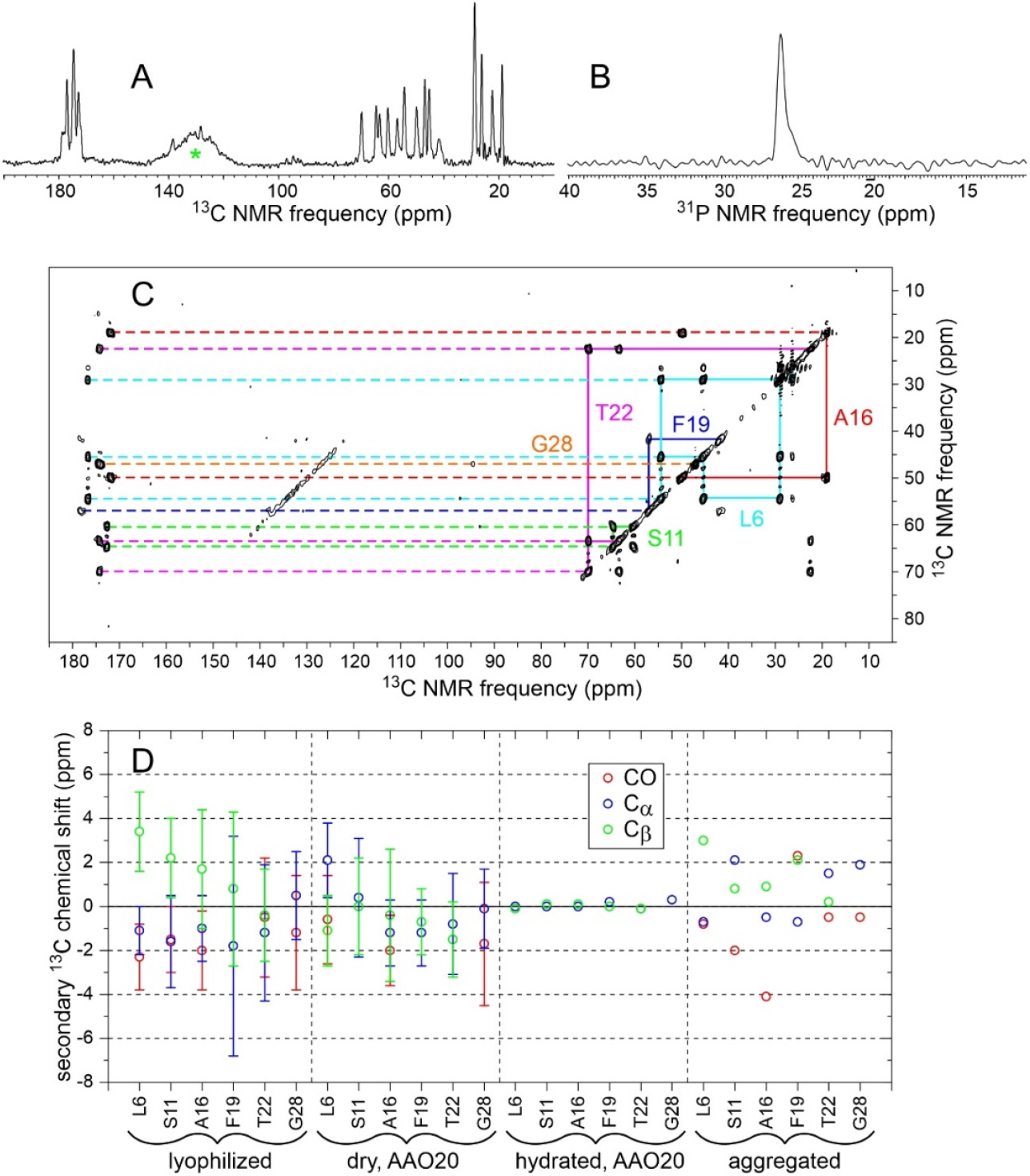
(A,B) Cross-polarized 1D ^13^C NMR and directly-pulsed 1D ^31^P spectra of aggregated MBPA-C-FG30-LSAFTG in a fully hydrated state. Green asterisk in panel A indicates a probe background signal. (C) 2D ^13^C-^13^C NMR spectrum of the same sample, obtained with ^1^H-^13^C cross-polarization and a 30 ms DARR mixing period. Contour levels increase by successive factors of 1.8. All spectra were recorded at 20° C. (D) Secondary ^13^C chemical shifts for backbone CO, C_α_, and C_β_ sites determined from 2D ^13^C-^13^C NMR spectra of MBPA-C-FG30-LSAFTG in lyophilized, dry AAO20-bound, hydrated AAO20-bound, and aggregated states (Figs. S6B, 3A, 3B, and 6C, respectively). Error bars for lyophilized and dry AAO20 bound states represent FWHM ^13^C linewidths, determined from 1D slices through crosspeaks in the 2D spectra.

The ^31^P linewidth in Fig. 6B is 20 times smaller than the ^31^P linewidth for hydrated MBPA-C-FG30-LSAFTG/AAO20 in Fig. 2B. The ^31^P signal area in the directly-pulsed spectrum is 2.4 times larger than the signal area in a cross-polarized spectrum. In addition, the ^31^P spin-lattice relaxation time in aggregated MBPA-C-FG30-LSAFTG at 20° C, determined from inversion-recovery measurements, is 0.22 ± 0.02 s, compared with 15 ± 5 s in hydrated MBPA-C-FG30-LSAFTG/AAO20 (Fig. S23). These results indicate that phosphonate groups execute rapid, large-amplitude motions in hydrated, aggregated MBPA-C-FG30. The slow ^31^P spin-lattice relaxation in MBPA-C-FG30/AAO20 provides additional evidence for covalent bonding of phosphonate groups to AAO nanopore wall surfaces.

The observation of a single set of crosspeaks in the 2D ^13^C-^13^C NMR spectrum (Fig. 6C) indicates a homogeneous local structure in aggregated MBPA-C-FG30, despite the morphological heterogeneities in TEM images (Fig. 5). Secondary ^13^C chemical shifts in Fig. 6D suggest β-strand conformations at L6 and A16 (negative ^13^C_α_ and ^13^CO and positive ^13^C_β_ secondary shifts), but not at other labeled residues.

### Characterization of FG30 density within AAO nanopores by confocal fluorescence microscopy

To determine the dependence of peptide density on distance from the outer AAO20 surfaces into the nanopores, wafer pieces were incubated in a 1.0 mM MBPA-C-FG30-G solution in which 2.5% of the peptide molecules were fluorescently labeled with FITC. As shown in Fig. 7A, confocal fluorescence microscope images were recorded with excitation at 405 nm and detection at 514-585 nm, using 1.0 μm increments in the distance z perpendicular to the flat wafer surface. Plots of the average fluorescence intensity within the images as a function of z can then be interpreted as plots of peptide density versus z (averaged over directions perpendicular to z, and assuming that FITC-labeled peptide molecules have the same spatial distribution within AAO20 as unlabeled molecules). As shown in Figs. 7B and 7C, the peptide density increases with increasing incubation time. Peptide densities within the nanopores are larger at z values near the two outer surfaces than near the midpoint between these surfaces (z ≈ 25 μm), by factors of 1.4-2.2 after 24 h incubation. Moreover, the peptide densities are not symmetric about the midpoint.

**Figure 7.**
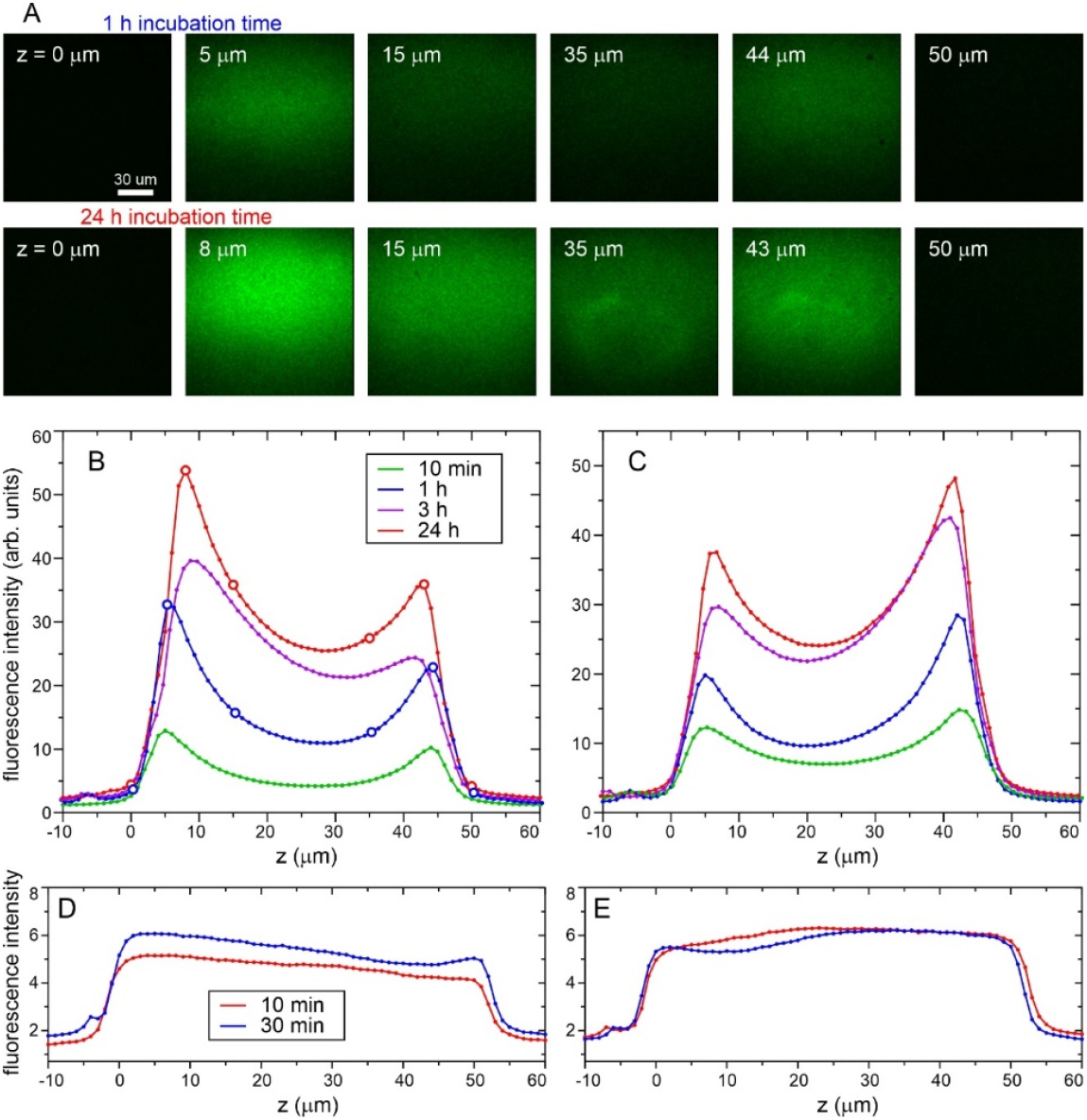
(A) Confocal fluorescence microscope images of AAO20 wafers after incubation at 40° C for 1 h or 24 h in 1.0 mM FITC-labeled MBPA-C-FG30 (2.5% labeling). Fluorescence was excited at 405 nm and collected at 514–585 nm. Images are shown for indicated distances z into the wafer, with z = 0 μm defined as a position of the confocal point slightly outside the wafer. (B) Dependence of fluorescence intensity on z for samples with the indicated incubation times, obtained by averaging the intensity within images at increasing z values. Open circles are the positions shown in panel A. (C) Fluorescence intensity profiles after flipping each sample in panel B, so that excitation light enters and fluorescence is detected from the opposite side of the wafer. (D) Fluorescence intensity profiles for AAO20 wafers that were incubated at 24° C in solutions containing 50 μg/ml fluorescein and 6 M GuHCl for 10 min or 30 min. (E) Fluorescence intensity profiles after flipping the samples in panel D.

Fluorescence intensity profiles in Figs. 7B and 7C were measured with opposite wafer surfaces directed towards the excitation light (*i*.*e*., by flipping the wafer pieces over in their glass-bottom microwell dishes). The fact that the profiles in Figs. 7B and 7C have opposite asymmetries implies that the asymmetries are not due to optical distortions. We speculate that the asymmetries of the fluorescence intensity profiles are a consequence of structural or chemical inequivalence between the two outer surfaces, arising from the inherent asymmetry of the AAO wafer fabrication process.^36–37^ The lower fluorescence intensity near the midpoint in these profiles may indicate that diffusion of MBPA-C-FG30 molecules into the midpoint of AAO20 nanopores is impeded by the accumulation of high densities of tethered MPBA-C-FG30 near the two ends of the nanopores. It is also conceivable that a fraction of the nanopores are not fully open over their entire 50 μm lengths or become partially blocked by extraneous particles in the incubating solutions.

For comparison, Figs. 7D and 7E show fluorescence intensity profiles for AAO20 wafer pieces that were incubated for short times in solutions of fluorescein, which we found to adsorb to the nanopore walls. Although these profiles are not fully symmetric, they do not show minima near z = 25 μm. Thus, small molecules can penetrate through the entire lengths of the nanopores to create a nearly uniform distribution of adsorbed species.

As an additional test of the uniformity of MBPA-C-FG30 density within AAO20 pores, we obtained AAO20 flakes with 5 μm thickness from InRedox LLC. We treated this thinner material with dilute H_2_SO_4_ as described above, then incubated it in 1.0 mM MBPA-C-FG30-4G at 40° C for 24 h. The signal area in the cross-polarized ^13^C NMR spectra of 4.9 mg of the final dried material indicated a loading of 5.13 ± 0.33 nmol per mg (sample 4 in Table S1 and Fig. S7), similar to the loading of wafers with 50 μm thickness. Together with the confocal microscopy results, these NMR measurements indicate that peptide densities in our samples may vary by ±35% along the lengths of AAO20 nanopores, but the densities are high throughout the material. A full understanding of the peptide density distribution will require additional measurements.

### Generality of the approach

To test the applicability of our approach to other peptides in nanopore environments, we incubated 53.0 mg of H_2_SO_4_-treated AAO20 wafer pieces at 40° C in 0.95 ml of 1.0 mM MBPA-C-melittin-GVA, 50 mM Bis-Tris, pH 6.5 for 50 h. After washing with H_2_O and drying under N^2^ gas, 1D ^13^C NMR spectra of the dry MBPA-C-melittin-GVA/AAO20 sample indicated a total of 0.40 ± 0.03 μmoles of peptide in 47.6 mg of crushed AAO20 (0.38 μmole based on the aliphatic signal area in the cross-polarized spectrum in Fig. 8A, 0.43 μmole based on the directly-pulsed spectrum in Fig. 8B). With the 3.2 kDa molecular weight of MBPA-C-melittin-GVA (Fig. S1H), this result implies a peptide concentration of approximately 140 mM (440 mg/ml) in the AAO20 nanopores.

**Figure 8.**
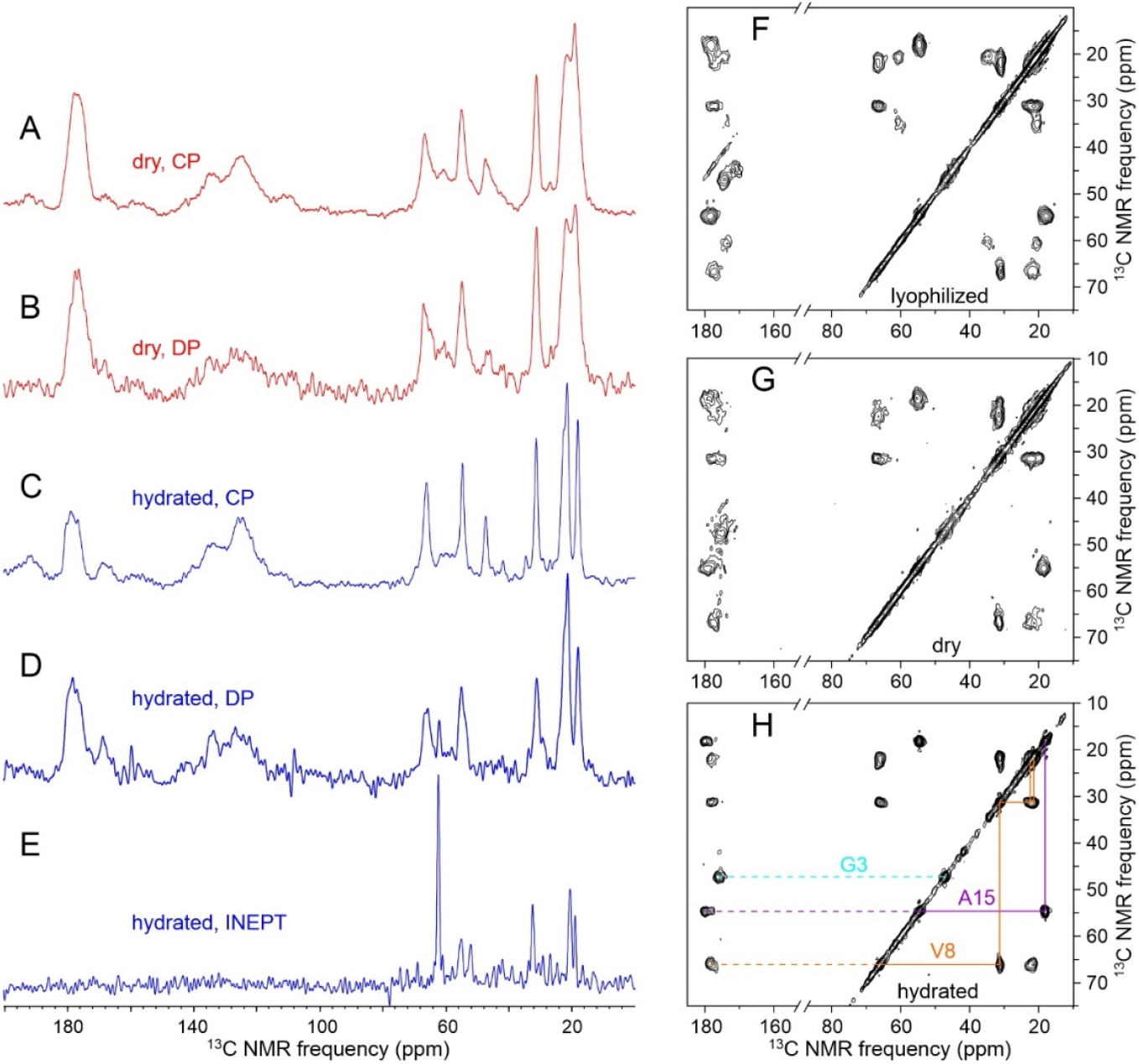
(A,B) 1D ^13^C NMR spectra of dry MBPA-C-melittin-GVA/AAO20, recorded with ^1^H-^13^C cross-polarization or direct pulsing and with 1024 scans. Total sample mass was 48.9 mg, including 0.40 ± 0.03 μmole of MBPA-C-melittin-GVA. (C,D) 1D ^13^C NMR spectra of MBPA-C-melittin-GVA/AAO20 after hydration with 50 mM Bis-Tris, pH 6.5, recorded with ^1^H-^13^C cross-polarization or direct pulsing and with 1024 scans. (E) 1D ^13^C NMR spectrum of the hydrated sample, recorded with ^1^H-^13^C INEPT and low-power ^1^H decoupling. Aliphatic signal areas per scan in panels A-E have ratios of 100.0:36.9:59.8:26.2:2.5. Approximately 30% of the dry sample was lost in the hydration process. (F,G,H) 2D ^13^C-^13^C NMR spectra of a 1.4 mg sample of lyophilized MPBA-C-melittin-GVA powder, the dry MBPA-C-melittin-GVA/AAO20 sample, and the hydrated MBPA-C-melittin-GVA/AAO20 sample. All 2D spectra used ^1^H-^13^C cross-polarization and 30 ms DARR mixing. Contour levels in 2D spectra increase by successive factors of 1.5. All spectra were recorded at 32° C and 17.5 T, with MAS at 12.5 kHz.

After hydrating the MBPA-C-melittin-GVA/AAO20 sample with 50 mM Bis-Tris, pH 6.5, strong signals were observed in 1D ^13^C NMR spectra obtained with ^1^H-^13^C cross-polarization and with direct pulsing of ^13^C polarization (Figs. 8C and 8D). Aliphatic signals in a 1D ^13^C NMR spectrum obtained with ^1^H-^13^C INEPT (Fig. 8E) were weaker than in the cross-polarized spectra by a factor of 24. This behavior of hydrated MBPA-C-melittin-GVA/AAO20 is opposite to the behavior of hydrated MBPA-C-FG30-LSAFTG/AAO20, where aliphatic signals in the INEPT ^13^C NMR spectrum were stronger than signals in the cross-polarized spectrum by an approximate factor of 7 (Fig. 2C).

Figs. 8F-8H show 2D ^13^C-^13^C NMR spectra of lyophilized C-melittin-GVA, dry MBPA-C-melittin-GVA/AAO20, and hydrated MBPA-C-melittin-GVA/AAO20, respectively. Again in contrast to observations for MBPA-C-FG30-LSAFTG (Figs. 3A, 3B, and S6B), crosspeak positions in the three 2D spectra in Fig. 8 are nearly the same, indicating similar peptide conformations. Crosspeaks in the hydrated state of MBPA-C-melittin-GVA/AAO20 are sharper than in the dry state (1.1-1.8 ppm versus 1.5-2.9 ppm FWHM), consistent with solvation and rapid small-amplitude motions. Comparison of ^13^C chemical shifts in Fig. 8H with standard random-coil values^59^ indicates an α-helical conformation at the isotopically labeled residues (176.1 and 47.4 ppm for G3; 178.4, 65.9, 31.2, 22.6, and 21.6 ppm for V8; 179.8, 54.6, and 18.2 ppm for A15). Thus, when melittin is tethered to walls of AAO20 nanopores at high concentrations, it adopts an α-helical conformation similar to the known conformation in melittin tetramers in free solution.^68–70^ The contrasting behavior of hydrated MBPA-C-melittin/AAO20 and hydrated MBPA-C-FG30/AAO20 implies that the random-coil-like properties of tethered FG30 at high concentrations in AAO20 nanopores are not universal for tethered peptides of similar size at similar concentrations.

## DISCUSSION

Results described above are significant both from the standpoint of sample preparation and from the standpoint of FG-repeat sequence behavior. Regarding sample preparation, our results show that: (i) Phosphonate moieties are readily attached to polypeptide chains through cysteine-maleimide linkages. In experiments described above, the phosphonate moieties were attached to the N-termini of FG30 and melittin peptides, but attachment to the C-termini or other positions are possible; (ii) Spontaneous reaction of phosphonates with aluminum oxide surfaces allows covalent tethering of polypeptides to nanopore walls in AAO. Reaction times on the order of 24 h result in high peptide densities within the nanopores; (iii) In the case of MBPA-C-FG30/AAO20, average peptide concentrations of 90 mM are achievable, corresponding to a 300 mg/ml density that is similar to the density of FG-repeat domains in the central channel of an NPC^8^; (iii) Importantly, covalent tethering to AAO nanopore walls is stable in aqueous solution near pH 7. We observed no changes in NMR spectra of hydrated MBPA-C-FG30-LSAFTG/AAO20 over more than 8 weeks of measurements, with sample temperatures ranging from −5° C to 50° C during these measurements. We also observed no changes in NMR spectra of hydrated MBPA-C-melittin-GVA/AAO20 over more than 5 weeks of measurements; (iv) The high porosity of AAO wafers allows the preparation of samples in which the quantity of nanopore-confined peptide molecules is sufficient for a variety of ^13^C-detected NMR measurements, including 2D spectroscopy and spin relaxation measurements, with good signal-to-noise ratios for individual ^13^C-labeled sites. We note that AAO has been used in previous NMR studies of membranes and membrane proteins, without covalent tethering to pore surfaces.^71–72^

Measurements on MBPA-C-FG30/AAO20 lead to the following conclusions regarding the behavior of the nanopore-confined FG-repeat peptide: (i) In the hydrated state, ^13^C and ^1^H NMR chemical shifts indicate that FG30 has a random-coil conformational distribution at residues distributed over its sequence. The absence of temperature-dependence for chemical shifts of most sites rules out a significant change in conformational preferences from −5° C to 50° C; (ii) Spin relaxation data for ^13^C-labeled sites show smooth dependences of relaxation times on temperature. These measurements rule out a phase transition, for example between a state in which tethered FG30 molecules within a nanopore move independently of one another and a state in which groups of molecules move together as a cohesive cluster, since such a transition would produce a discontinuous change in the amplitudes or time scales of molecular motions that drive spin relaxation; (iii) Spin relaxation data indicate multiple modes of molecular motion for tethered FG30 molecules at high concentrations in AAO20 nanopores, including motions on the ~1 ns time scale that randomize bond directions over a range of approximately ±55° and motions on the ~100 ns time scale that cover a larger range of angles. T_2_^H^ measurements at low MAS frequencies indicate nearly complete isotropy of motional averaging. Interestingly, motional parameters derived from the relaxation data are similar for all isotopically labeled residues, implying approximately uniform motion from residue 6 to residue 28 in FG30; (iv) In stark contrast to its behavior when tethered in nanopores, MPBA-C-FG30 aggregates to form fibrillar assemblies in free solution at concentrations above 2 mM at 24° C or 40° C (above 5 mM at 4° C), using the same buffer conditions in nanopores or free solution. Thus, tethering to nanopore walls strongly inhibits aggregation of this FG-repeat peptide.

A likely explanation for inhibition of aggregation is that tethering to nanopore walls interferes with the close association of multiple peptide chains required for aggregation. At 90 mM concentration in a 20-nm-diameter pore, the average nearest-neighbor distance between phosphonate groups of tethered MBPA-C-FG30 molecules is approximately 1.9 nm. The length of a fully extended molecule is approximately 10 nm. These numbers imply that contacts between neighboring MBPA-C-FG30 molecules certainly exist within AAO20 nanopores. Therefore, it is conceivable that groups of MBPA-C-FG30 molecules could associate into stable or long-lived oligomeric assemblies within which molecular motions are restricted and conformations are not random-coil-like. Our finding that formation of such assemblies does not occur in MBPA-C-FG30/AAO20 suggests that aggregation of an FG-repeat sequence requires multiple molecules to interact simultaneously through identical sets of intermolecular contacts, similar to the formation of ordered cross-β structures in amyloid fibrils by intrinsically disordered polypeptides.^28,44^ Tethering to nanopore walls may limit the number of molecules that can coalesce into an ordered assembly with sufficient size to be stable.

In the case of hydrated MBPA-C-melittin/AAO20, data in Fig. 8 show that tethering to nanopore walls at high concentrations produces a state in which the majority of peptide chains have restricted motions and α-helical conformations. Thus, effects of tethering and nanopore confinement are clearly sequence-dependent for peptides of similar length. In free solution near neutral pH and ambient temperatures, melittin is known to form tetrameric, α-helical assemblies in which adjacent melittin helices have an antiparallel alignment.^68,70^ At low pH or elevated temperatures, melittin is random-coil-like and monomeric.^69–70,73^ Since all MBPA-C-melittin molecules in our experiments are tethered to the nanopore walls at their N-termini, antiparallel alignment of neighboring molecules is not possible. Thus, intermolecular interactions in MBPA-C-melittin/AAO20 must be different from intermolecular interactions within a melittin tetramer in free solution. The observation of an α-helical melittin conformation in hydrated MBPA-C-melittin/AAO20 implies that intermolecular contacts promote helix formation by melittin at high concentrations, even when the known tetrameric structure is inaccessible.

Results presented above suggest many directions for future studies. To establish the relevance of our results for nanopore-confined FG30 to the properties of Nup FG-repeat domains in NPC channels, it will be important to extend these experiments to longer FG-repeat sequences, using AAO wafers with larger pore diameters. Since the FG-repeat domains within an NPC channel are segments of diverse Nups and therefore have heterogeneous sequences, it is possible that sequence heterogeneity also has an inhibitory effect on aggregation in the true biological context. Simultaneous tethering of heterogeneous FG-repeat sequences within AAO nanopores will therefore be of interest. Effects of post-translational modifications (PTMs) on FG-repeat chain dynamics within AAO nanopores will also be of interest, as PTMs are known to affect NPC function.^74–76^

Molecular dynamics simulations of tethered FG-repeat peptides within nanopores, similar to simulations reported by others,^15,77–80^ are likely to provide insights into the aggregation inhibition mechanism discussed above, allowing the density of peptide chains, the spacings between their tethering points, and other geometric factors to be varied systematically. Molecular dynamics simulations may also assist with the interpretation and modeling of NMR spin relaxation measurements. Additional spin relaxation measurements will be informative, including ^15^N relaxation measurements similar to those applied by others in studies of a variety of intrinsically disordered proteins.^81^

In addition to FG-repeat domains, the central channels of NPCs contain a high concentration of NTRs, which may affect the dynamics and the association state of the FG-repeat domains.^82^ By including NTRs in the solution used to hydrate the AAO nanopores after covalent attachment of FG-repeat sequences to the pore surfaces, it may be possible to study the effects of NTRs (or other macromolecules) on the properties of the tethered FG-repeat sequences with NMR measurements. In principle, translational diffusion within FG-repeat-filled nanopores could also be measured with pulsed field-gradient NMR methods,^83^ allowing dependences of diffusivity on protein sequence, temperature, and other factors to be characterized.

In conclusion, we have introduced an experimental approach for studying properties of polypeptides that are tethered to walls of nanopores with NMR. Measurements on a 30-residue FG-repeat peptide show that tethering within AAO nanopores through phosphonate-surface bonds results in samples that are stable for months or longer, that 90 mM (300 mg/ml) peptide concentrations within nanopores can be achieved, that the conformational and dynamical properties of tethered peptides can be probed with a variety of NMR techniques, and that tethering to nanopore walls strongly inhibits aggregation of the FG-repeat peptide at high concentrations.

## SUPPORTING INFORMATION

The Supporting Information is available free of charge at XXXXX: LC-MS data; NMR pulse sequences; SEM and AFM images of AAO20; ^31^C and ^31^P NMR spectra of lyophilized MBPA-C-FG30-LSAFTG; effects of H_2_SO_4_ treatment on ^31^P NMR spectra of AAO20; pH-dependent effects of incubation on AAO20 wafers; ^31^C and ^31^P NMR spectra of samples prepared with various incubation conditions; ^13^C and ^31^P NMR spectra used to calibrate signal areas; temperature-dependent 1D and 2D NMR spectra of hydrated MBPA-C-FG30-LSAFTG/AAO20; full sets of spin relaxation data and temperature-dependent relaxation times; experimental and simulated T_2_ ^H^ data at low MAS frequencies; ^31^P spin-lattice relaxation measurements; tables of MBPA-C-FG30 loading in AAO20; tables of nuclear spin relaxation times; tables of motional parameters from analyses of the relaxation times.

### AUTHOR CONTRIBUTIONS

All authors have given approval to the final version of the manuscript. All authors prepared samples, performed measurements, analyzed data, and contributed to manuscript preparation.

### NOTES

The authors declare no competing financial interest.

## ACKNOWLEDGEMENTS

This research was supported by the Intramural Research Program of the National Institute of Diabetes and Digestive and Kidney Diseases (NIDDK) within the National Institutes of Health (NIH). The contributions of the NIH authors are considered Works of the United States Government. The findings and conclusions presented in this paper are those of the authors and do not necessarily reflect the views of the NIH or the U.S. Department of Health and Human Services. We thank Jeff Reece for assistance with confocal fluorescence microscopy, the NIDDK Advanced Light Microscopy and Image Analysis Core for use of its confocal microscope, Dr. Zhuang Wei for assistance with scanning electron microscopy, and the Trans-NIH Shared Resource on Biomedical Engineering and Physical Science for use of its scanning electron microscope. We thank Dr. Ralf G. Nuzzo for suggesting phosphonate/aluminum oxide chemistry as a method for attaching polypeptides within AAO nanopores. This work was supported by the National Institute of Diabetes and Digestive and Kidney Diseases under project number 1-ZIA-DK075031.

## Notes

### Competing Interest Statement

The authors have declared no competing interest.

### Summary of Updates

Two figures and one table moved from main manuscript to supporting information. One additional figure in supporting information, showing new data. Additional analyses of nuclear spin relaxation data, described in main manuscript and new tables in supporting information. Multiple simplifications and clarifications in the main text. No changes in conclusions.

