## supporting figures and tables for "Effects of Nanopore Confinement on the Conformational, Dynamical, and Self-Assembly Properties of an FG-Repeat Peptide"

Wancheng Zhao, Wai-Ming Yau, and Robert Tycko
Laboratory of Chemical Physics
National Institute of Diabetes and Digestive and Kidney Diseases
National Institutes of Health
Bethesda, MD 20892-0520

Figures S1-S23, pages S2-S24

Tables S1-S8, pages S26-S33

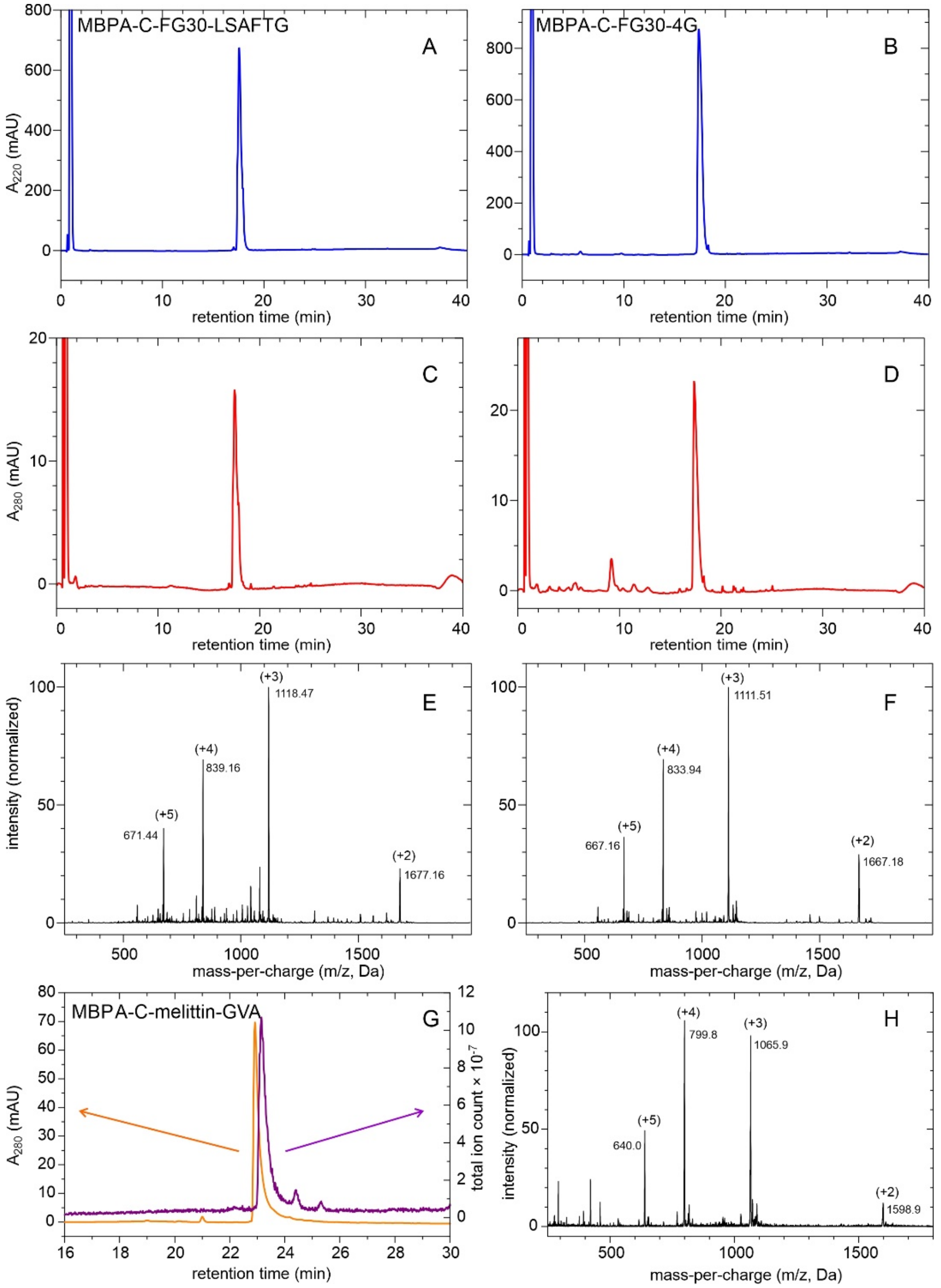


Figure S1: LC-MS data for MBPA-C-FG30-LSAFTG (A,C,E), MBPA-C-FG30-4G (B,D,F), and MBPA-C-melittin-GVA (G,H). Panels A and B show ultraviolet absorption chromatograms at 220 nm. Panels C and D show chromatograms at 280 nm. Panel G shows 280 nm absorption and ion count chromatograms. Panels E, F, and H show electrospray ionization mass spectra for the absorption chromatogram peaks at 17.6 min (E,F) or 30.0 min (H). Mass peaks for ions with charges from +2 to +5 indicate molecular masses of 3352.4 Da for MBPA-C-FG30-LSAFTG, 3331.6 Da for MBPA-C-FG30-4G, and 3195.2 Da for MBPA-C-melittin-GVA. Theoretical values are 3351.5 Da, 3330.5 Da, and 3193.8 Da, respectively.

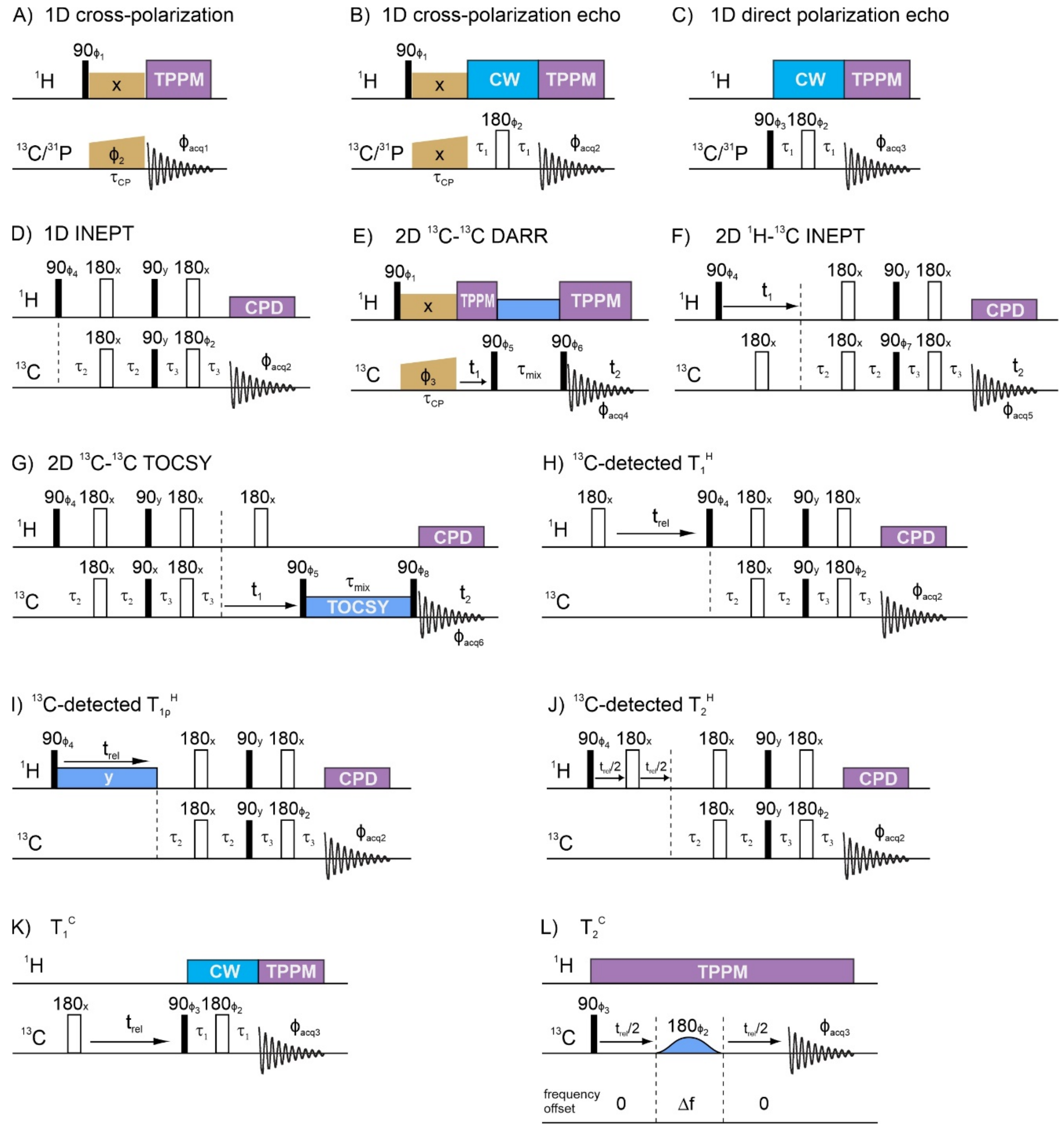


Figure S2: Pulse sequences for all NMR measurements, with TPPM representing two-pulse phase modulated decoupling, CW representing continuous-wave decoupling, and CPD representing composite pulse decoupling using a sequence of 125 phase-shifted π pulses. The same sequence of phase-shifted π pulses was used for $^{13}$C-$^{13}$C mixing in 2D total correlation spectroscopy (TOCSY). Delays in $^{1}$H-$^{13}$C INEPT polarization transfers were $\tau_2$ = 1.5 ms and $\tau_3$ = 0.9 ms. Spin echo delays were $\tau_1$ = 78.5 μs for $^{13}$C spectra and $\tau_1$ =76.3 μs for $^{31}$P spectra. Mixing periods were $\tau_{mix}$ = 30 ms in 2D $^{13}$C-$^{13}$C DARR measurements and $\tau_{mix}$ = 12.5 ms in 2D $^{13}$C-$^{13}$C TOCSY measurements. Frequency offsets Δf were adjusted to select signals of interest in $T_2^C$ measurements, with a selective π pulse length of 2.35 ms (approximate ±10 ppm bandwidth). Nonselective $\pi$ and $\pi/2$ pulse lengths were 8.6 μs and 4.3 μs for $^{1}H$, 5.0 μs and 10.0 μs for $^{13}C$, and 13.0 μs and 6.5 μs for $^{31}P$. Cross-polarization contact times $\tau_{CP}$ were optimized for each sample and were typically 1.2-1.7 ms for $^{1}H$-$^{13}C$ and 0.7-1.1 ms for $^{1}H$-$^{31}P$. Phase cycling was implemented as follows: $\phi_1$ = {y,-y}; $\phi_2$ = {x,x,y,y,-x,-x,-y,-y}; $\phi_3$ = {x,y}; $\phi_4$ = {x,-x}; $\phi_5$ = {y,y,-y,-y}; $\phi_6$ = {x,x,x,x,y,y,y,y}; $\phi_7$ = {x,x,-x,-x,y,y,-y,-y}; $\phi_8$ = {x,x,x,x,y,y,y,y,-x,-x,-x,-x,-y,-y,-y,-y}; $\phi_{acq1}$ = {x,-x,y,-y,-x,x,-y,y}; $\phi_{acq2}$ = {x,-x,-x,x,x,-x,-x,x}; $\phi_{acq3}$ = {y,x,-y,-x,y,x,-y,-x}; $\phi_{acq4}$ = {x,-x,-x,x,y,-y,-y,y}; $\phi_{acq5}$ = {x,-x,-x,x,-y,y,y,-y}; $\phi_{acq6}$ = {x,-x,-x,x,y,-y,-y,y,-x,x,x,-x,-y,y,y,-y}.

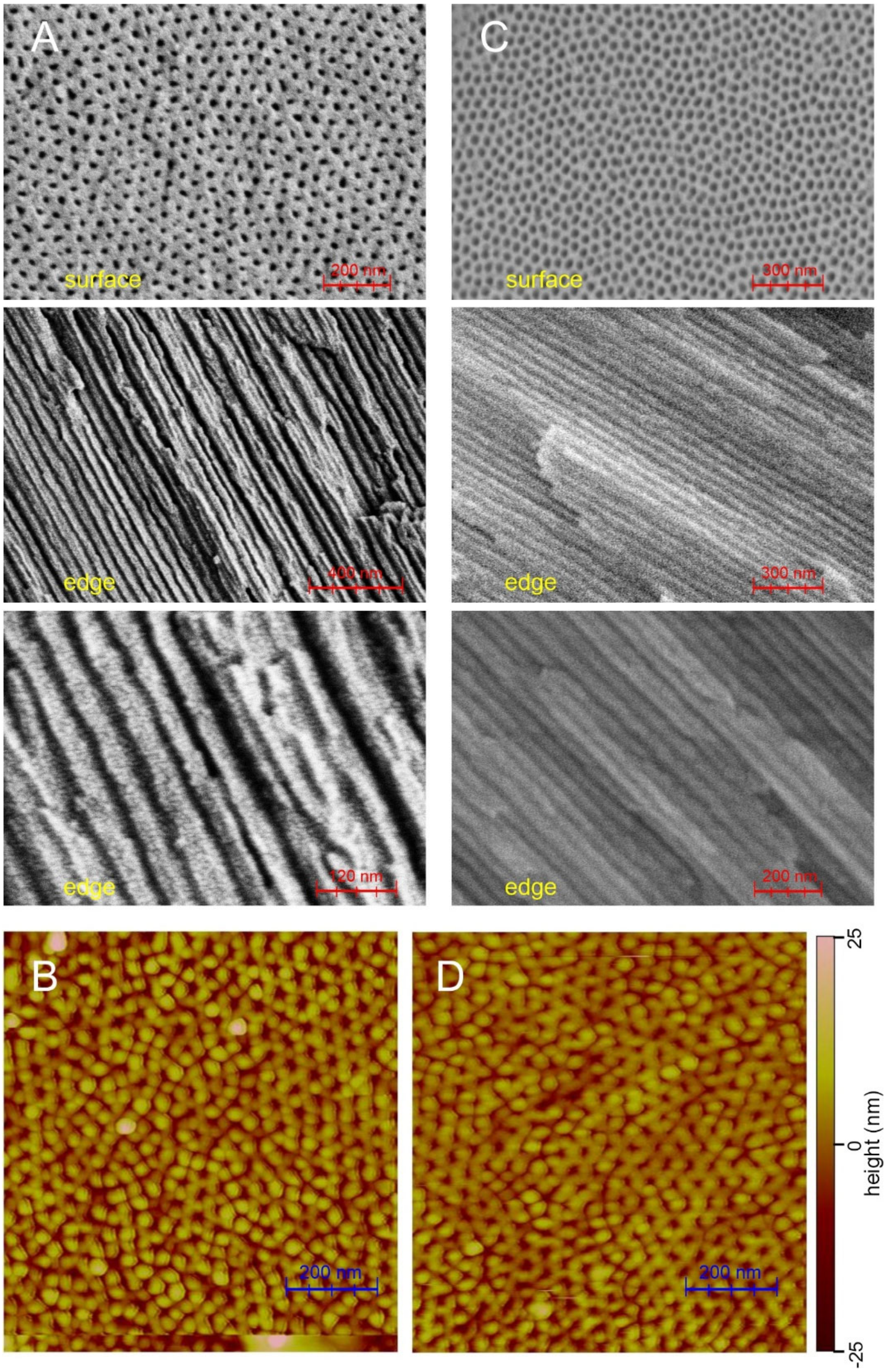


Figure S3: (A) SEM images of an untreated AAO20 wafer. Upper image shows the wafer surface. Images below show edges that were exposed after breaking the wafer. (B) AFM image of the surface of an untreated AAO20 wafer. (C) SEM images of an AAO20 wafer after treatment with 0.1 M $H_2SO_4$ to remove surface phosphate, followed by incubation with 1.0 mM MBPA-C-FG30 in 0.2 M Bis-Tris buffer, pH 6.5, at 40° C for 24 h. (D) AFM image after treatment with 0.1 M $H_2SO_4$ (6 min treatment followed by $H_2O$ rinses, repeated twice as described in the text).

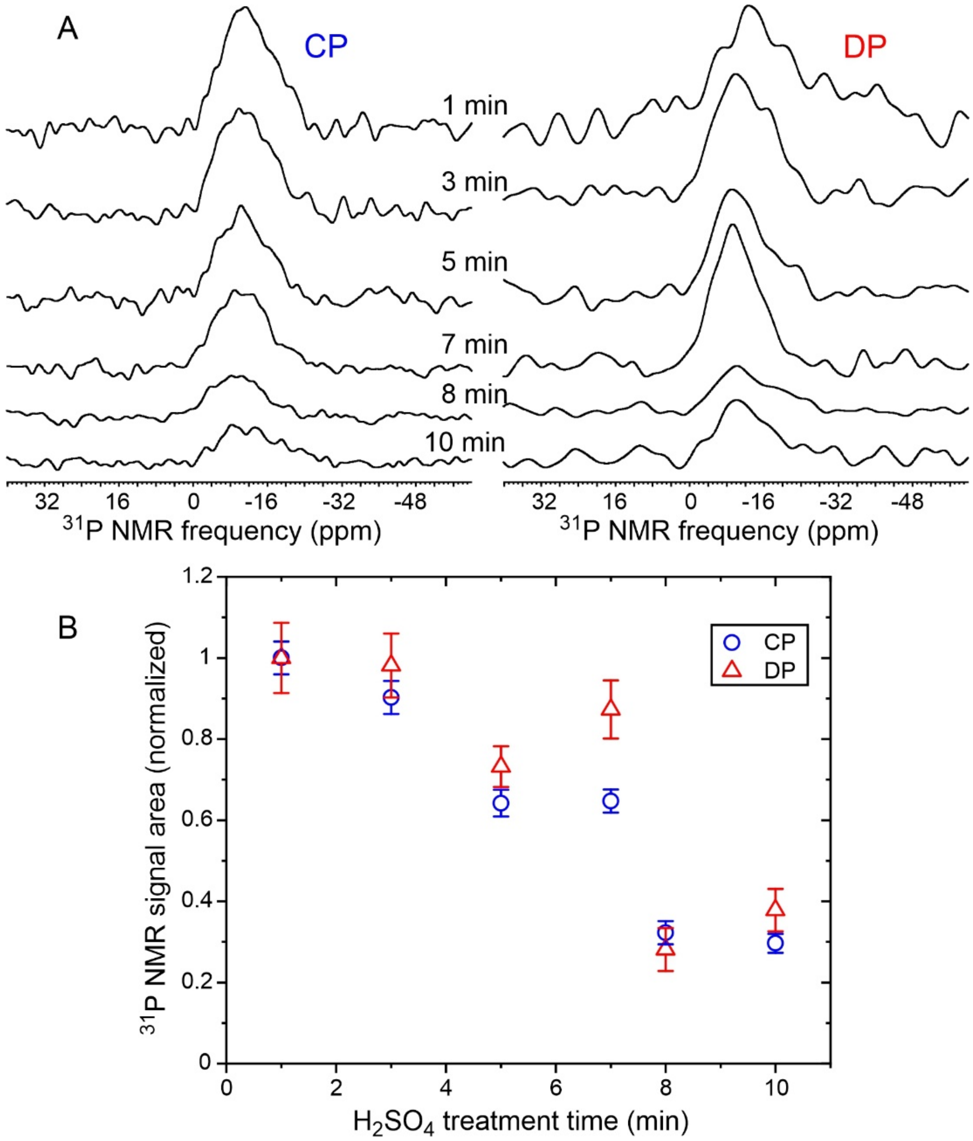


Figure S4: (A) $^{31}$P NMR spectra of AAO20 after the indicated treatment times with 0.1 M $H_2SO_4$, showing the time-dependence of signal from phosphate groups on AAO20 surfaces. Spectra were recorded with $^{1}$H-$^{31}$P cross polarization (left) or with direct pulsing (right). Vertical scales are adjusted to account for differences in sample masses and signal averaging times. (B) $^{31}$P peak areas from spectra in panel A, normalized to 1.0 at 1 min treatment time.

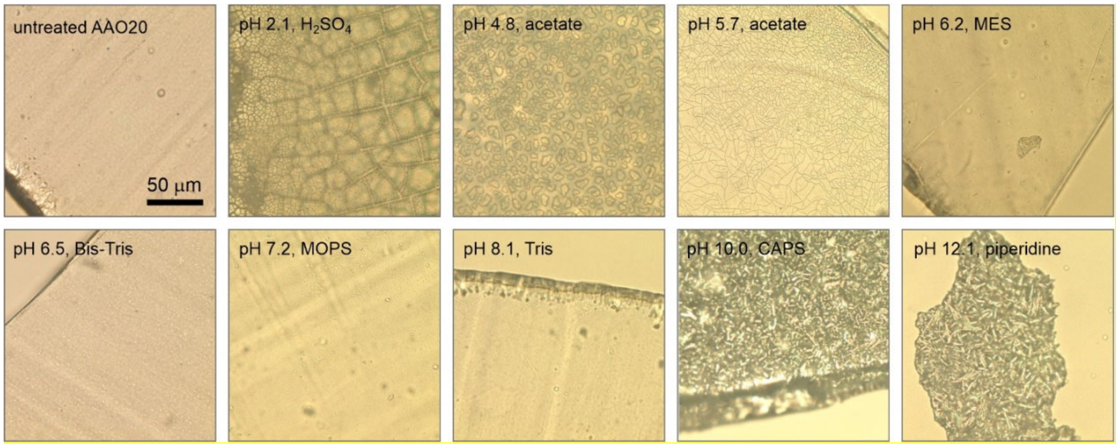


Figure S5: Optical microscope images of AAO20 wafers after incubation for 94 h at 40° C in the indicated buffers (0.2 M buffer concentration, one wafer in 1.5 ml buffer volume, continuous rotation during incubation). Images after incubation at pH values in the 6.2-8.1 range are indistinguishable from images of an untreated wafer. In contrast, incubation at pH values outside this range resulted in obvious damage to the AAO20 material. Incubation at pH 0.4 and pH 14.7 caused complete dissolution of AAO20.

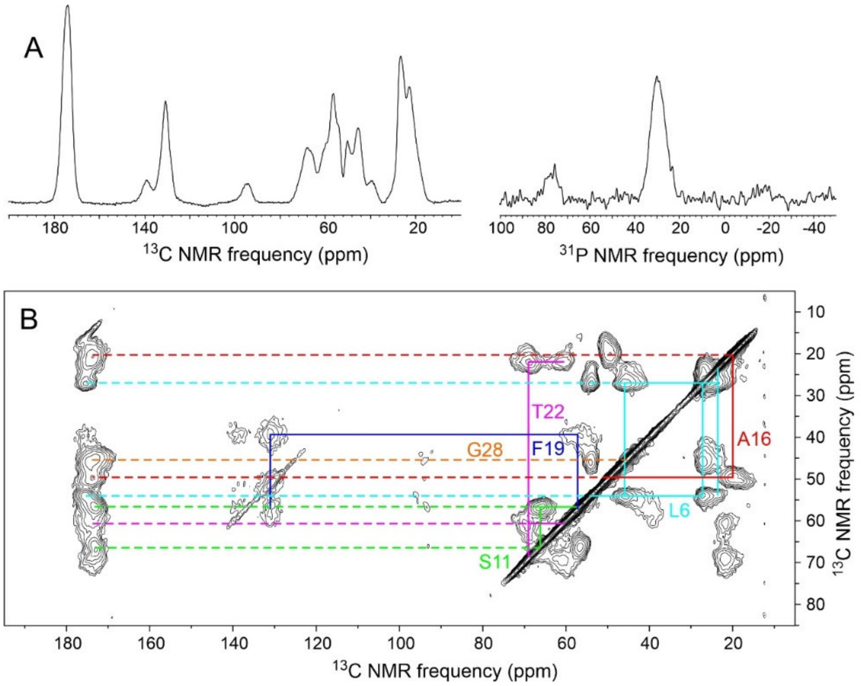


Figure S6: (A) 1D $^{13}C$ and $^{31}P$ spectra of lyophilized MBPA-C-FG30-LSAFTG, recorded in a 14.1 T field at 20° C with $^{1}H$-$^{13}C$ cross-polarization, high-power $^{1}H$ decoupling, and MAS at 12.00 kHz. The sample mass was 6.0 mg, and spectra were obtained with 512 scans and 1024 scans, respectively. (B) 2D $^{13}C$-$^{13}C$ spectrum of the same sample, recorded with a 30 ms DARR mixing period. Color-coded lines indicate assignments of crosspeaks to the six $^{15}N$,$^{13}C$-labeled residues. Contour levels increase by successive factors of 1.5.

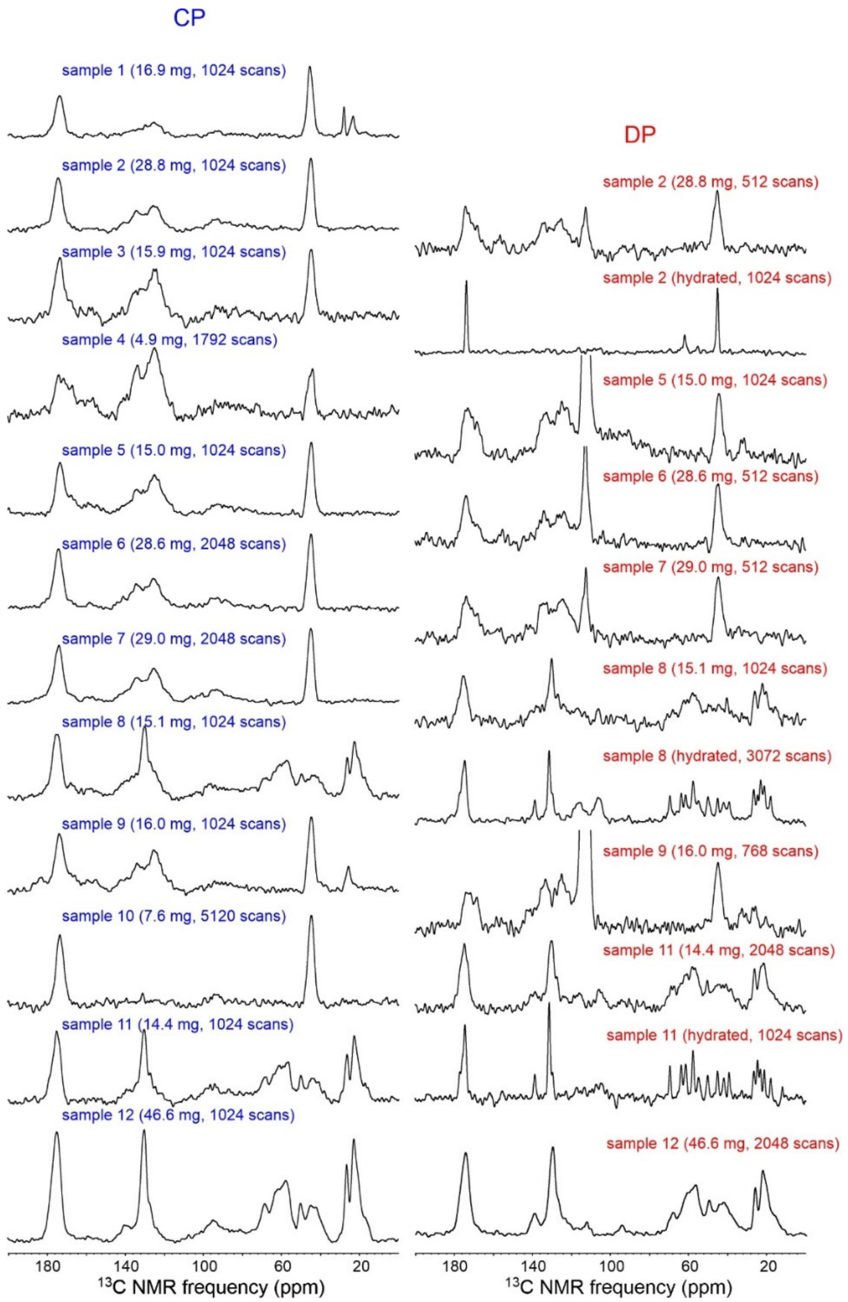


Figure S7: 1D $^{13}$C NMR spectra of MBPA-C-FG30/AAO20 samples listed in Table S1. Spectra were recorded at 14.1 T with 12.00 kHz MAS and were obtained with $^{1}$H-$^{13}$C cross-polarization (left) or direct pulsing (right). Loading values in Table S1 were determined by comparing aliphatic signal areas in these spectra with aliphatic signal areas in Fig. S9A.

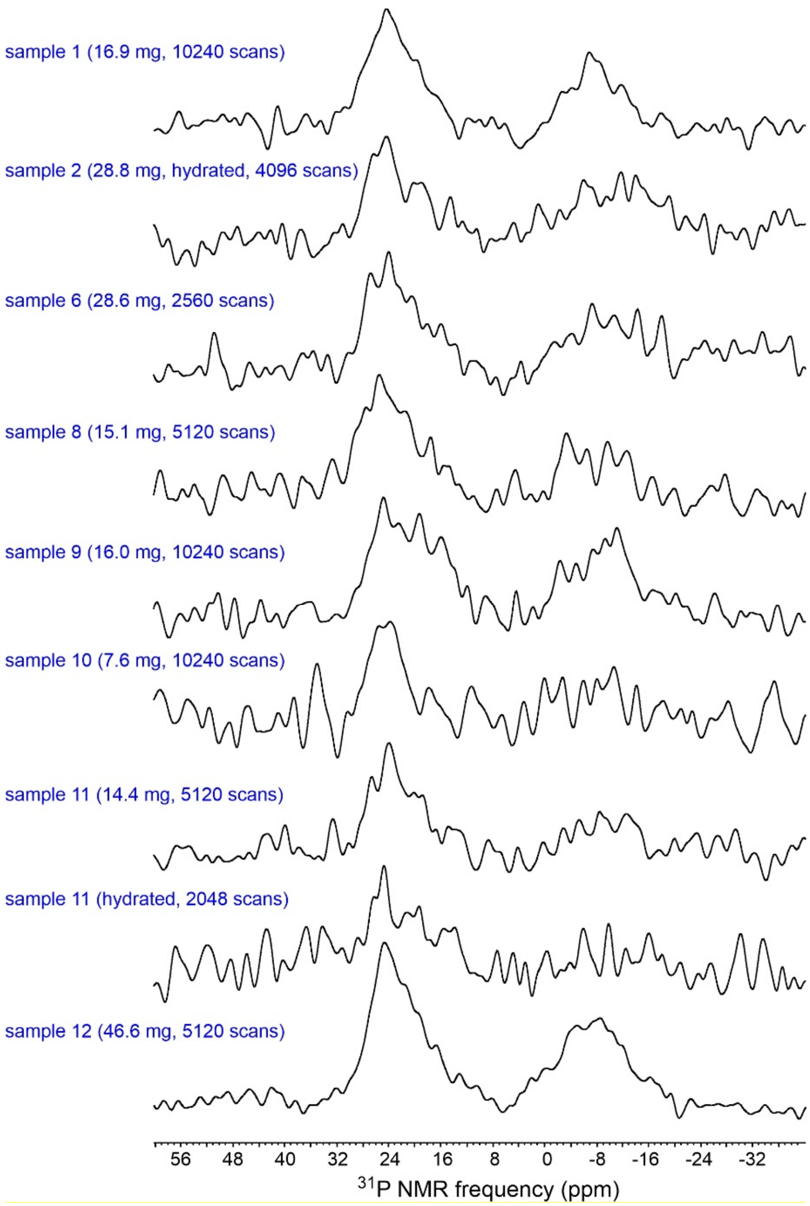


Figure S8: 1D $^{31}$P NMR spectra of MBPA-C-FG30/AAO20 samples listed in Table S1 of the main text, obtained with $^{1}$H-$^{31}$P cross-polarization. Loading values in Table S1 were determined by comparing signal areas in the 6-32 ppm range with the signal area in the cross-polarized spectrum in Fig. S9B. Signals upfield of 6 ppm arise from residual phosphate groups on AAO20 surfaces that were not fully removed by the treatment with 0.1 M $H_2SO_4$ shown in Fig. S4.

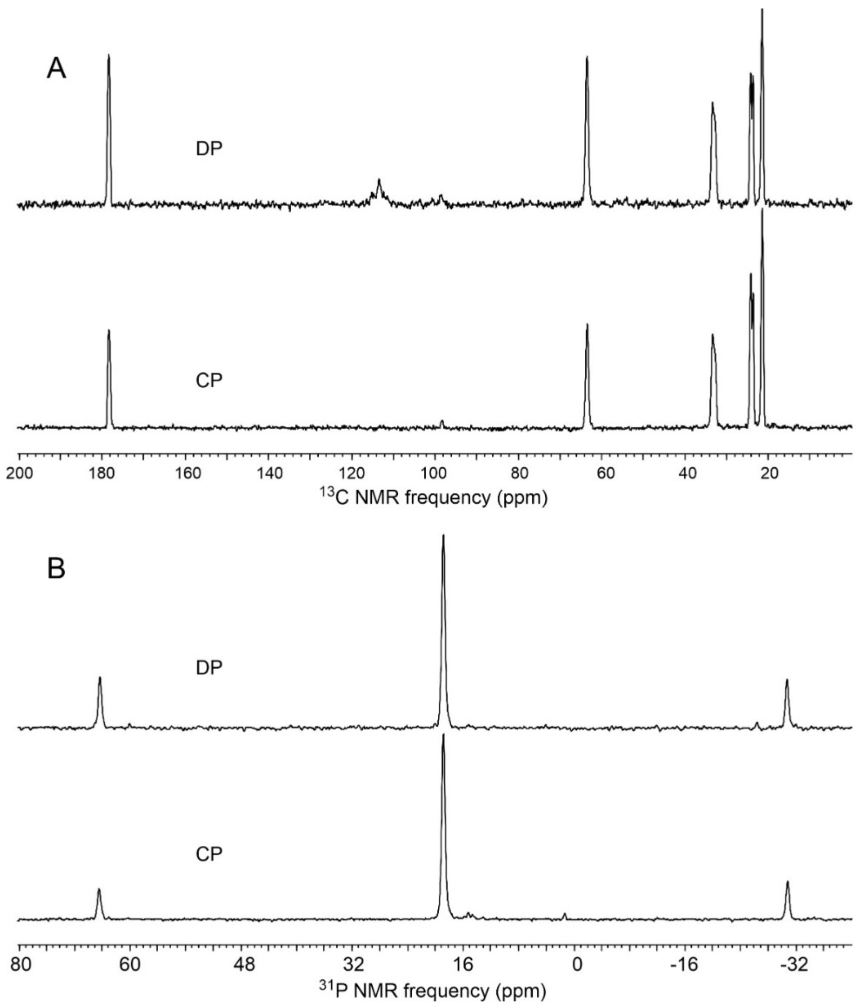


Figure S9: (A) 1D $^{13}$C NMR spectra of a 1.25 mg sample of uniformly $^{15}$N,$^{13}$C-labeled L-valine powder, obtained with direct pulsing, four scans, and a 9.0 s recycle delay (top) or with $^{1}$H-$^{13}$C cross-polarization, two scans, and 2.0 s recycle delay (bottom). (B) 1D $^{31}$P NMR spectra of a 5.0 mg sample of aminomethyl phosphonic acid (AMPA) powder, obtained with direct pulsing and one scan after a 60 min delay (top) or with $^{1}$H-$^{31}$P cross-polarization, 32 scans, and 40 s recycle delay (bottom). Spectra were recorded at 14.1 T with 12.00 kHz MAS. Signal areas in these spectra of standard samples, with known quantities of $^{13}$C and $^{31}$P, were used to calibrate areas in 1D spectra of MBPA-FG30/AAO20 samples.

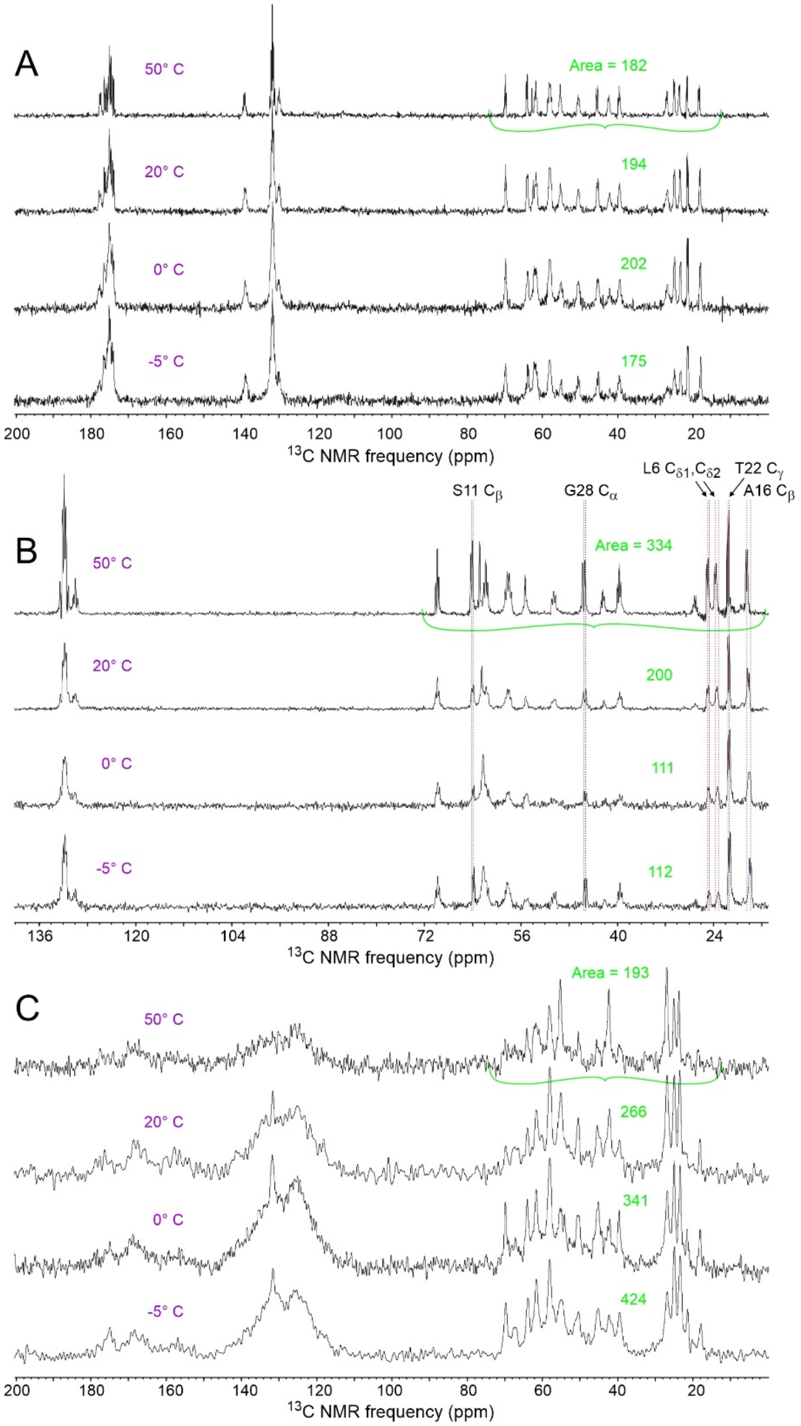


Figure S10: 1D $^{13}C$ NMR spectra of hydrated MBPA-C-FG30-LSAFTG/AAO20 obtained with direct pulsing and 1024 scans (A), $^{1}H$-$^{13}C$ INEPT and 1024 scans (B), or $^{1}H$-$^{13}C$ cross-polarization and 5120 scans (C). Spectra were recorded at the indicated sample temperatures. Areas of aliphatic signals are indicated in green. The same units are used for areas in all spectra, making them directly comparable. Uncertainties in these areas are approximately 10%. Peak positions with the strongest temperature dependences are indicated by vertical dashed lines in panel B, with blue and red lines indicating positions at -5° C and 50° C, respectively.

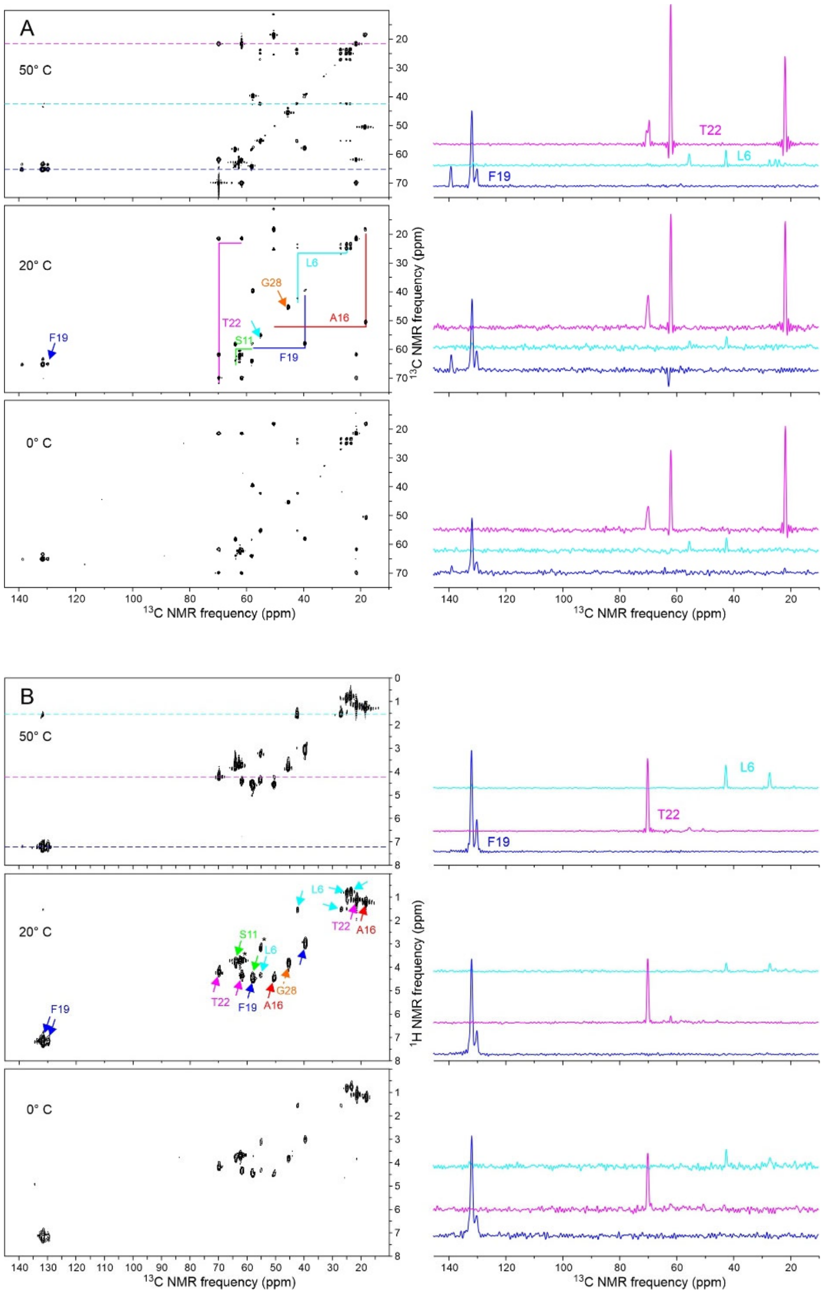


Figure S11: (A) 2D $^{13}$C-$^{13}$C NMR spectra of hydrated MBPA-C-FG30-LSAFTG/AAO20, obtained with $^{1}$H-$^{13}$C INEPT and 12.5 ms isotropic $^{13}$C-$^{13}$C mixing at the indicated sample temperatures. 1D slices, at the positions of color-coded dashed lines the 2D spectra, are shown to the right of each 2D spectrum. Spectra at 0° C, 20° C, and 50° C were recorded with 512, 832, and 864 scans per complex $t_1$ point, respectively. (B) 2D $^{1}$H-$^{13}$C NMR spectra of hydrated MBPA-C-FG30-LSAFTG/AAO20, obtained with $^{1}$H-$^{13}$C INEPT at the indicated sample temperatures. 1D slices, at the positions of color-coded dashed lines the 2D spectra, are shown to the right of each 2D spectrum. All 2D $^{1}$H-$^{13}$C NMR spectra were recorded with 512 scans per complex $t_1$ point. Contour levels increase by successive factors of 1.8 in all 2D spectra in panels A and B.

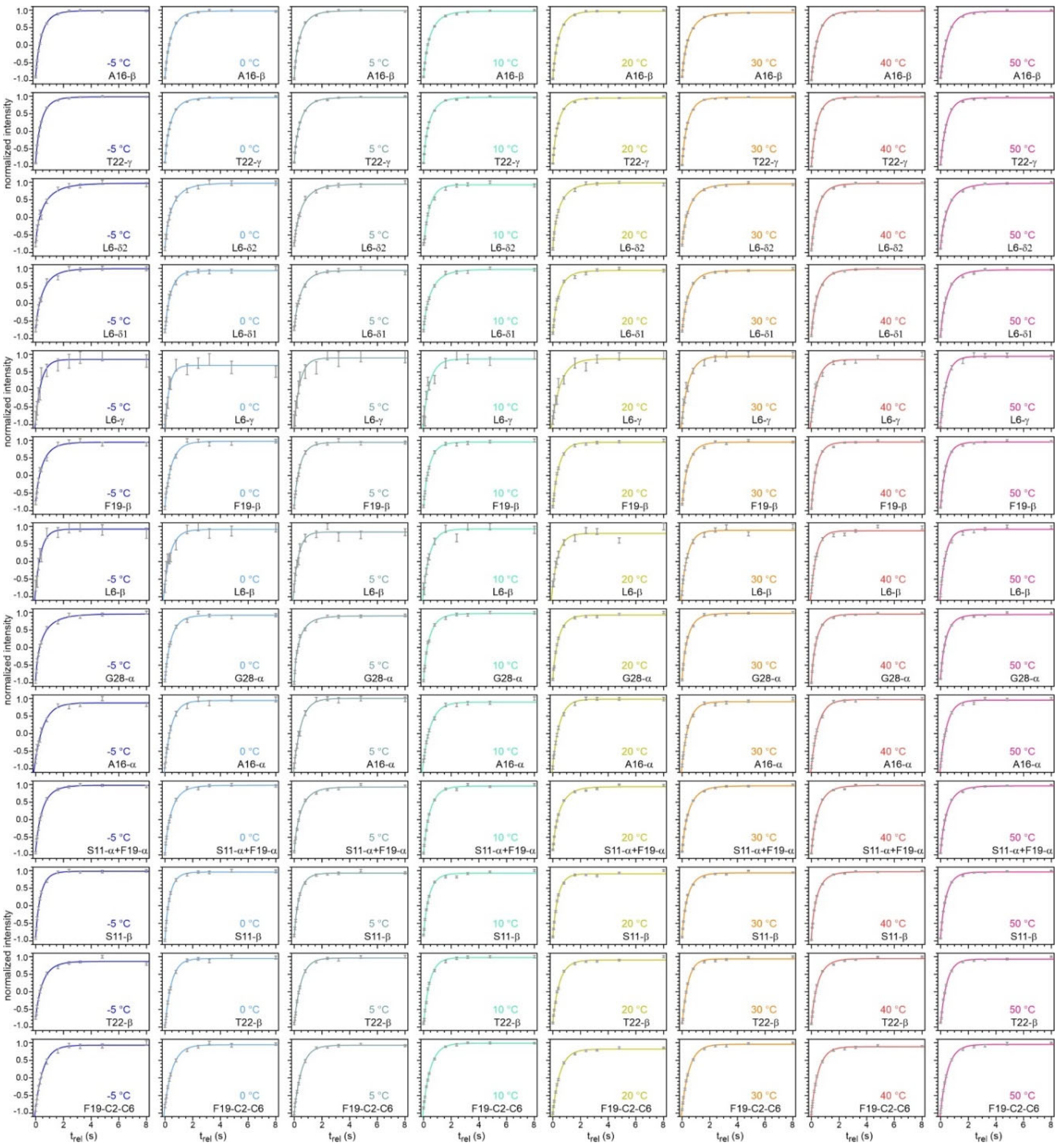


Figure S12: Full set of $T_1^H$ data for hydrated MBPA-C-FG30-LSAFTG/AAO20, obtained with the pulse sequence in Fig. S2H at sample temperatures from -5° C to 50° C. $^1$H inversion-recovery data were detected through $^{13}$C NMR signals of $^{13}$C-labeled sites. For example, data labeled T22-γ were detected through signals of the $C_\gamma$ site of T22. Signals from S11 $C_\alpha$ and F19 $C_\alpha$ were not resolved and were therefore measured as a single set of data. Signals from the $C_2$-$C_6$ aromatic sites of F19 are labeled F19-C2-6. Error bars were calculated from the RMS noise in the individual $^{13}$C NMR spectra. Solid lines are single-exponential (L6-γ, L6-β, A16-α, F19-C2-6) or stretched-exponential (all others) fits.

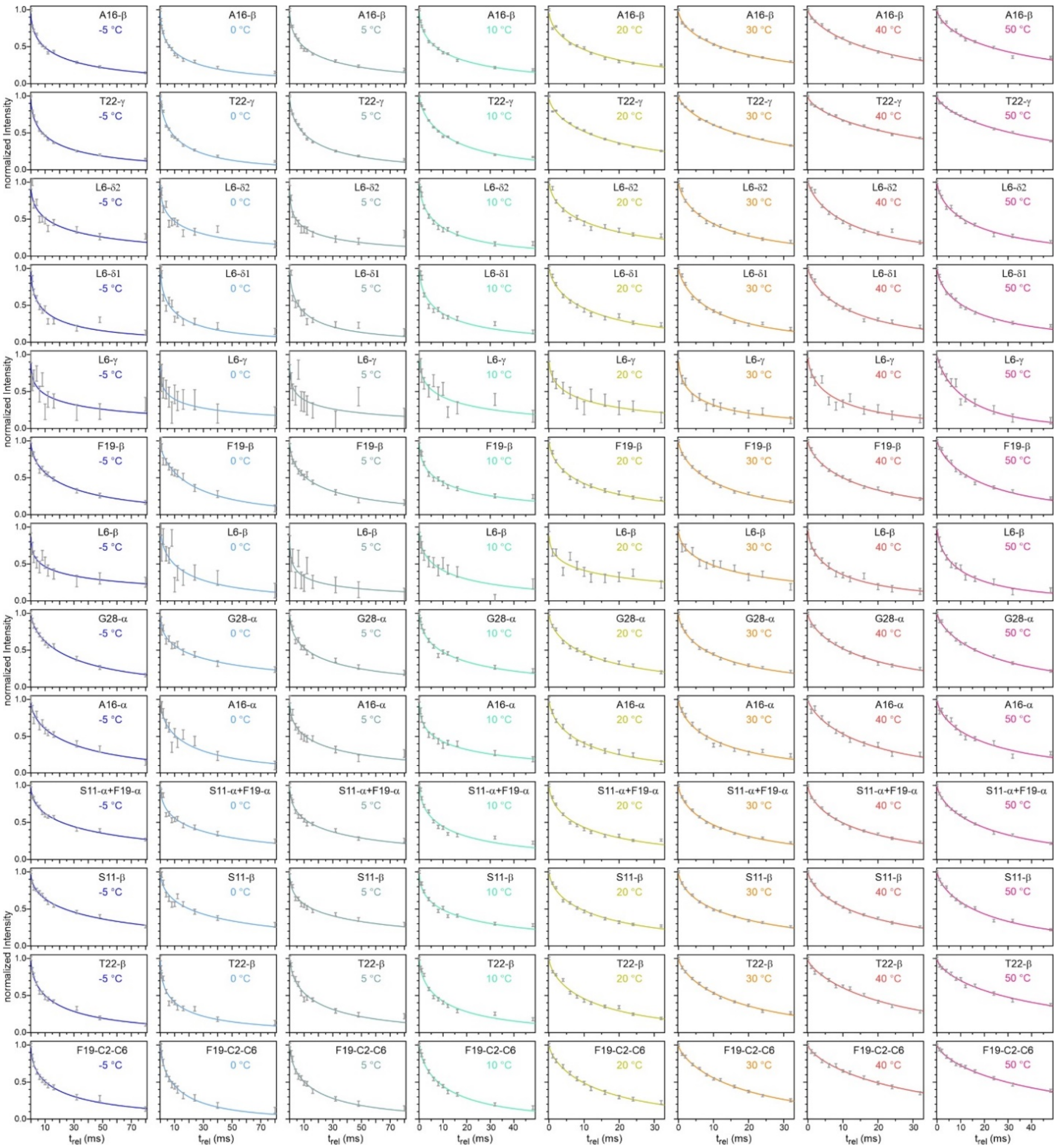


Figure S13: Full set of $T_{1\rho}^H$ data for hydrated MBPA-C-FG30-LSAFTG/AAO20, obtained with the pulse sequence in Fig. S2I and labeled as in Fig. S12. Solid lines are stretched-exponential fits.

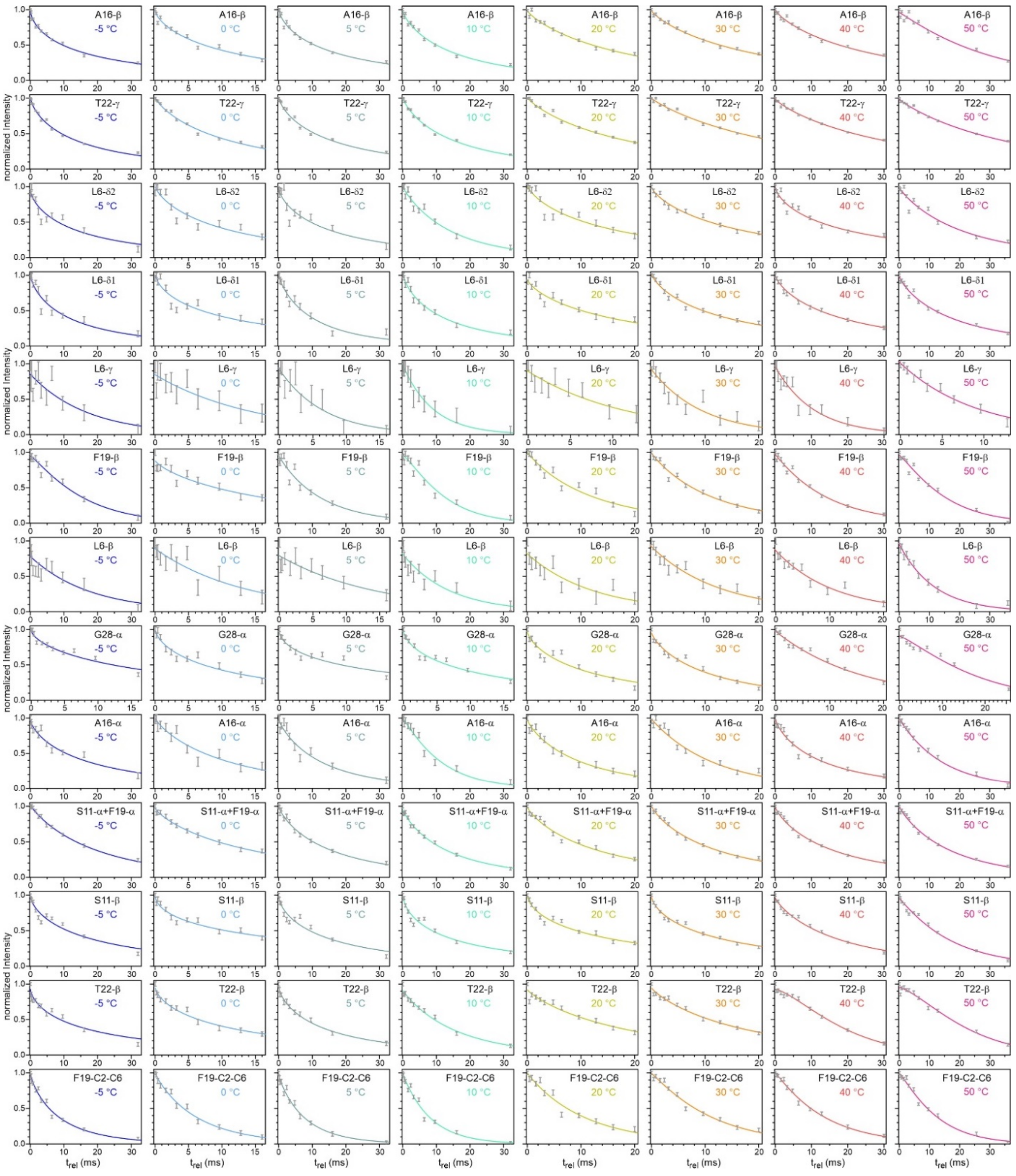


Figure S14: Full set of $T_2^H$ data for hydrated MBPA-C-FG30-LSAFTG/AAO20, obtained with the pulse sequence in Fig. S2J and labeled as in Fig. S12. Solid lines are single-exponential (L6-γ, L6-β) or stretched-exponential (all others) fits.

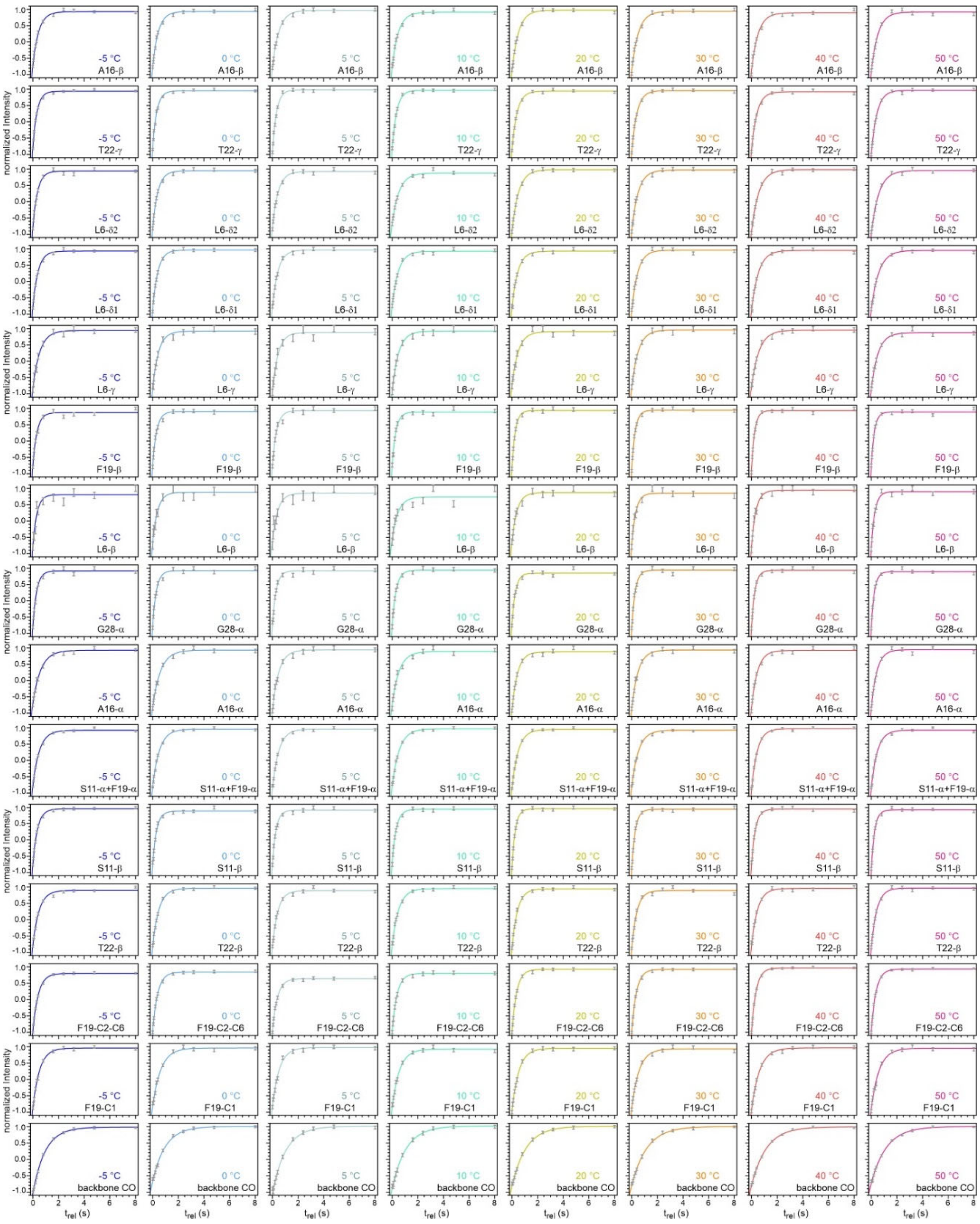

Figure S15: Full set of $T_1^C$ data for hydrated MBPA-C-FG30-LSAFTG/AAO20, obtained with the pulse sequence in Fig. S2K and labeled as in Fig. S12. Solid lines are single-exponential fits.

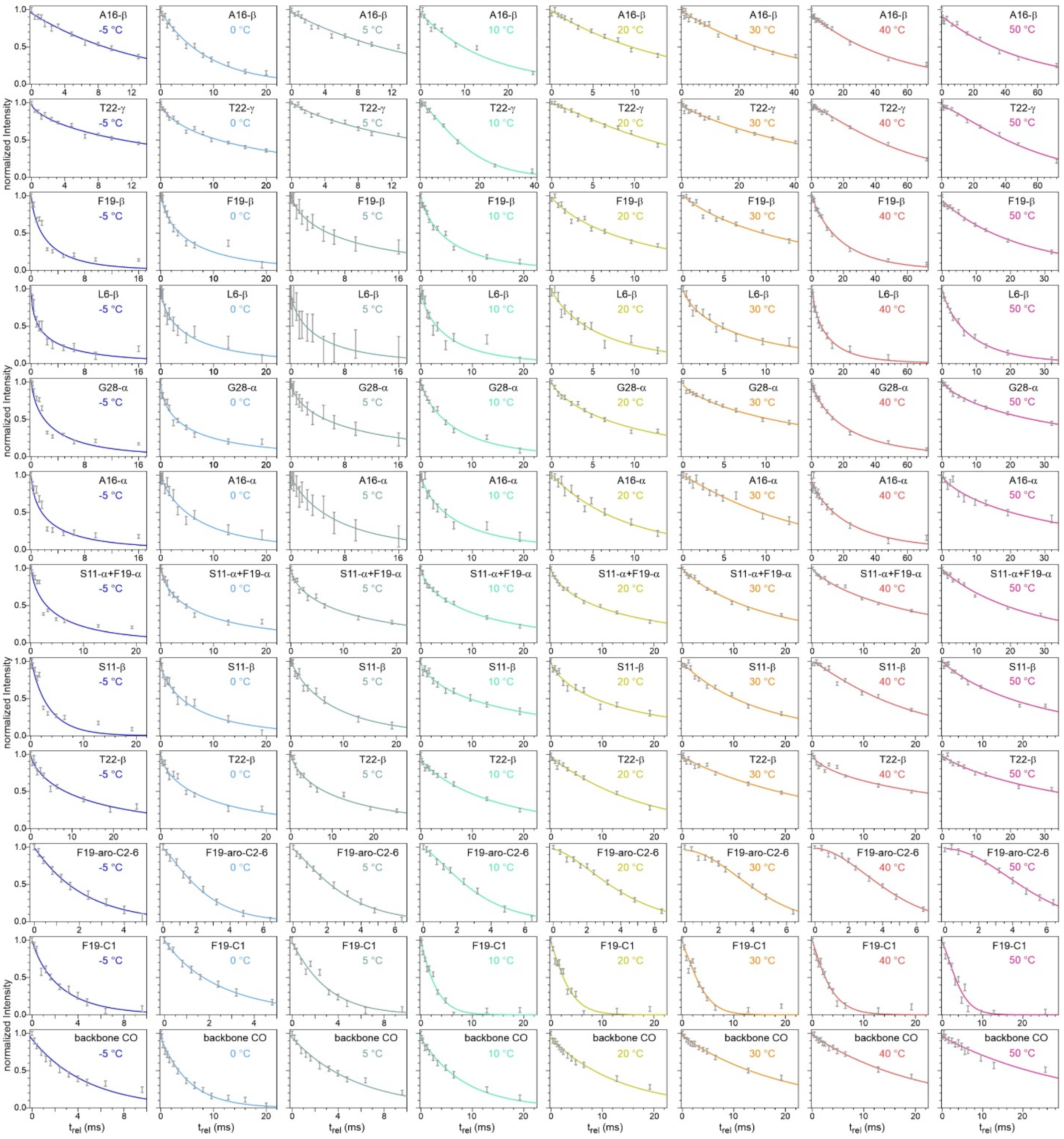


Figure S16: Full set of $T_2^C$ data for hydrated MBPA-C-FG30-LSAFTG/AAO20, obtained with the pulse sequence in Fig. S2L and labeled as in Fig. S12. Solid lines are single-exponential (A16-β, backbone CO) or stretched-exponential (all others) fits.

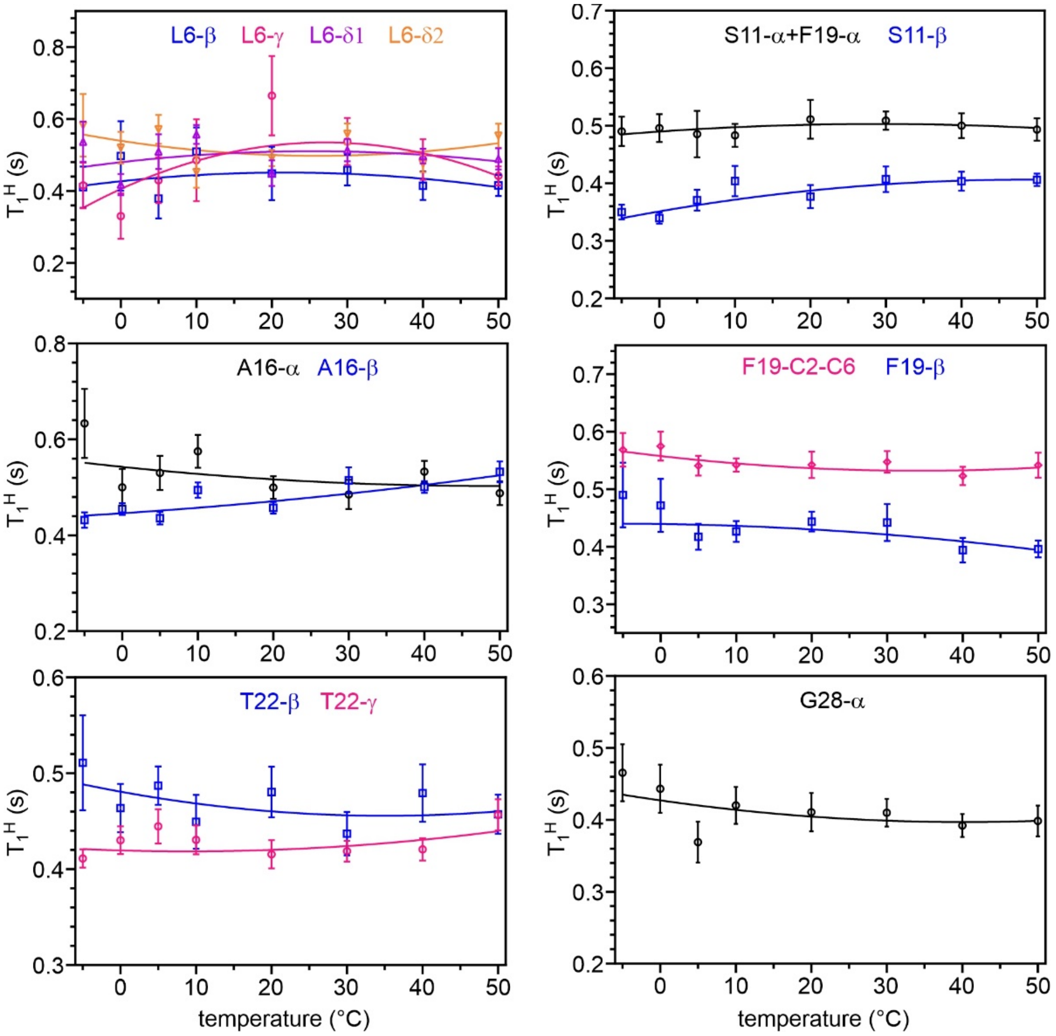


Figure S17: Temperature dependences of $T_1^H$ values determined from data in Fig. S12. Error bars are standard errors reported by the GraphPad fitting software. Solid lines are second-order polynomial functions, serving as guides to the eye.

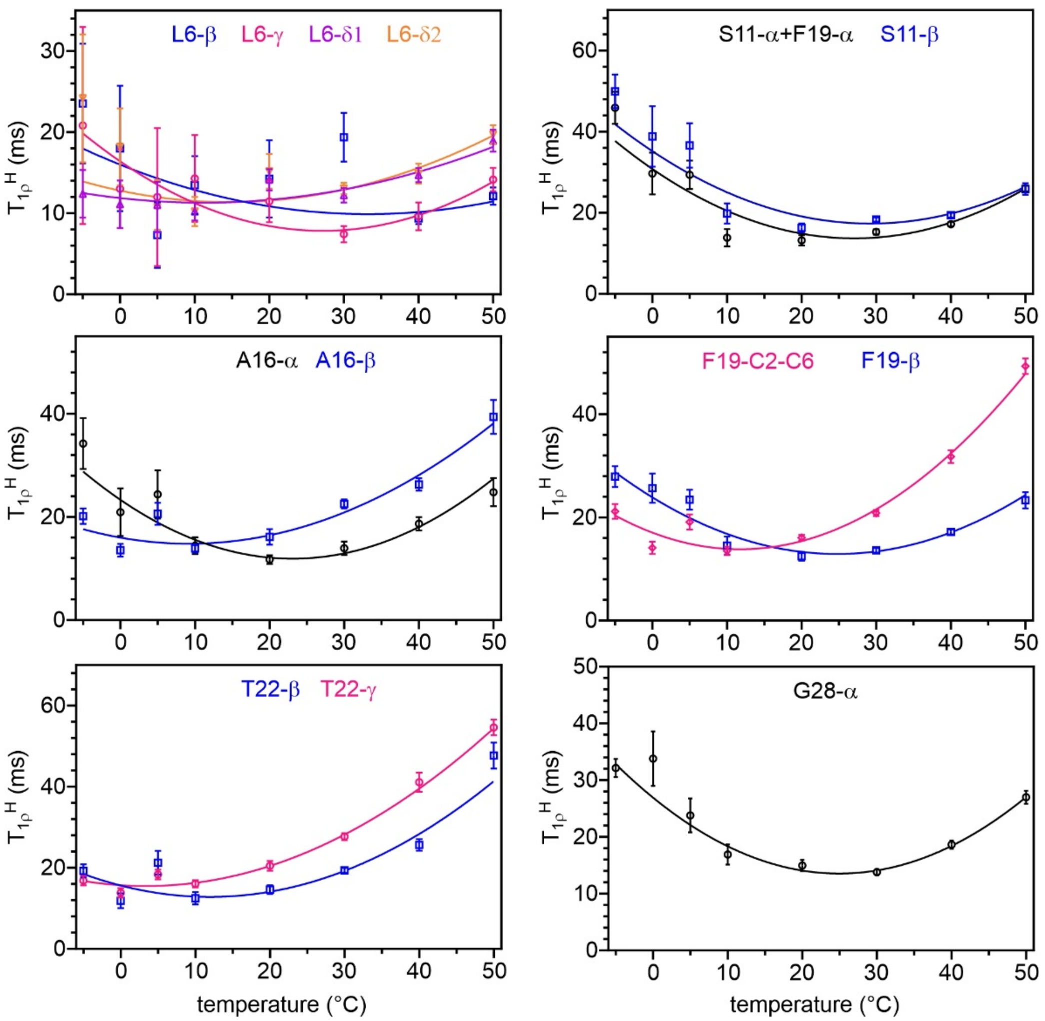


Figure S18: Temperature dependences of $T_{1\rho}^{H}$ values determined from data in Fig. S13. Error bars are standard errors reported by the GraphPad fitting software. Solid lines are second-order polynomial functions, serving as guides to the eye.

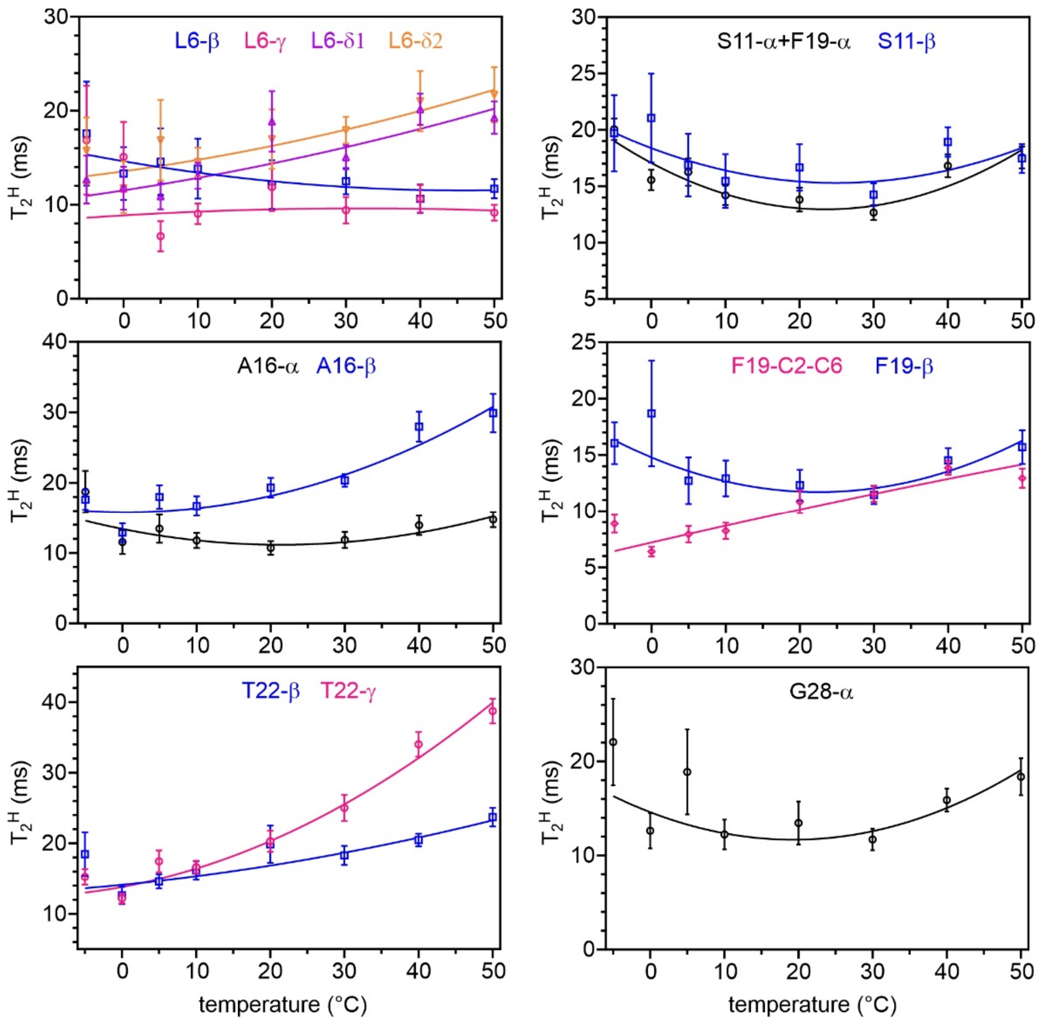


Figure S19: Temperature dependences of $T_2^H$ values determined from data in Fig. S14. Error bars are standard errors reported by the GraphPad fitting software. Solid lines are second-order polynomial functions, serving as guides to the eye.

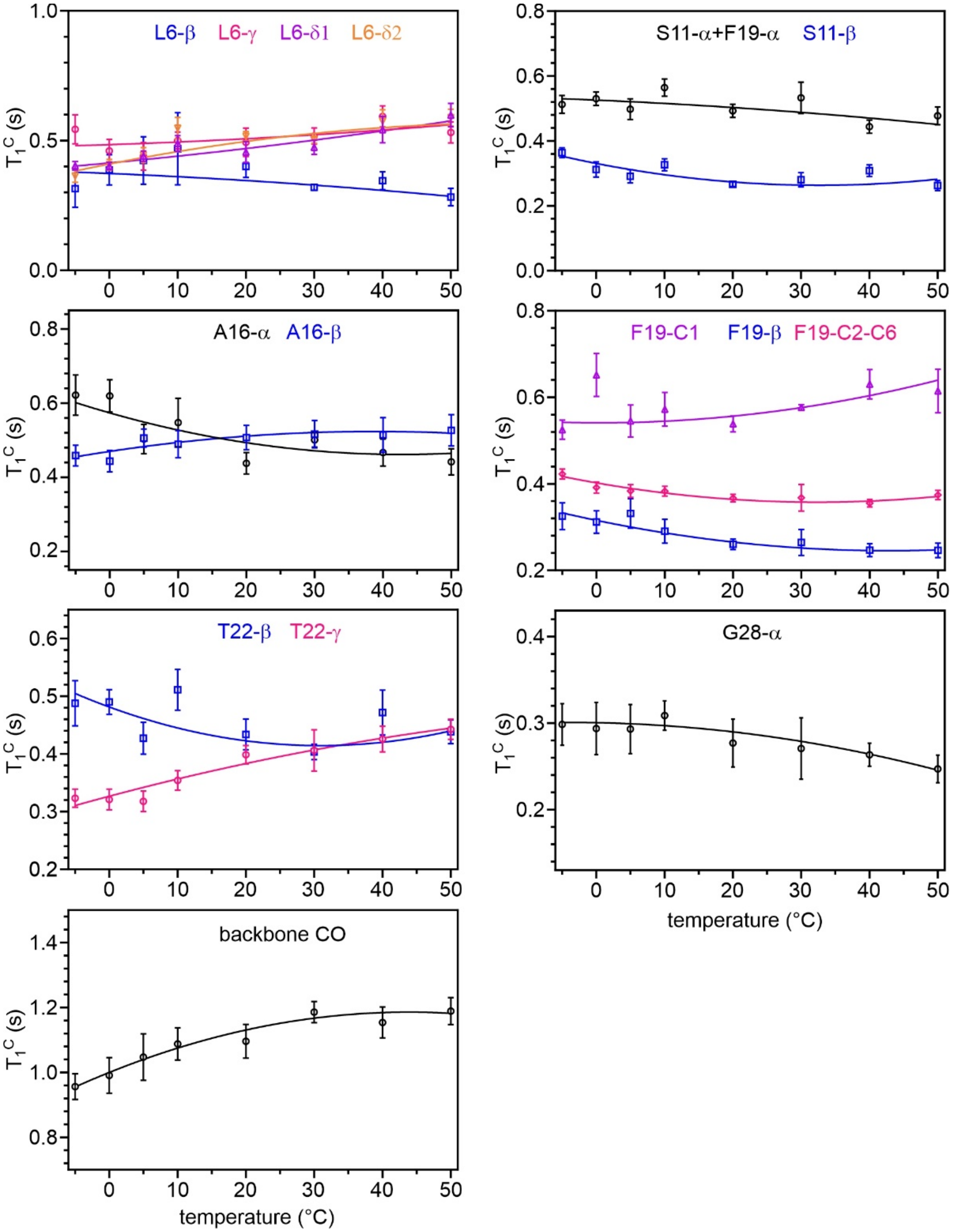


Figure S20: Temperature dependences of $T_1^C$ values determined from data in Fig. S15. Error bars are standard errors reported by the GraphPad fitting software. Solid lines are second-order polynomial functions, serving as guides to the eye.

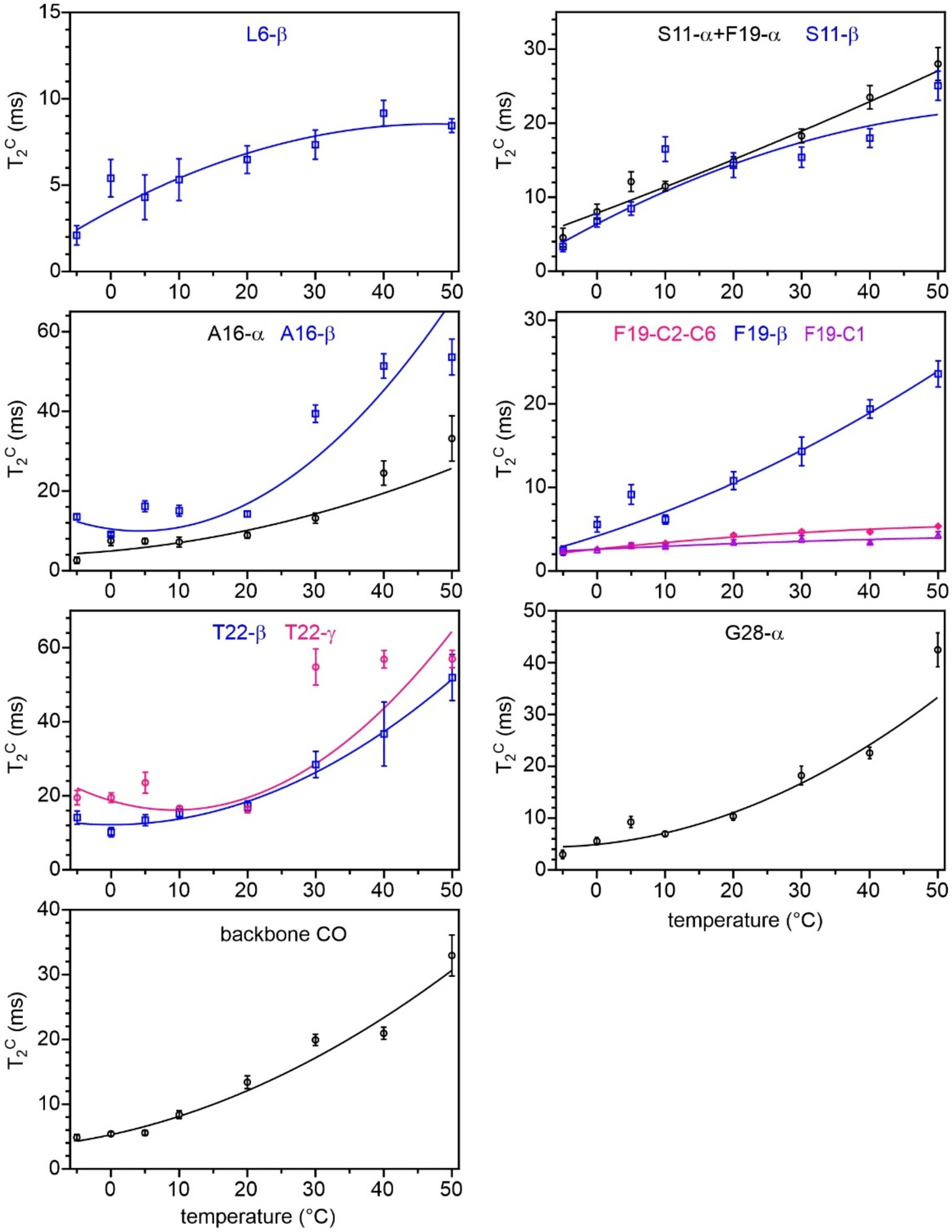


Figure S21: Temperature dependences of $T_2^C$ values determined from data in Fig. S16. Error bars are standard errors reported by the GraphPad fitting software. Solid lines are second-order polynomial functions, serving as guides to the eye.

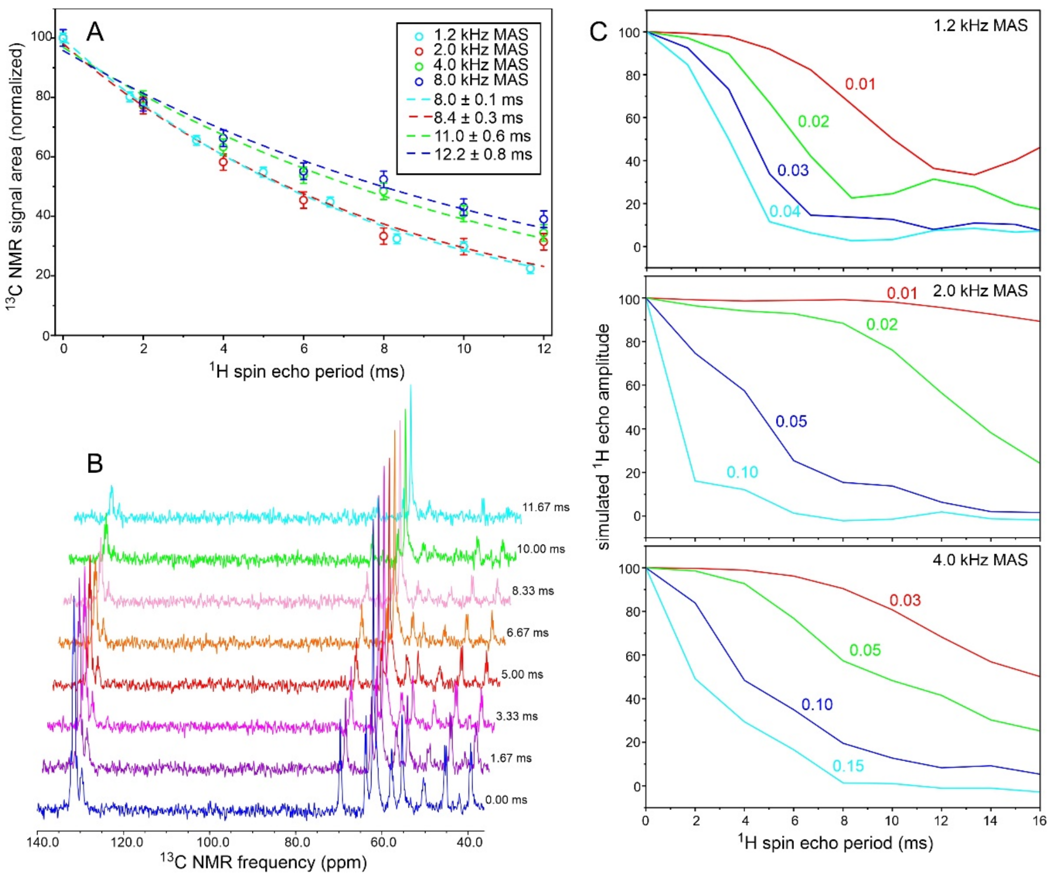


Figure S22: Test of isotropy of molecular motions in hydrated MBPA-C-FG30-LSAFTG. (A) $^{13}$C-detected measurements of $T_2^H$, using the pulse sequence in Fig. S2J, with $t_{rel}/2$ incremented in multiples of the MAS rotation period. The sample temperature was 24° C. Data are shown for MAS frequencies of 1.2 kHz, 2.0 kHz, 4.0 kHz, and 8.0 kHz (cyan, red, green, and blue symbols, respectively). Data values are signal areas in the 36-72 ppm range, normalized to initial values of 100, with error bars determined from the RMS noise in the $^{13}$C NMR spectra. Color-coded dashed lines are least-squares fits to single-exponential decays, yielding the indicated $T_2^H$ values. $T_2^H$ decreases with decreasing MAS frequency, but only by a factor of approximately 0.75. (B) $^{13}$C NMR spectra at 1.2 kHz MAS frequency with the indicated values of $t_{rel}$. (C) Simulated spin-echo decays at MAS frequencies of 1.2 kHz, 2.0 kHz, and 4.0 kHz for a system of four dipole-coupled $^1$H nuclei, with dipole-dipole couplings scaled by the indicated factors to represent partial motional averaging by rapid but anisotropic molecular motions. Nuclear coordinates (Å units) in a "molecule-fixed" axis system were set to (0.9,0.0,1.0), (-0.9,0.0,1.0), (0.0,0.9,-1.0), and (0.0,-0.9,-1.0), chosen to approximate the $^1$H positions of two neighboring methylene groups in an alkyl chain. $^1$H chemical shifts were set to 2.0, 1.0, -1.0, and -2.0 ppm (without chemical shift anisotropy). Simulations include averaging over 729 molecular orientations in the MAS rotor. Comparison of simulations at 1.2 kHz with experimental results in panel A indicates that molecular motions in MBPA-C-FG30-LSAFTG produce a scaling factor less than 0.02.

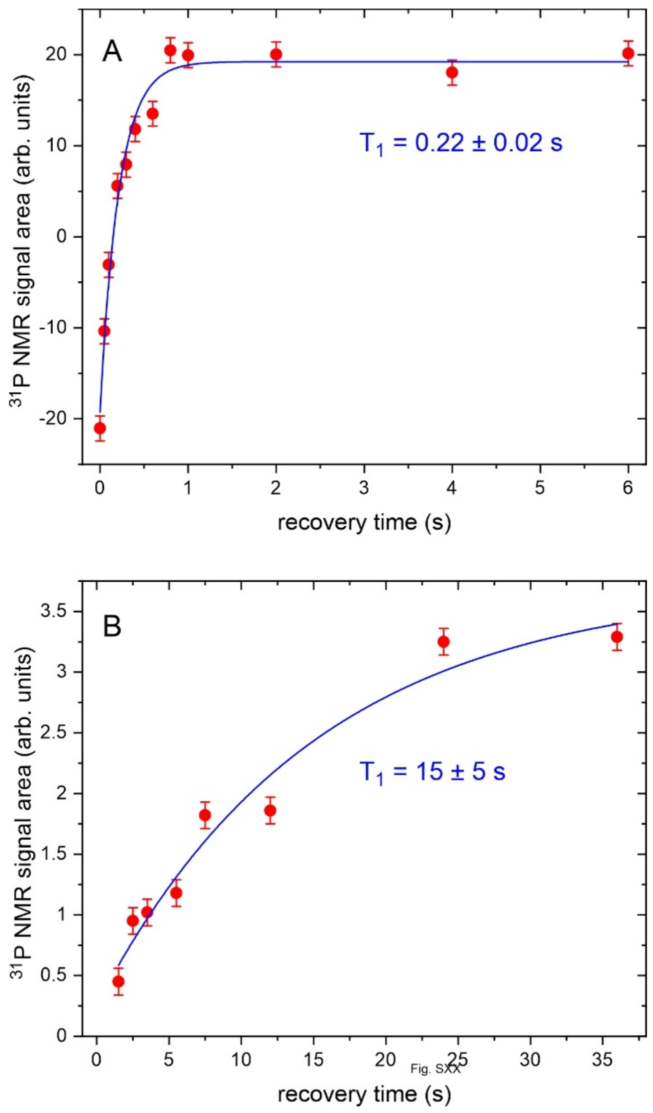


Figure S23: Measurements of $^{31}$P spin-lattice relaxation times $T_1$ in aggregated MBPA-C-FG30-LSAFTG (A) and hydrated MBPA-C-FG30-LSAFTG/AAO20 (B), using inversion-recovery and saturation-recovery methods, respectively. Error bars are uncertainties in signal areas due to noise in the $^{31}$P NMR spectra. Blue lines are least-squares fits with single-exponential functions, yielding the indicated values of $T_1$.

Table S1: Loading of MBPA-C-FG30 in AAO20 produced by various incubation conditions, as determined from signal areas in 1D $^{13}$C and $^{31}$P NMR spectra. Uncertainties were calculated from the RMS noise in the spectra.

| sample number | labeled residues in MBPA-C-FG30 | incubation time | dry mass in MAS rotor (mg) | type of NMR spectrum | total FG30 loading (µmol) | FG30 loading per milligram of AAO20 (nmol) |
|---|---|---|---|---|---|---|
| 1 | G2, G10, G20, G29 | 24 h | 16.9 | $^{13}$C CP | 0.125 ± 0.003 | 7.39 ± 0.20 |
| 1 | G2, G10, G20, G29 | 24 h | 16.9 | $^{31}$P CP | 0.082 ± 0.005 | 4.85 ± 0.30 |
| 2[a] | G2, G10, G20, G29 | 24 h | 28.8 | $^{13}$C CP | 0.131 ± 0.002 | 4.57 ± 0.07 |
| 2[a] | G2, G10, G20, G29 | 24 h | 28.8 | $^{13}$C DP | 0.145 ± 0.007 | 5.02 ± 0.25 |
| 2[a,b] | G2, G10, G20, G29 | 24 h | 28.8 | $^{13}$C DP | 0.097 ± 0.003 | 3.38 ± 0.09 |
| 2[a,b] | G2, G10, G20, G29 | 24 h | 28.8 | $^{31}$P CP | 0.098 ± 0.011 | 3.41 ± 0.39 |
| 3[c] | G2, G10, G20, G29 | 24 h | 15.9 | $^{13}$C CP | 0.057 ± 0.001 | 3.57 ± 0.08 |
| 4[b,d] | G2, G10, G20, G29 | 24 h | 4.9 | $^{13}$C CP | 0.025 ± 0.002 | 5.13 ± 0.33 |
| 5 | G2, G10, G20, G29 | 41 h | 15.0 | $^{13}$C CP | 0.105 ± 0.002 | 7.01 ± 0.18 |
| 5 | G2, G10, G20, G29 | 41 h | 15.0 | $^{13}$C DP | 0.080 ± 0.007 | 5.35 ± 0.44 |
| 6[a] | G2, G10, G20, G29 | 64 h | 28.6 | $^{13}$C CP | 0.124 ± 0.004 | 4.34 ± 0.14 |
| 6[a] | G2, G10, G20, G29 | 64 h | 28.6 | $^{13}$C DP | 0.134 ± 0.013 | 4.70 ± 0.46 |
| 6[a] | G2, G10, G20, G29 | 64 h | 28.6 | $^{31}$P CP | 0.152 ± 0.009 | 5.32 ± 0.33 |
| 7[a,e] | G2, G10, G20, G29 | 64 h | 29.0 | $^{13}$C CP | 0.128 ± 0.004 | 4.42 ± 0.14 |
| 7[a,e] | G2, G10, G20, G29 | 64 h | 29.0 | $^{13}$C DP | 0.118 ± 0.010 | 4.05 ± 0.34 |
| 8 | L6, S11, A16, F19, T22, G28 | 65 h | 15.1 | $^{13}$C CP | 0.095 ± 0.003 | 6.27 ± 0.20 |
| 8 | L6, S11, A16, F19, T22, G28 | 65 h | 15.1 | $^{13}$C DP | 0.091 ± 0.004 | 6.01 ± 0.24 |
| 8 | L6, S11, A16, F19, T22, G28 | 65 h | 15.1 | $^{31}$P CP | 0.099 ± 0.008 | 6.53 ± 0.53 |
| 8[b] | L6, S11, A16, F19, T22, G28 | 65 h | 15.1 | $^{13}$C DP | 0.089 ± 0.002 | 5.88 ± 0.12 |
| 9[f] | G2, G10, G20, G29 | 66 h | 16.0 | $^{13}$C CP | 0.094 ± 0.002 | 5.90 ± 0.14 |
| 9[f] | G2, G10, G20, G29 | 66 h | 16.0 | $^{13}$C DP | 0.109 ± 0.009 | 6.82 ± 0.58 |
| 9[f] | G2, G10, G20, G29 | 66 h | 16.0 | $^{31}$P CP | 0.077 ± 0.005 | 4.80 ± 0.34 |
| 10 | G2, G10, G20, G29 | 96 h | 7.6 | $^{13}$C CP | 0.037 ± 0.001 | 4.86 ± 0.14 |
| 10 | G2, G10, G20, G29 | 96 h | 7.6 | $^{31}$P CP | 0.034 ± 0.008 | 4.42 ± 1.00 |
| 11 | L6, S11, A16, F19, T22, G28 | 185 h | 14.4 | $^{13}$C CP | 0.090 ± 0.002 | 6.26 ± 0.17 |
| 11 | L6, S11, A16, F19, T22, G28 | 185 h | 14.4 | $^{13}$C DP | 0.093 ± 0.002 | 6.45 ± 0.14 |
| 11 | L6, S11, A16, F19, T22, G28 | 185 h | 14.4 | $^{31}$P CP | 0.080 ± 0.003 | 5.54 ± 0.24 |
| 11[b] | L6, S11, A16, F19, T22, G28 | 185 h | 14.4 | $^{13}$C DP | 0.081 ± 0.002 | 5.63 ± 0.15 |
| 11[b] | L6, S11, A16, F19, T22, G28 | 185 h | 14.4 | $^{31}$P CP | 0.097 ± 0.014 | 6.75 ± 1.00 |
| 12 | L6, S11, A16, F19, T22, G28 | 24 h | 46.6 | $^{13}$C CP | 0.266 ± 0.003 | 5.71 ± 0.07 |

| | | | | | | |
|---|---|---|---|---|---|---|
| 12 | L6, S11, A16, F19, T22, G28 | 24 h | 46.6 | $^{13}C$ DP | 0.273 ± 0.005 | 5.87 ± 0.11 |
| 12 | L6, S11, A16, F19, T22, G28 | 24 h | 46.6 | $^{31}P$ CP | 0.265 ± 0.006 | 5.68 ± 0.14 |

[a] Three AA20 wafers incubated in one tube containing 1.0 ml of 1.0 mM MBPA-C-FG30, rather than 1.00 ml of 1.0 mM MBPA-C-FG30 per wafer in separate tubes.
[b]NMR measurements performed on hydrated MBPA-C-FG30/AAO20, rather than dry MBPA-C-FG30/AAO20.
[c]AAO with 40 nm pores (AAO40) used instead of AAO20.
[d]AAO20 with 5 μm thickness, rather than 50 μm.
[e]AAO20 wafers were crushed into pieces with diameters of approximately 2 mm before incubation.
[f]1.0 mM MBPA-C-FG30 in 25 mM acetate buffer, pH 5.6, rather than 0.2 M Bis-Tris buffer, pH 6.5.

Table S2: $T_1^H$ relaxation time values (seconds) at sample temperatures from -5° C to 50° C, determined from single-exponential or stretched-exponential fits to the data in Fig. S12 and plotted in Fig. S17. For stretched-exponential fits, stretching exponents are given in parentheses. Identical values are listed for the S11 Cα and F19 Cα sites because signals from these sites were not resolved. Uncertainties are standard errors from the fits.

| residue and carbon site | | -5 °C | 0 °C | 5 °C | 10 °C | 20 °C | 30 °C | 40 °C | 50 °C |
|---|---|---|---|---|---|---|---|---|---|
| L6 | β | 0.41 ± 0.06 | 0.50 ± 0.10 | 0.38 ± 0.06 | 0.51 ± 0.07 | 0.45 ± 0.07 | 0.46 ± 0.04 | 0.41 ± 0.04 | 0.42 ± 0.03 |
| | γ | 0.42 ± 0.06 | 0.33 ± 0.06 | 0.43 ± 0.06 | 0.49 ± 0.11 | 0.66 ± 0.11 | 0.54 ± 0.07 | 0.49 ± 0.06 | 0.44 ± 0.03 |
| | $\delta_1$ | 0.54 ± 0.06<br>(0.83 ± 0.09) | 0.42 ± 0.03<br>(0.92 ± 0.08) | 0.51 ± 0.05<br>(0.86 ± 0.09) | 0.56 ± 0.03<br>(0.82 ± 0.04) | 0.45 ± 0.04<br>(0.86 ± 0.08) | 0.51 ± 0.03<br>(0.88 ± 0.06) | 0.50 ± 0.02<br>(0.85 ± 0.04) | 0.49 ± 0.03<br>(0.83 ± 0.05) |
| | $\delta_2$ | 0.58 ± 0.09<br>(0.75 ± 0.10) | 0.52 ± 0.04<br>(0.78 ± 0.06) | 0.57 ± 0.04<br>(0.82 ± 0.06) | 0.45 ± 0.04<br>(0.98 ± 0.13) | 0.49 ± 0.02<br>(0.79 ± 0.04) | 0.56 ± 0.03<br>(0.84 ± 0.05) | 0.48 ± 0.02<br>(0.90 ± 0.05) | 0.55 ± 0.03<br>(0.81 ± 0.05) |
| S11 | α | 0.49 ± 0.03<br>(0.86 ± 0.05) | 0.50 ± 0.02<br>(0.86 ± 0.05) | 0.49 ± 0.04<br>(0.83 ± 0.07) | 0.48 ± 0.02<br>(0.93 ± 0.05) | 0.51 ± 0.03<br>(0.86 ± 0.06) | 0.51 ± 0.02<br>(0.86 ± 0.03) | 0.50 ± 0.02<br>(0.88 ± 0.04) | 0.49 ± 0.02<br>(0.87 ± 0.04) |
| | β | 0.35 ± 0.01<br>(0.87 ± 0.04) | 0.34 ± 0.01<br>(0.89 ± 0.03) | 0.37 ± 0.02<br>(0.90 ± 0.06) | 0.40 ± 0.03<br>(0.91 ± 0.08) | 0.38 ± 0.02<br>(0.95 ± 0.07) | 0.41 ± 0.02<br>(0.85 ± 0.05) | 0.40 ± 0.02<br>(0.82 ± 0.04) | 0.41 ± 0.01<br>(0.90 ± 0.03) |
| A16 | α | 0.63 ± 0.07 | 0.50 ± 0.04 | 0.53 ± 0.04 | 0.57 ± 0.03 | 0.50 ± 0.02 | 0.49 ± 0.03 | 0.53 ± 0.02 | 0.49 ± 0.02 |
| | β | 0.43 ± 0.02<br>(0.86 ± 0.04) | 0.45 ± 0.01<br>(0.87 ± 0.03) | 0.44 ± 0.01<br>(0.82 ± 0.03) | 0.49 ± 0.02<br>(0.84 ± 0.03) | 0.46 ± 0.01<br>(0.89 ± 0.03) | 0.51 ± 0.03<br>(0.83 ± 0.05) | 0.50 ± 0.01<br>(0.92 ± 0.03) | 0.53 ± 0.02<br>(0.92 ± 0.04) |
| F19 | α | 0.49 ± 0.03<br>(0.86 ± 0.05) | 0.50 ± 0.02<br>(0.86 ± 0.05) | 0.49 ± 0.04<br>(0.83 ± 0.07) | 0.48 ± 0.02<br>(0.93 ± 0.05) | 0.51 ± 0.03<br>(0.86 ± 0.06) | 0.51 ± 0.02<br>(0.86 ± 0.03) | 0.50 ± 0.02<br>(0.88 ± 0.04) | 0.49 ± 0.02<br>(0.87 ± 0.04) |
| | β | 0.49 ± 0.06<br>(0.86 ± 0.11) | 0.47 ± 0.05<br>(0.91 ± 0.11) | 0.42 ± 0.02<br>(0.92 ± 0.06) | 0.43 ± 0.02<br>(0.92 ± 0.05) | 0.44 ± 0.02<br>(0.95 ± 0.05) | 0.44 ± 0.03<br>(0.90 ± 0.08) | 0.39 ± 0.02<br>(0.92 ± 0.06) | 0.40 ± 0.01<br>(0.93 ± 0.04) |
| | C2-C6 | 0.57 ± 0.03 | 0.57 ± 0.03 | 0.54 ± 0.02 | 0.54 ± 0.01 | 0.54 ± 0.02 | 0.55 ± 0.02 | 0.52 ± 0.02 | 0.54 ± 0.02 |
| T22 | β | 0.51 ± 0.05<br>(0.88 ± 0.10) | 0.46 ± 0.03<br>(1.00 ± 0.07) | 0.49 ± 0.02<br>(0.98 ± 0.05) | 0.45 ± 0.03<br>(0.98 ± 0.08) | 0.48 ± 0.03<br>(0.96 ± 0.07) | 0.44 ± 0.02<br>(1.05 ± 0.08) | 0.48 ± 0.03<br>(0.89 ± 0.07) | 0.46 ± 0.02<br>(1.00 ± 0.06) |
| | γ | 0.41 ± 0.01<br>(0.84 ± 0.02) | 0.43 ± 0.01<br>(0.82 ± 0.03) | 0.44 ± 0.02<br>(0.81 ± 0.04) | 0.43 ± 0.02<br>(0.82 ± 0.03) | 0.42 ± 0.01<br>(0.87 ± 0.04) | 0.42 ± 0.01<br>(0.88 ± 0.03) | 0.42 ± 0.01<br>(0.91 ± 0.03) | 0.46 ± 0.02<br>(0.94 ± 0.04) |
| G28 | α | 0.47 ± 0.04<br>(0.70 ± 0.05) | 0.44 ± 0.03<br>(0.89 ± 0.08) | 0.37 ± 0.03<br>(0.73 ± 0.06) | 0.42 ± 0.03<br>(0.86 ± 0.06) | 0.41 ± 0.03<br>(0.90 ± 0.07) | 0.41 ± 0.02<br>(0.86 ± 0.05) | 0.39 ± 0.02<br>(0.85 ± 0.04) | 0.40 ± 0.02<br>(0.87 ± 0.06) |

Table S3: $T_{1\rho}^{H}$ relaxation time values (milliseconds) at sample temperatures from -5° C to 50° C, determined from stretched-exponential fits to the data in Fig. S13 and plotted in Fig. S18. Stretching exponents are given in parentheses. Identical values are listed for the S11 $C_\alpha$ and F19 $C_\alpha$ sites because signals from these sites were not resolved. Uncertainties are standard errors from the fits.

| residue and carbon site | | -5 °C | 0 °C | 5 °C | 10 °C | 20 °C | 30 °C | 40 °C | 50 °C |
|---|---|---|---|---|---|---|---|---|---|
| L6 | β | 23.5 ± 7.4 (0.31 ± 0.06) | 18.0 ± 7.7 (0.52 ± 0.19) | 7.3 ± 4.0 (0.31 ± 0.10) | 13.5 ± 3.6 (0.47 ± 0.10) | 14.2 ± 4.8 (0.36 ± 0.08) | 19.4 ± 3.0 (0.50 ± 0.08) | 9.4 ± 0.7 (0.56 ± 0.04) | 12.1 ± 1.1 (0.59 ± 0.05) |
| | γ | 20.8 ± 12.2 (0.34 ± 0.12) | 13.0 ± 4.9 (0.30 ± 0.06) | 12.0 ± 8.5 (0.31 ± 0.13) | 14.3 ± 5.4 (0.42 ± 0.12) | 11.5 ± 2.6 (0.42 ± 0.07) | 7.4 ± 1.0 (0.46 ± 0.05) | 9.6 ± 1.7 (0.56 ± 0.10) | 14.2 ± 1.4 (0.74 ± 0.09) |
| | $\delta_1$ | 12.4 ± 2.9 (0.46 ± 0.08) | 11.1 ± 2.9 (0.49 ± 0.10) | 11.0 ± 3.1 (0.48 ± 0.10) | 10.2 ± 1.8 (0.50 ± 0.07) | 14.1 ± 1.3 (0.61 ± 0.06) | 12.2 ± 0.9 (0.67 ± 0.06) | 14.7 ± 0.9 (0.69 ± 0.05) | 18.9 ± 1.3 (0.60 ± 0.04) |
| | $\delta_2$ | 24.2 ± 7.9 (0.44 ± 0.10) | 18.2 ± 4.7 (0.42 ± 0.08) | 10.7 ± 3.1 (0.36 ± 0.06) | 10.2 ± 1.1 (0.53 ± 0.05) | 15.2 ± 2.1 (0.54 ± 0.07) | 13.1 ± 0.7 (0.64 ± 0.04) | 14.9 ± 1.2 (0.70 ± 0.07) | 19.7 ± 1.2 (0.62 ± 0.04) |
| S11 | α | 45.9 ± 4.0 (0.48 ± 0.03) | 29.7 ± 5.1 (0.44 ± 0.06) | 29.4 ± 3.5 (0.44 ± 0.04) | 13.8 ± 2.1 (0.50 ± 0.07) | 13.2 ± 1.3 (0.54 ± 0.05) | 15.3 ± 0.7 (0.61 ± 0.03) | 17.2 ± 0.6 (0.65 ± 0.03) | 26.0 ± 0.7 (0.67 ± 0.02) |
| | β | 50.0 ± 4.2 (0.50 ± 0.03) | 38.8 ± 7.5 (0.47 ± 0.07) | 36.6 ± 5.5 (0.40 ± 0.04) | 19.8 ± 2.5 (0.45 ± 0.05) | 16.3 ± 1.1 (0.57 ± 0.04) | 18.3 ± 0.5 (0.60 ± 0.02) | 19.4 ± 0.6 (0.67 ± 0.03) | 25.9 ± 1.4 (0.68 ± 0.04) |
| A16 | α | 34.2 ± 5.0 (0.59 ± 0.08) | 20.9 ± 4.6 (0.54 ± 0.11) | 24.4 ± 4.7 (0.45 ± 0.06) | 14.4 ± 1.7 (0.41 ± 0.04) | 11.7 ± 0.8 (0.61 ± 0.04) | 13.9 ± 1.3 (0.61 ± 0.06) | 18.7 ± 1.3 (0.75 ± 0.07) | 24.8 ± 2.7 (0.64 ± 0.08) |
| | β | 20.1 ± 1.5 (0.47 ± 0.03) | 13.5 ± 1.2 (0.46 ± 0.03) | 20.6 ± 2.1 (0.47 ± 0.04) | 13.9 ± 0.8 (0.50 ± 0.02) | 16.1 ± 1.5 (0.56 ± 0.05) | 22.5 ± 0.9 (0.59 ± 0.03) | 26.3 ± 1.2 (0.71 ± 0.04) | 39.4 ± 3.3 (0.63 ± 0.06) |
| F19 | α | 45.9 ± 4.0 (0.48 ± 0.03) | 29.7 ± 5.1 (0.44 ± 0.06) | 29.4 ± 3.5 (0.44 ± 0.04) | 13.8 ± 2.1 (0.50 ± 0.07) | 13.2 ± 1.3 (0.54 ± 0.05) | 15.3 ± 0.7 (0.61 ± 0.03) | 17.2 ± 0.6 (0.65 ± 0.03) | 26.0 ± 0.7 (0.67 ± 0.02) |
| | β | 27.9 ± 2.0 (0.56 ± 0.03) | 25.7 ± 2.8 (0.66 ± 0.08) | 23.4 ± 1.9 (0.52 ± 0.04) | 14.5 ± 1.7 (0.45 ± 0.04) | 12.4 ± 0.8 (0.57 ± 0.04) | 13.6 ± 0.5 (0.67 ± 0.03) | 17.2 ± 0.5 (0.67 ± 0.02) | 23.3 ± 1.6 (0.69 ± 0.05) |
| | C2-C6 | 21.1 ± 1.4 (0.50 ± 0.03) | 14.1 ± 1.2 (0.59 ± 0.05) | 19.1 ± 1.5 (0.56 ± 0.04) | 13.5 ± 0.8 (0.63 ± 0.04) | 16.1 ± 0.6 (0.70 ± 0.03) | 20.8 ± 0.6 (0.75 ± 0.03) | 31.8 ± 1.2 (0.75 ± 0.04) | 49.3 ± 1.5 (0.71 ± 0.02) |
| T22 | β | 19.2 ± 1.6 (0.53 ± 0.04) | 11.9 ± 1.8 (0.47 ± 0.05) | 21.2 ± 3.0 (0.54 ± 0.06) | 12.5 ± 1.5 (0.54 ± 0.06) | 14.6 ± 1.1 (0.65 ± 0.05) | 19.4 ± 0.6 (0.71 ± 0.03) | 25.6 ± 1.5 (0.77 ± 0.06) | 47.7 ± 3.2 (0.67 ± 0.05) |
| | γ | 16.8 ± 1.2 (0.48 ± 0.03) | 13.8 ± 1.2 (0.57 ± 0.04) | 18.3 ± 1.2 (0.55 ± 0.03) | 16.0 ± 0.9 (0.64 ± 0.04) | 20.5 ± 1.2 (0.64 ± 0.05) | 27.7 ± 0.8 (0.69 ± 0.03) | 41.1 ± 2.4 (0.68 ± 0.04) | 54.6 ± 2.0 (0.69 ± 0.03) |
| G28 | α | 32.1 ± 1.6 (0.63 ± 0.03) | 33.8 ± 4.8 (0.44 ± 0.05) | 23.8 ± 3.0 (0.48 ± 0.05) | 16.9 ± 1.8 (0.51 ± 0.05) | 15.0 ± 1.0 (0.59 ± 0.04) | 13.8 ± 0.5 (0.60 ± 0.02) | 18.7 ± 0.7 (0.70 ± 0.04) | 27.0 ± 1.1 (0.71 ± 0.04) |

Table S4: $T_2^H$ relaxation time values (milliseconds) at sample temperatures from -5° C to 50° C, determined from single-exponential or stretched-exponential fits to the data in Fig. S14 and plotted in Fig. S19. For stretched-exponential fits, stretching exponents are given in parentheses. Identical values are listed for the S11 $C_\alpha$ and F19 $C_\alpha$ sites because signals from these sites were not resolved. Uncertainties are standard errors from the fits.

| residue and carbon site | | -5 °C | 0 °C | 5 °C | 10 °C | 20 °C | 30 °C | 40 °C | 50 °C |
|---|---|---|---|---|---|---|---|---|---|
| L6 | β | 17.6 ± 5.5 | 13.3 ± 2.8 | 14.6 ± 3.6 | 13.8 ± 3.2 | 12.1 ± 2.6 | 12.5 ± 1.4 | 10.6 ± 1.5 | 11.7 ± 1.0 |
| | γ | 16.9 ± 5.8 | 15.1 ± 3.7 | 6.7 ± 1.6 | 9.0 ± 1.1 | 11.9 ± 2.5 | 9.4 ± 1.4 | 10.6 ± 1.5 | 9.2 ± 0.8 |
| | $\delta_1$ | 12.7 ± 2.5 (0.7 ± 0.16) | 11.9 ± 2.7 (0.61 ± 0.17) | 10.9 ± 1.4 (0.78 ± 0.13) | 13.1 ± 1.4 (0.72 ± 0.09) | 18.9 ± 3.2 (0.73 ± 0.17) | 15.1 ± 1.3 (0.73 ± 0.08) | 20.2 ± 1.7 (0.74 ± 0.08) | 19.3 ± 1.7 (0.80 ± 0.09) |
| | $\delta_2$ | 15.7 ± 3.6 (0.68 ± 0.16) | 11.8 ± 2.3 (0.73 ± 0.19) | 16.9 ± 4.3 (0.69 ± 0.21) | 14.6 ± 1.5 (0.93 ± 0.14) | 17.0 ± 3.2 (0.72 ± 0.18) | 17.9 ± 1.4 (0.75 ± 0.09) | 21.0 ± 3.2 (0.66 ± 0.12) | 21.7 ± 2.9 (0.89 ± 0.17) |
| S11 | α | 20.1 ± 1.0 (0.89 ± 0.05) | 15.6 ± 0.9 (0.81 ± 0.07) | 16.3 ± 1.2 (0.79 ± 0.07) | 14.2 ± 0.9 (0.83 ± 0.07) | 13.8 ± 1.1 (0.82 ± 0.09) | 12.7 ± 0.7 (0.82 ± 0.06) | 16.8 ± 1.0 (0.80 ± 0.07) | 17.6 ± 1.0 (0.89 ± 0.06) |
| | β | 19.7 ± 3.4 (0.66 ± 0.12) | 21.1 ± 3.9 (0.53 ± 0.10) | 16.9 ± 2.8 (0.66 ± 0.12) | 15.5 ± 2.4 (0.61 ± 0.10) | 16.7 ± 2.1 (0.60 ± 0.08) | 14.3 ± 1.0 (0.65 ± 0.05) | 18.9 ± 1.3 (0.82 ± 0.08) | 17.5 ± 1.3 (1.05 ± 0.11) |
| A16 | α | 18.7 ± 3.0 (0.7 ± 0.12) | 11.6 ± 1.7 (0.83 ± 0.18) | 13.4 ± 2.0 (0.84 ± 0.17) | 11.8 ± 1.1 (1.08 ± 0.16) | 10.7 ± 1.0 (0.81 ± 0.10) | 11.9 ± 1.2 (0.98 ± 0.16) | 13.9 ± 1.4 (0.75 ± 0.09) | 14.8 ± 1.1 (0.98 ± 0.10) |
| | β | 17.6 ± 1.4 (0.61 ± 0.05) | 12.9 ± 1.3 (0.69 ± 0.09) | 18.0 ± 1.7 (0.66 ± 0.07) | 16.7 ± 1.4 (0.76 ± 0.08) | 19.3 ± 1.4 (0.83 ± 0.09) | 20.4 ± 0.9 (0.82 ± 0.06) | 28.0 ± 2.1 (0.81 ± 0.09) | 29.9 ± 2.7 (1.07 ± 0.17) |
| F19 | α | 20.1 ± 1.0 (0.89 ± 0.05) | 15.6 ± 0.9 (0.81 ± 0.07) | 16.3 ± 1.2 (0.79 ± 0.07) | 14.2 ± 0.9 (0.83 ± 0.07) | 13.8 ± 1.1 (0.82 ± 0.09) | 12.7 ± 0.7 (0.82 ± 0.06) | 16.8 ± 1.0 (0.80 ± 0.07) | 17.6 ± 1.0 (0.89 ± 0.06) |
| | β | 16.1 ± 1.9 (1.14 ± 0.2) | 18.7 ± 4.7 (0.78 ± 0.26) | 12.7 ± 2.1 (0.96 ± 0.23) | 12.9 ± 1.6 (1.22 ± 0.25) | 12.3 ± 1.4 (0.92 ± 0.16) | 11.5 ± 0.8 (0.94 ± 0.10) | 14.5 ± 1.1 (1.03 ± 0.12) | 15.7 ± 1.5 (1.17 ± 0.17) |
| | C2-C6 | 8.9 ± 0.8 (0.82 ± 0.09) | 6.4 ± 0.4 (0.93 ± 0.09) | 7.9 ± 0.7 (0.9 ± 0.11) | 8.2 ± 0.7 (1.00 ± 0.13) | 10.9 ± 1.0 (0.97 ± 0.14) | 11.5 ± 0.7 (1.07 ± 0.11) | 13.9 ± 0.6 (1.05 ± 0.07) | 12.9 ± 0.8 (1.14 ± 0.12) |
| T22 | β | 18.5 ± 3.1 (0.62 ± 0.10) | 12.6 ± 1.3 (0.63 ± 0.08) | 14.6 ± 1.0 (0.68 ± 0.05) | 16.1 ± 1.3 (0.94 ± 0.11) | 19.8 ± 2.6 (0.86 ± 0.18) | 18.3 ± 1.3 (0.86 ± 0.10) | 20.5 ± 0.9 (1.40 ± 0.13) | 23.7 ± 1.3 (1.40 ± 0.15) |
| | γ | 15.2 ± 1.1 (0.69 ± 0.05) | 12.2 ± 0.7 (0.79 ± 0.07) | 17.4 ± 1.6 (0.7 ± 0.08) | 16.6 ± 0.9 (0.76 ± 0.05) | 20.3 ± 1.5 (0.84 ± 0.09) | 25.0 ± 1.9 (0.91 ± 0.10) | 34.0 ± 1.8 (0.85 ± 0.06) | 38.7 ± 1.8 (0.91 ± 0.06) |
| G28 | α | 22.1 ± 4.6 (0.64 ± 0.13) | 12.6 ± 1.9 (0.64 ± 0.12) | 18.9 ± 4.5 (0.54 ± 0.12) | 12.2 ± 1.6 (0.63 ± 0.1) | 13.5 ± 2.3 (0.70 ± 0.15) | 11.7 ± 1.2 (0.77 ± 0.10) | 15.9 ± 1.2 (1.00 ± 0.13) | 18.4 ± 2.0 (1.22 ± 0.22) |

Table S5: $T_1^C$ relaxation time values (seconds) at sample temperatures from -5° C to 50° C, determined from single-exponential fits to the data in Fig. S15 and plotted in Fig. S20. Identical values are listed for the S11 $C_\beta$ and F19 $C_\alpha$ sites because signals from these sites were not resolved. A single set of values is given for backbone CO sites, which also did not have resolved signals. Uncertainties are standard errors from the fits.

| residue and carbon site | | -5 °C | 0 °C | 5 °C | 10 °C | 20 °C | 30 °C | 40 °C | 50 °C |
|---|---|---|---|---|---|---|---|---|---|
| L6 | β | 0.32 ± 0.07 | 0.39 ± 0.06 | 0.42 ± 0.09 | 0.47 ± 0.14 | 0.40 ± 0.04 | 0.32 ± 0.01 | 0.35 ± 0.03 | 0.28 ± 0.03 |
| | γ | 0.54 ± 0.06 | 0.46 ± 0.04 | 0.44 ± 0.05 | 0.50 ± 0.03 | 0.49 ± 0.05 | 0.52 ± 0.03 | 0.59 ± 0.04 | 0.53 ± 0.04 |
| | $\delta_1$ | 0.40 ± 0.02 | 0.40 ± 0.01 | 0.44 ± 0.02 | 0.49 ± 0.03 | 0.46 ± 0.04 | 0.47 ± 0.03 | 0.54 ± 0.05 | 0.60 ± 0.04 |
| | $\delta_2$ | 0.36 ± 0.02 | 0.41 ± 0.02 | 0.44 ± 0.03 | 0.55 ± 0.04 | 0.52 ± 0.02 | 0.50 ± 0.01 | 0.58 ± 0.04 | 0.59 ± 0.03 |
| S11 | α | 0.51 ± 0.03 | 0.53 ± 0.02 | 0.50 ± 0.03 | 0.56 ± 0.03 | 0.49 ± 0.02 | 0.53 ± 0.05 | 0.44 ± 0.02 | 0.48 ± 0.03 |
| | β | 0.36 ± 0.02 | 0.31 ± 0.02 | 0.29 ± 0.02 | 0.33 ± 0.02 | 0.27 ± 0.01 | 0.28 ± 0.02 | 0.31 ± 0.02 | 0.26 ± 0.02 |
| A16 | α | 0.62 ± 0.05 | 0.62 ± 0.04 | 0.50 ± 0.04 | 0.55 ± 0.07 | 0.44 ± 0.03 | 0.50 ± 0.02 | 0.47 ± 0.04 | 0.44 ± 0.04 |
| | β | 0.46 ± 0.03 | 0.44 ± 0.03 | 0.51 ± 0.02 | 0.49 ± 0.04 | 0.51 ± 0.03 | 0.52 ± 0.04 | 0.51 ± 0.05 | 0.53 ± 0.04 |
| F19 | α | 0.51 ± 0.03 | 0.53 ± 0.02 | 0.50 ± 0.03 | 0.56 ± 0.03 | 0.49 ± 0.02 | 0.53 ± 0.05 | 0.44 ± 0.02 | 0.48 ± 0.03 |
| | β | 0.33 ± 0.03 | 0.31 ± 0.03 | 0.33 ± 0.03 | 0.29 ± 0.03 | 0.26 ± 0.01 | 0.26 ± 0.03 | 0.25 ± 0.01 | 0.25 ± 0.02 |
| | C1 | 0.53 ± 0.02 | 0.65 ± 0.05 | 0.55 ± 0.04 | 0.57 ± 0.04 | 0.54 ± 0.02 | 0.58 ± 0.01 | 0.63 ± 0.03 | 0.61 ± 0.05 |
| | C2-C6 | 0.42 ± 0.01 | 0.39 ± 0.01 | 0.38 ± 0.01 | 0.38 ± 0.01 | 0.37 ± 0.01 | 0.37 ± 0.03 | 0.36 ± 0.01 | 0.37 ± 0.01 |
| T22 | β | 0.49 ± 0.04 | 0.49 ± 0.02 | 0.43 ± 0.03 | 0.51 ± 0.04 | 0.43 ± 0.03 | 0.40 ± 0.01 | 0.47 ± 0.04 | 0.44 ± 0.02 |
| | γ | 0.32 ± 0.02 | 0.32 ± 0.02 | 0.32 ± 0.02 | 0.35 ± 0.02 | 0.40 ± 0.02 | 0.41 ± 0.04 | 0.43 ± 0.02 | 0.44 ± 0.02 |
| G28 | α | 0.30 ± 0.02 | 0.29 ± 0.03 | 0.29 ± 0.03 | 0.31 ± 0.02 | 0.28 ± 0.03 | 0.27 ± 0.04 | 0.26 ± 0.01 | 0.25 ± 0.02 |
| CO | | 0.96 ± 0.04 | 0.99 ± 0.06 | 1.05 ± 0.07 | 1.09 ± 0.05 | 1.10 ± 0.05 | 1.19 ± 0.03 | 1.15 ± 0.05 | 1.19 ± 0.04 |

Table S6: $T_2^C$ relaxation time values (milliseconds) at sample temperatures from -5° C to 50° C, determined from single-exponential or stretched-exponential fits to the data in Fig. S16 and plotted in Fig. S21. For stretched-exponential fits, stretching exponents are given in parentheses. Identical values are listed for the S11 $C_\alpha$ and F19 $C_\alpha$ sites because signals from these sites were not resolved. A single set of values is given for backbone CO sites, which also did not have resolved signals. Uncertainties are standard errors from the fits.

| $^{13}C$-$T_2$ (ms) | | -5 °C | 0 °C | 5 °C | 10 °C | 20 °C | 30 °C | 40 °C | 50 °C |
|---|---|---|---|---|---|---|---|---|---|
| L6 | β | 2.1 ± 0.6 (0.49 ± 0.09) | 5.4 ± 1.1 (0.62 ± 0.12) | 4.3 ± 1.3 (0.67 ± 0.21) | 5.3 ± 1.2 (0.76 ± 0.19) | 6.5 ± 0.8 (0.82 ± 0.14) | 7.3 ± 0.8 (0.70 ± 0.10) | 9.2 ± 0.7 (0.71 ± 0.06) | 8.5 ± 0.4 (0.81 ± 0.05) |
| S11 | α | 4.6 ± 1.3 (0.60 ± 0.15) | 8.1 ± 1.0 (0.59 ± 0.07) | 12.1 ± 1.3 (0.62 ± 0.07) | 11.5 ± 0.6 (0.68 ± 0.04) | 14.8 ± 0.7 (0.68 ± 0.04) | 18.3 ± 1.0 (0.88 ± 0.07) | 23.5 ± 1.6 (0.81 ± 0.07) | 28.0 ± 2.2 (0.91 ± 0.10) |
| | β | 3.3 ± 0.7 (0.88 ± 0.26) | 6.8 ± 0.9 (0.71 ± 0.10) | 8.5 ± 0.9 (0.82 ± 0.11) | 16.5 ± 1.7 (0.68 ± 0.08) | 14.3 ± 1.7 (0.71 ± 0.11) | 15.4 ± 1.4 (0.95 ± 0.14) | 18.0 ± 1.3 (1.11 ± 0.14) | 25.1 ± 2.0 (0.88 ± 0.12) |
| A16 | α | 2.6 ± 0.8 (0.57 ± 0.14) | 7.5 ± 1.3 (0.76 ± 0.15) | 7.4 ± 0.7 (0.83 ± 0.11) | 7.2 ± 1.2 (0.74 ± 0.14) | 8.9 ± 0.8 (0.98 ± 0.14) | 13.2 ± 1.3 (1.07 ± 0.16) | 24.5 ± 3.1 (0.84 ± 0.12) | 33.2 ± 5.7 (0.78 ± 0.16) |
| | β | 13.5 ± 0.7 | 9.1 ± 0.5 | 16.2 ± 1.4 | 15.1 ± 1.4 | 14.2 ± 0.8 | 39.4 ± 2.2 | 51.4 ± 3.1 | 53.6 ± 4.5 |
| F19 | α | 4.6 ± 1.3 (0.60 ± 0.15) | 8.1 ± 1 (0.59 ± 0.07) | 12.1 ± 1.3 (0.62 ± 0.07) | 11.5 ± 0.6 (0.68 ± 0.04) | 14.8 ± 0.7 (0.68 ± 0.04) | 18.3 ± 1.0 (0.88 ± 0.07) | 23.5 ± 1.6 (0.81 ± 0.07) | 28 ± 2.2 (0.91 ± 0.10) |
| | β | 2.5 ± 0.6 (0.67 ± 0.15) | 5.6 ± 0.9 (0.65 ± 0.11) | 9.2 ± 1.2 (0.62 ± 0.09) | 6.2 ± 0.5 (0.77 ± 0.08) | 10.8 ± 1.1 (0.84 ± 0.12) | 14.3 ± 1.7 (0.87 ± 0.13) | 19.4 ± 1.1 (0.86 ± 0.06) | 23.6 ± 1.6 (0.95 ± 0.10) |
| | C1 | 2.5 ± 0.2 (0.87 ± 0.09) | 2.5 ± 0.1 (0.92 ± 0.06) | 3.1 ± 0.3 (1.06 ± 0.18) | 3.0 ± 0.3 (1.12 ± 0.17) | 3.4 ± 0.3 (1.18 ± 0.20) | 3.9 ± 0.4 (1.32 ± 0.31) | 3.5 ± 0.2 (1.13 ± 0.14) | 4.3 ± 0.4 (1.36 ± 0.30) |
| | C2-C6 | 2.2 ± 0.1 (1.02 ± 0.06) | 2.6 ± 0.1 (1.23 ± 0.09) | 3.1 ± 0.1 (1.21 ± 0.10) | 3.3 ± 0.1 (1.35 ± 0.11) | 4.3 ± 0.1 (1.54 ± 0.12) | 4.7 ± 0.2 (1.87 ± 0.19) | 4.7 ± 0.1 (1.81 ± 0.13) | 5.4 ± 0.1 (1.95 ± 0.10) |
| T22 | β | 14.1 ± 1.8 (0.66 ± 0.09) | 10.2 ± 1.3 (0.66 ± 0.09) | 13.4 ± 1.5 (0.64 ± 0.07) | 15.2 ± 1.3 (0.87 ± 0.10) | 17.4 ± 1.3 (1.05 ± 0.12) | 28.4 ± 3.6 (0.97 ± 0.16) | 36.7 ± 8.7 (0.65 ± 0.13) | 51.9 ± 6.2 (0.82 ± 0.12) |
| | γ | 19.5 ± 2.0 (0.67 ± 0.08) | 19.5 ± 1.3 (0.66 ± 0.06) | 23.5 ± 2.8 (0.82 ± 0.11) | 16.6 ± 0.6 (1.33 ± 0.08) | 16.3 ± 0.9 (1.14 ± 0.10) | 54.8 ± 4.9 (0.93 ± 0.12) | 56.9 ± 2.3 (1.21 ± 0.08) | 56.9 ± 2.4 (1.26 ± 0.10) |
| G28 | α | 3.0 ± 0.8 (0.61 ± 0.15) | 5.6 ± 0.7 (0.58 ± 0.07) | 9.3 ± 1.1 (0.60 ± 0.07) | 7.0 ± 0.5 (0.80 ± 0.07) | 10.4 ± 0.7 (0.77 ± 0.08) | 18.2 ± 1.8 (0.75 ± 0.08) | 22.6 ± 1.1 (0.78 ± 0.04) | 42.5 ± 3.3 (0.70 ± 0.06) |
| CO | | 4.9 ± 0.5 | 5.4 ± 0.3 | 5.6 ± 0.4 | 8.4 ± 0.6 | 13.4 ± 1 | 19.9 ± 0.9 | 21.0 ± 0.9 | 33.0 ± 3.2 |

Table S7: Results from fitting $^{13}C$ $T_1$ and $T_2$ data for $^{13}C$-$^{1}H$ sites in MBPA-C-FG30-LSAFTG/AAO20 with a spectral density of the form $a_1\tau_{c1}/(1+\omega^2\tau_{c1}{}^2)+(1-a_1)\tau_{c2}/(1+\omega^2\tau_{c2}{}^2)$, with $\tau_{c1}$ constrained to the 10-500 ns range and $\tau_{c2}$ constrained to the 0.1-9 ns range. Values of the three fitting parameters are probability-weighted averages, calculated as described in the text, with standard deviations in parentheses.

| site | temperature (° C) | $\tau_{c1}$ (ns) | $\tau_{c2}$ (ns) | $a_1$ |
|---|---|---|---|---|
| S11/F19 $C_\alpha$ | -5 | 161 (95) | 1.00 (0.89) | 0.38 (0.13) |
| | 0 | 124 (104) | 1.11 (1.08) | 0.35 (0.16) |
| | 5 | 125 (118) | 1.17 (1.16) | 0.29 (0.17) |
| | 10 | 134 (122) | 1.35 (1.40) | 0.29 (0.19) |
| | 20 | 127 (122) | 1.20 (1.20) | 0.25 (0.17) |
| | 30 | 121 (123) | 1.39 (1.42) | 0.24 (0.19) |
| | 40 | 110 (121) | 1.13 (1.06) | 0.22 (0.16) |
| | 50 | 100 (120) | 1.26 (1.24) | 0.23 (0.19) |
| A16 $C_\alpha$ | -5 | 206 (90) | 1.04 (0.98) | 0.47 (0.13) |
| | 0 | 112 (95) | 1.29 (1.32) | 0.41 (0.18) |
| | 5 | 164 (118) | 1.13 (1.11) | 0.29 (0.16) |
| | 10 | 152 (116) | 1.24 (1.27) | 0.33 (0.17) |
| | 20 | 163 (119) | 1.03 (0.95) | 0.25 (0.14) |
| | 30 | 112 (113) | 1.15 (1.12) | 0.30 (0.18) |
| | 40 | 103 (120) | 1.21 (1.19) | 0.24 (0.18) |
| | 50 | 92 (117) | 1.17 (1.10) | 0.22 (0.18) |
| T22 $C_\beta$ | -5 | 123 (120) | 1.18 (1.17) | 0.27 (0.17) |
| | 0 | 119 (109) | 1.08 (1.02) | 0.31 (0.16) |
| | 5 | 137 (121) | 1.05 (0.97) | 0.23 (0.14) |
| | 10 | 124 (122) | 1.27 (1.28) | 0.26 (0.18) |
| | 20 | 130 (123) | 1.08 (1.02) | 0.21 (0.15) |
| | 30 | 80 (105) | 1.00 (0.83) | 0.23 (0.14) |
| | 40 | 81 (113) | 1.26 (1.23) | 0.24 (0.19) |
| | 50 | 100 (122) | 1.32 (1.30) | 0.12 (0.12) |

Table S8: Results from fitting [1]H $T_1$, $T_2$, and $T_{1\rho}$ data for methylene sites in MBPA-C-FG30-LSAFTG/AAO20 with a spectral density of the form $a_1\tau_{c1}/(1+\omega^2\tau_{c1}{}^2)+(1-a_1)\tau_{c2}/(1+\omega^2\tau_{c2}{}^2)$, with $\tau_{c1}$ constrained to the 10-500 ns range and $\tau_{c2}$ constrained to the 0.1-9 ns range. Values of the three fitting parameters are probability-weighted averages, calculated as described in the text, with standard deviations in parentheses.

| site | temperature (° C) | $\tau_{c1}$ (ns) | $\tau_{c2}$ (ns) | $a_1$ |
|---|---|---|---|---|
| L6 $C_\beta$ | -5 | 98 (122) | 0.61 (0.22) | 0.13 (0.13) |
| | 0 | 97 (122) | 0.73 (0.32) | 0.20 (0.18) |
| | 5 | 86 (116) | 0.47 (0.23) | 0.20 (0.16) |
| | 10 | 93 (120) | 0.72 (0.30) | 0.22 (0.20) |
| | 20 | 87 (117) | 0.57 (0.28) | 0.24 (0.19) |
| | 30 | 62 (103) | 0.49 (0.28) | 0.29 (0.19) |
| | 40 | 96 (119) | 0.50 (0.24) | 0.23 (0.18) |
| | 50 | 90 (117) | 0.51 (0.24) | 0.23 (0.18) |
| S11 $C_\beta$ | -5 | 54 (91) | 0.45 (0.18) | 0.15 (0.12) |
| | 0 | 38 (65) | 0.34 (0.17) | 0.22 (0.12) |
| | 5 | 99 (122) | 0.58 (0.14) | 0.07 (0.07) |
| | 10 | 85 (116) | 0.52 (0.22) | 0.18 (0.15) |
| | 20 | 83 (115) | 0.47 (0.21) | 0.20 (0.16) |
| | 30 | 97 (123) | 0.58 (0.22) | 0.15 (0.14) |
| | 40 | 109 (125) | 0.61 (0.17) | 0.11 (0.12) |
| | 50 | 73 (109) | 0.60 (0.16) | 0.14 (0.11) |
| F19 $C_\beta$ | -5 | 113 (127) | 0.90 (0.13) | 0.05 (0.07) |
| | 0 | 69 (105) | 0.64 (0.28) | 0.21 (0.20) |
| | 5 | 104 (124) | 0.67 (0.14) | 0.10 (0.10) |
| | 10 | 66 (105) | 0.46 (0.25) | 0.28 (0.19) |
| | 20 | 77 (112) | 0.50 (0.26) | 0.27 (0.20) |
| | 30 | 86 (117) | 0.54 (0.26) | 0.24 (0.19) |
| | 40 | 86 (118) | 0.49 (0.22) | 0.19 (0.16) |
| | 50 | 41 (82) | 0.40 (0.22) | 0.26 (0.13) |
| G28 $C_\alpha$ | -5 | 123 (129) | 0.87 (0.07) | 0.03 (0.04) |
| | 0 | 108 (125) | 0.76 (0.09) | 0.06 (0.06) |
| | 5 | 100 (123) | 0.54 (0.18) | 0.11 (0.10) |
| | 10 | 69 (107) | 0.46 (0.25) | 0.26 (0.18) |
| | 20 | 85 (117) | 0.51 (0.24) | 0.22 (0.17) |
| | 30 | 85 (116) | 0.49 (0.24) | 0.23 (0.18) |
| | 40 | 92 (120) | 0.52 (0.21) | 0.16 (0.14) |
| | 50 | 119 (128) | 0.69 (0.07) | 0.06 (0.06) |
